# Fatty Acid β-oxidation and Ferroptosis Define a Survival-Productivity Trade Off in CHO fed-Batch Bioreactors

**DOI:** 10.64898/2026.08.20.745185

**Authors:** Charles Eldrid, John Raven, Robyn Hoare, Sam Whitwam, Alan Dickson, Andrew Pitt, Leon Pybus, Perdita Barran

## Abstract

Fed-batch production in Chinese hamster ovary cells is usually optimised empirically, yet the intracellular mechanisms that determine whether cells sustain productivity or enter terminal decline remain poorly resolved. Here, we combine longitudinal proteomics and intracellular metabolomics across differential ambr250 fed-batch processes to define the metabolic programmes associated with CHO cell viability and antibody production. Across media/feed combinations and an intensified seeding regime, culture progression followed a conserved trajectory from proliferation to metabolic transition and terminal stress. The high-stress state was characterised by a switch toward mitochondrial and peroxisomal fatty acid β-oxidation, lipid remodelling, oxidative burden and activation of ferroptosis-associated pathways. Enriched feed conditions delayed this transition through enhanced redox and glutathione-linked defence, but did not proportionally increase antibody titre, revealing a trade-off in which cellular resources are diverted from recombinant protein production toward survival. These data identify fatty acid metabolism and ferroptosis as key constraints on late-stage CHO fed-batch performance and provide a mechanistic framework for rational feed design and host-cell engineering.

## Introduction

Chinese hamster ovary (CHO) are the predominant host system for the manufacturing of recombinant biotherapeutics in the global biopharmaceutical market.^1,2^ They are highly prized for their productivity, resilience, high growth rates under bioreactor conditions and capacity for human-like post-translational modifications (PTMs). Improvement of CHO productivity and efficiency will enable biotherapeutics to be manufactured at reduced costs and lower energy expenditure.

The current strategies for improving CHO productivity involve empirical screening of diverse media formulations, nutrient feeds, and operational process parameters^3,45^ Historically, the monitoring and control of bioreactors have relied on gross physiological parameters such as viable cell count (VCC) and dissolved oxygen (dO_2_).^6^ These macroscopic readouts cannot elucidate the complex intracellular metabolic processes that underpin improved biomass accumulation and high product titres. Further performance gains can be achieved through cell line engineering, but this remains a laborious process, that requires a deep understanding of the complex interactions occurring within the cellular chassis under the strain of antibody production within the changing environment of a bioreactor. ^7–11^

Quantitative ‘omics of CHO cell bioreactors provides biological insight into cellular states at a molecular level.^12,13^ Such an approach enables comprehensive identification and quantification of diverse metabolites and proteins driving cellular phenotypes.^14–16^ Our motivation is to enable bioprocess engineers to make more informed decisions regarding bioreactor feeds and operating conditions to reduce reliance on brute force empirical parameterisation.^16^ Our prior investigation of a CHO DG44 monoclonal antibody (mAb) expressing cell line under fed-batch bioreactor conditions, provided proteomic evidence for three distinct phases: proliferation, production and stress.^13^ We identified a clear temporal shift from an initial growth stage (days 0-3), to a stationary phase where the majority of the product is produced (days 4-9), culminating in a terminal stress phase where productivity and cellular viability decline (days 10-14).^13^

Here, we ask why increased viable biomass does not necessarily translate into increased antibody production in fed-batch CHO culture. By comparing standard, enriched and intensified fed-batch processes using longitudinal proteomics and intracellular metabolomics, we show that late-stage culture decline^13^ is governed by a conserved metabolic transition into fatty acid oxidation, oxidative lipid stress and ferroptosis^17^. This identifies a survival–productivity trade-off that limits process performance and provides a mechanistic framework for rational feed and host-cell engineering.

## Methods

### Media and feed combinations

Samples that were subjected to an intensified fed-batch (IFB) process were evaluated within ambr250 high throughput bioreactor systems (Table 1).

**Table 1.** Media/Feed Conditions. Two basal media were explored: HNP-006 (H6) and JM-05B (J5), which were combined with two different feeds: BSEL and Cytiva, to create 4 combinations: HNP-006 BSEL (H6-B), HNP-006 Cytiva (H6-C), JM-05B BSEL (J5-B) and JM-05B Cytiva (J5-C). H6 medium is an enriched form of J5, with approximately 35% higher concentration of starting compounds. The BSEL feed contains differing concentrations of amino acids compared to Cytiva: higher Arg, Asn, His and Lys, and lower Asp. J5-C represents an entirely unenriched media and feed strategy and is designated as the “control” baseline for this study.

| **Functional label** | **Code** | **Rationale** |
| --- | --- | --- |
| **Control baseline** | J5-C | unenriched medium/feed |
| **High viability** | H6-B | enriched condition with highest late-stage VCD |
| **High stress** | IFB | intensified seeding, accelerated decline |
| **Comparator conditions** | H6-C, J5-B | distinguish medium/feed effects |

#### ambr250 Bioreactor Operation

Fed-batch production was performed in stirred ambr250 bioreactors (Sartorius) using the specified chemically defined production media. Cells were inoculated at seeding densities between 0.5 and 10 × 10^6^ cells/mL, as indicated.

Nutrient feed was added once glucose fell below 4.0 g/L and the feed quantity was adjusted daily based on glucose measurements and continued until day 13 of culture, with maximum daily addition of 3 % (v/v). In addition, L-glutamine supplementation was performed by adding 0.5 % (v/v) of a 200 mM solution daily from day 4 to day 13. Glucose levels were maintained by a bolus addition of a 500 g/L glucose solution when levels dropped below 4.0 g/L. Bioreactors were operated under controlled conditions at 36 °C, 40 % dO_2_, and pH 6.9.

Specific productivity (Qp) was calculated using the following equation:

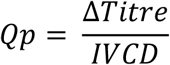

Where IVCD (integrated viable cell density) is:

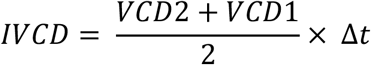

Where VCD is the viable cell density at timepoint t.

#### Proteomics Sample Preparation

Cell pellets (1 x 10^7^ cells) were prepared according to the S-trap 96 well plate protocol.^18^ Briefly, cell pellets were kept on ice before lysing with 50 µL 10 % (w/v) SDS lysis solution, and sonicated for 10 minutes. To remove DNA, 2 μL Benzonase (Cambridge Bioscience Limited) were added, and the samples were shaken in a ThermoMixer (Starlab, UK). Samples were clarified by centrifugation for 8 minutes at 13,000 g. 10 µL of lysate, equivalent to approximately 30 µg of protein based on Bradford assay calculations, was taken for reduction and alkylation. 2 µL of 120 mM TCEP was added and the samples were incubated at 55 °C for 15 minutes, then allowed to cool. 2 µL of 500 mM IAA was added and the samples were incubated at room temperature in the dark for an hour. 5 µL of 27.5 % phosphoric acid to make a concentration of 2.5 % was added, and samples were vortexed. 350 µL of the binding wash buffer was added, the samples were then mixed and the sample was transferred to the S-trap plate. 200 µL of the wash buffer was applied 3 times, using a vacuum manifold to pull the solution through the plate. 3 µg of trypsin gold was applied to each sample, and the plates were incubated at 47 °C for 2 hours, before samples were eluted using 80 µL of elution buffer 1 (50 mM TEAB), 40 µL of elution buffer 2 (0.2 % formic acid v:v), and 40 µL of elution buffer 3 (50 % acetonitrile v:v), pulling the solution into 96 well plates using the vacuum manifold each time. The samples were dried down using the Genevac miVac centrifugal evaporator (Scientific Products, UK). Samples were resuspended in 0.1 % formic acid to a concentration of 40 ng/µL.

#### LC-MS

Samples were randomised, and 200 ng of peptides were directly injected onto a nanoEase M/Z Peptide CSH C18 Column, 130 Å, 1.7 μm, 300 μm × 150 mm (Waters Corp) on an Acquity UPLC M-class system. ZenoTOF 7600+ data were collected in same manner as previously^13^: positive data independent acquisition (DIA) mode using a 50-min gradient at 2 μl/min (**Table S1**), with an OptiFlow 50 to 200 μl Micro/MicroCal source. Data were collected using 105 variable windows from 450 to 2050 m/z, and autocalibration was performed every five injections. For data independent acquisition (DIA) mode instrument parameters see **Table S2**.

#### Data Analysis

Data were analysed using Spectronaut v18.4 (Biognosys)^19^ using BGS factory settings in a peptide-centric fashion. Peptides were identified using a dynamic mass tolerance, with a 1 % false discovery rate based on mutated decoy sequences. Q-value of the precursor and the protein cut-off were set to 0.05, quantification was based on MS2 area, with the normalisation strategy set to “automatic”, which for under 500 runs is a MaxLFQ strategy based on inter-run peptide ratios. The data was searched initially with the spectral library created from previous work by Eldrid *et al.,*^13^ and then was searched with mammalian sequences from UniProt, with the additional mAb construct heavy and light chain sequences. Principal component analysis (PCA) was used to identify outlier runs, which were then manually inspected, and removed based on number of precursors, total ion chromatogram signal, or deviation from the average, and the data were reanalysed. Volcano plots were created directly from Spectronaut candidate tables using Log_2_ ratio and -Log_10_ Qvalue (a confidence modified p-value score) cutoffs as significance values. Correlation clustering analysis was performed in python 3.7 using the pandas, sklearn and seaborn packages. Data were correlated with a Pearson coefficient, and clustered using ward linkages and Euclidean distance metric. Gene names and Kyoto Encyclopaedia of Genes and Genomes (KEGG) identifiers were mapped to UniProt accessions using the UniProt ID mapping tool (https://www.UniProt.org/id-mapping). Gene set enrichment analysis was performed using the gseapy python package (https://github.com/zqfang/GSEApy)^20^ using the gene ontology (GO) Biological Processes 2025 library for *Mus musculus*.^21,22^ Mapping of pathways to volcano plots was performed using gene sets download from Harmonizome 3.0.^23,24^ KEGG mapping was performed using the KEGG mapping tool (https://www.genome.jp/kegg/mapper/color.html), where the mapped to unique KEGG orthology (KO) numbers from differentially regulated candidates identified from the volcano plots to the *Cricetulus griseus* KEGG metabolic map with aliases.^25–27^

#### Metabolomics Data Acquisition and Analysis

Data was acquired using the EdinOmics in-house rapid HILIC-Z Ion Mobility Mass Spectrometry (RHIMMS) method.^28^ Data was acquired in positive and negative ionisation modes on an Agilent 1290 Infinity II series UHPLC system hyphenated with an Agilent 6560 ion mobility quadrupole time-of-flight instrument. Experimental samples were blocked and randomised to minimise the impact of measurement error. Pooled quality control samples obtained from the experimental samples were collected after every 5 experimental sample injections to monitor the instrument state throughout data acquisition. The raw data was processed using the Agilent MassHunter software suite and underwent: demultiplexing (PNNL PreProcessor v2020.03.23), accurate mass calibration (AgtTofReprocessUI 10.0), drift time calibration (IM-MS Browser 10.0), feature extraction (Mass Profiler 10.0), high resolution demultiplexing (HRdm v41), final feature extraction (Mass Profiler 10.0), annotation using accurate mass and CCS values against selected personal compound databases (PCDLs) and libraries. Where multiple features were assigned the same metabolite annotation, one of each annotation was selected based on: the frequency of samples it was present in, the availability of collision cross section (CCS) data, its average intensity and its ion type (preferring [M+H]+, [M-H]-, or [M+Na]+ ions). Features with variable importance point scores which could not be matched to the library using accurate mass and CCS, were manually annotated using an HMDB^29^ search of 5 ppm mass tolerance and 3 % DeepCCS^30^ tolerance.

To increase the reliability of statistical analysis, the data underwent a Log_10_ transformation as well as mean-centre and Pareto (dividing the metabolite intensities for each sample by the square root for their standard deviation) scaling to give the frequency of metabolite intensities a normal/Gaussian distribution.

## Results & Discussion

All proteins and genes referred to in the text are referenced in Table 3.

### Media and Seeding Density Impact Viability and Productivity

We first asked whether media enrichment or process intensification altered the relationship between biomass, viability and antibody production. All cultures (**Figure 1**) demonstrated a conserved macroscopic growth trajectory consistent with our previously defined proteomic work: cellular growth with high levels of growth and DNA replication biomarkers (days 0 – 3); stationary phase where the cells have the highest productivity and high viability (days 4 – 9); and cellular stress and death phase where the cellular viability drops and there are high levels of biomarkers indicative of oxidative stress, inflammation and lysosomal activity (days 10 – 14) (**Figure 1**A-C).^13^ H6-B shows the highest level of growth amongst all tested bioreactor conditions and retains a higher viable cell density (VCD) than the other conditions during the stress and decline phase. Due to this we class this condition as a “high viability state”. Interestingly, although the H6-B and H6-C strategies enable higher peak biomass densities, this extended longevity did not yield a corresponding increase in product titre, even though productivity is maintained (**Figure 1**C-D). This mismatch indicates that the metabolic resources provided by the enriched medium were not effectively translated into increased mAb synthesis and instead were used to support cellular viability. High levels of cellular proliferation were observed from days 0 – 8, with growth plateauing at day 10 (**Figure 1**A), and an accompanying drop in cellular viability from days 10 – 14 (**Figure 1**B). While cultures with J5 basal media tracked closely with the control baseline, the enriched H6 cultures demonstrated superior viable cell accumulation between days 8 – 10, sustaining higher viable cell density (VCD) during the terminal stages (days 10 – 14).

**Figure 1:**
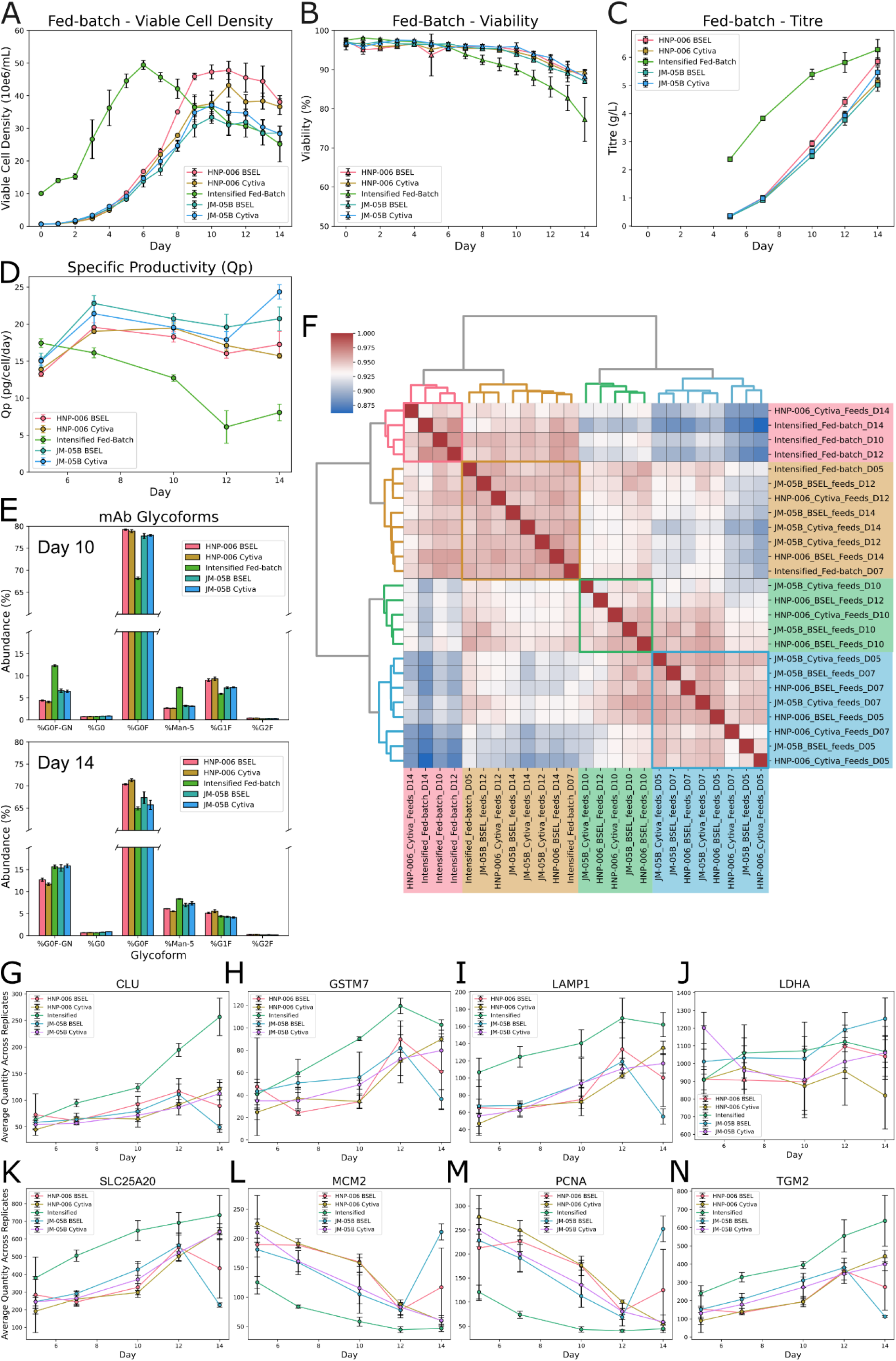
Collected Biostat data from ambr250 fed-batch bioreactors for HNP-006 BSEL, HNP-006 Cytiva, Intensified Fed-batch, JM-05B BSEL and JM-05B Cytiva **A)** viable cell density **B)** viability **C)** product titre and **D)** calculated specific productivity (Qp). Proteomics results **E)** relative glycoform abundance of the harvested product after 10 and 14 days, **F)** Correlation clustering of the bioreactors based on the averaged quantities of all identified protein groups. Average quantity of biomarkers previously identified in fed-batch bioreactors by Eldrid *et al.,* as being identifying particular bioreactor states: **G)** CLU **H)** GSTM7 **I)** LAMP1 **J)** LDHA **K)** SLC25A20 **L)** MCM2 **M)** PCNA and **N)** TGM2

The intensified fed-batch (IFB), seeded at a higher density, shows a temporally shifted growth curve. Inoculated at an initial VCD of 1 x 10^7^ cells on day 0 (a baseline equivalent to day 5 in standard operations), the IFB reached a peak VCD of 5 x 10^7^ cells by day 6, matching the maximum VCDs achieved with the H6-B condition. Following this peak, the IFB culture demonstrated a severe viability decline that was far more pronounced than standard operation. The IFB has consistently lower cell specific productivity (Qp) compared to the other cultures from day 7 (**Figure 1**D). Due to the increased cellular density in earlier days, the titre attains similar levels to that of H6-B at 6 g/L. The shifted growth curve and more intense loss in viability suggest that the IFB has an extended stress phase identified in our earlier publication.^13^

The drop in productivity in IFB is also matched by a drop in product quality: there are higher levels of high mannose (Man-5) or the monoantennary G0F-GlcNAc at both day 10 and 14 which is indicative of immature glycan processing or partial degradation (**Figure 1**E). HNP-006 media bioreactors show the lowest levels of these glycans, however all bioreactors show an increase between day 10 and 14, suggesting that this “stress” phase impacts product glycosylation. Product consistency is key for manufacturing control of biotherapeutics, and variations in glycan can lead to reduced activation of Fcγ receptors, increased product clearance and possible immunogenic effects.^31,32^ The drop in product quality suggests that the mechanisms which drive the production of the low viability state also impact mAb production, glycosylation and degradation.

The differential growth patterns, productivity and product quality of the H6-B and IFB bioreactors suggest that they will provide the most insights to the molecular pathways underlying growth and viability (see below). The differences between enriched media and high density seeding suggest that that control of media supply and consumption is key to improving the performance of CHO cells within bioreactors.

### All Bioreactors Show Consistent Proteome Progression

Proteomics analysis provides details on the metabolic transitions which define the different phases of cellular viability and productivity. On average 4199 ± 125 protein groups were identified across all the samples, with a highly conserved core of 2712 proteins consistently quantified across all conditions. Hierarchical cluster analysis of the proteomics data show that the bioreactors clearly segregated into three distinct temporal clusters: days 5 – 7 (proliferation), day 10 (metabolic transition), and days 12 – 14 (terminal stress) (**Figure 1**F).^13^ Interestingly, the early-stage IFB proteome (day 5 – 7) clustered with the late-stage profiles (days 12 – 14) of the standard bioreactors, while the late-stage IFB samples (days 10 – 14) formed an independent branch alongside the terminal H6-C proteome (day 14). This correlation underscores that the higher seeding density accelerates the onset of the cellular decline, effectively modelling the terminal biology of extended fed-batch operations, giving insight into the cellular processes which underpin loss of productivity and viability.

To track phase transitions, we evaluated a panel of eight established CHO bioprocess biomarkers: clusterin (CLU)^33,34^, glutathione-S-transferase Mu 7 (GSTM7)^35,36^, lysosomal associated membrane protein 1 (LAMP1)^37^, lactose dehydrogenase A (LDHA)^38^, mitochondrial carnitine/acylcarnitine transferase (SLC25A20)^39,40^, DNA licensing factor MCM2 (MCM2)^41,42^, proliferating cellular nuclear antigen (PCNA)^43^ and protein-glutamine gamma-glutamyl-transferase-2 (TGM2)^44–46^. These biomarker proteins demonstrated robust, highly reproducible trends: (**Figure 1**G-N): proliferation drivers (MCM2, PCNA) systematically decrease over time, while lysosomal, apoptotic and stress-responsive markers (GSTM7, LAMP, TGM2) rose sharply during late-stage culture. The chaperone CLUS, and the fatty acid transporter SLC25A20 both increased during the terminal phase, reinforcing the validity of these markers across diverse operational contexts.^13^

The longitudinal variation of these biomarkers followed similar trends irrespective of bioprocess strategies, indicating that baseline phase progression remains tightly constrained. The J5-B condition demonstrated an apparent late-stage abundance increase in cellular proliferation markers (MCM2, PCNA) and a decrease in apoptotic markers (LAMP1, TGM2) on day 14 (**Figure 1**I, L-N), this molecular response was insufficient to rescue culture VCD or viability (**Figure 1**G-H). Conversely, the IFB condition elicited the most extreme shifts in these stress biomarkers, matching its accelerated drop in culture viability. Due to the lower viability and higher levels of stress phase biomarkers, we designate the IFB as a “high stress state”.

Longitudinal comparison of differentially expressed protein across each bioreactor culture against the control identified key regulatory networks governing CHO cell growth rates and viability. To resolve how these dynamic proteomic shifts drive distinct cellular phenotypes, differentially abundant proteins were mapped to cellular pathways via gene set enrichment analysis (GSEA). Volcano distributions were characterised for each operational condition over time, with significantly altered candidates sequentially mapped onto global KEGG metabolic networks (**Figures S1-30**).

Broadly, enriched media and feed combinations demonstrated a systematic upregulation of energy production pathways and amino acid catabolism relative to the control (**Figures S1-10, S15-25**), consistent with higher Arg, Asn, His and Lys.^4^ Due to their highly divergent physiological outcomes, the H6-B (high viability) and IFB (high stress) cultures were selected for deep mechanistic review (**Figure 1**A).

### Switch to Fatty Acid Oxidation Promotes Ferroptosis

The high seeding density in IFB created a “high stress” microenvironment which yielded deep insights into the molecular networks governing cellular decline, demonstrating differential enrichment in fatty acid metabolism, energy production and ferroptosis relative to the control, where DNA replication is highly upregulated, suggesting that there is significant downregulation of growth pathways (Figures S16-20). During the late-stage transition (days 10 – 12), there is significant enrichment of fatty acid metabolism pathways, highlighting a shift towards fatty acid β-oxidation (FAO) as an energy source (**Figure 2**A, D-E, S16-20, S31, **Table 2**).

**Figure 2:**
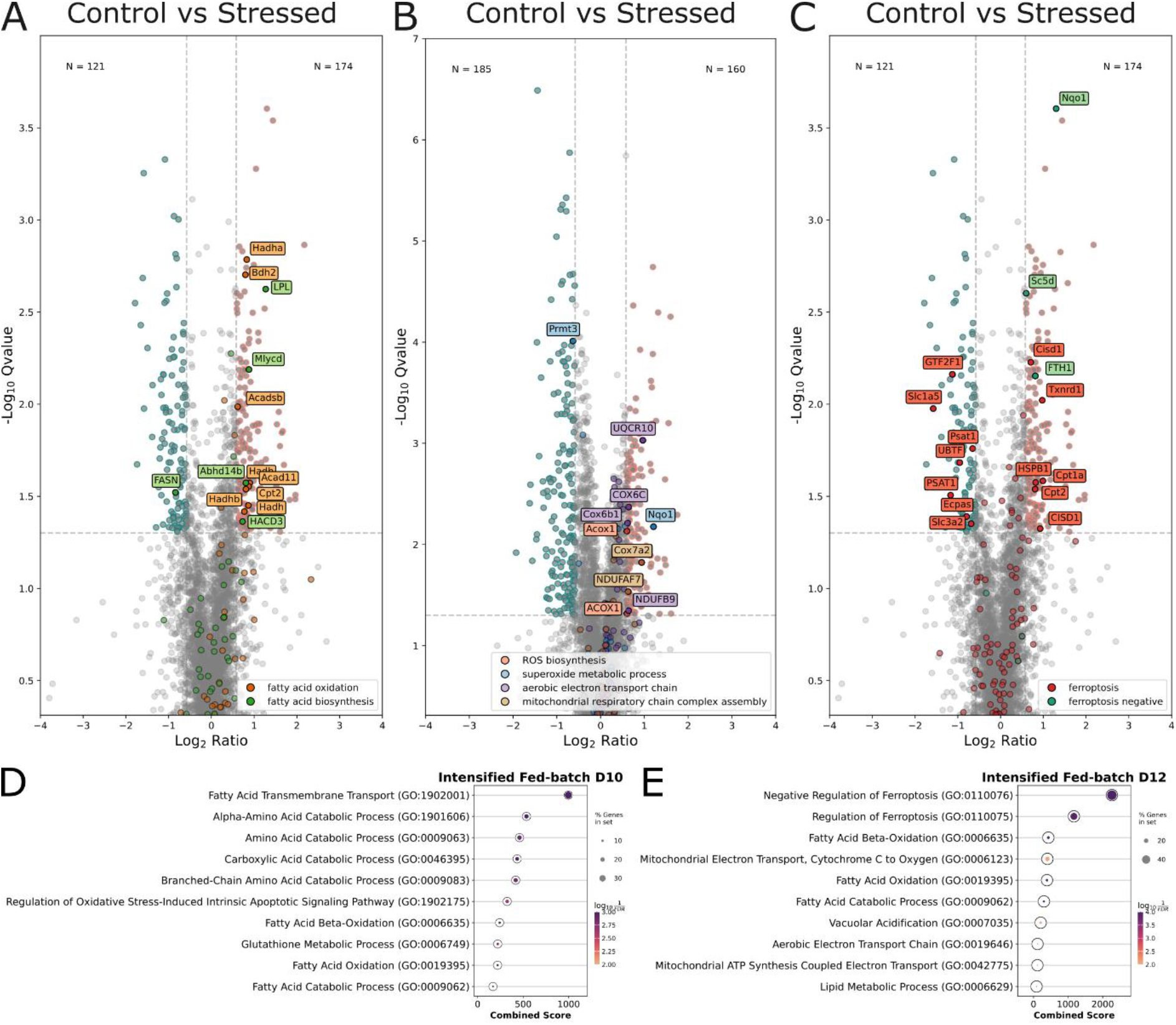
Differentially abundant hits between the control (blue) and high stress (pink) phenotypes with GO biological processes pathways marked **A)** fatty acid biosynthesis (green) and fatty acid oxidation (orange), day 10 **B)** ROS biosynthesis (red), superoxide metabolic process (blue), aerobic electron transport chain (purple) and mitochondrial respiratory chain complex (tan), day 12 **C)** positive (red) and negative (green) ferroptotic control, day 10 **D)** gene set enrichment for IFB on day 10 and **E)** day 12.

**Table 2:** enriched pathways found in the stressed phenotype (IFB) compared to the control (J5-C), showing upregulation of a variety of pathways involved in fatty acid utilisation and amino acid breakdown.

| GO BIOLOGICAL PROCESS TERM | ADJUSTED P-VALUE | ODDS RATIO | COMBINE D SCORE | GENES |
| --- | --- | --- | --- | --- |
| FATTY ACID TRANSMEMBRANE TRANSPORT | 4.81E-04 | 7.31E+01 | 9.96E+02 | CPT1A;CPT2;CD36;SLC25A20 |
| ALPHA-AMINO ACID CATABOLIC PROCESS | 4.81E-04 | 3.79E+01 | 5.36E+02 | ALDH6A1;HIBADH;PIPOX;PRODH;ACADSB |
| AMINO ACID CATABOLIC PROCESS | 1.25E-03 | 3.94E+01 | 4.60E+02 | BCKDHA;ALDH6A1;ALDH5A1;MCCC1 |
| CARBOXYLIC ACID CATABOLIC PROCESS | 4.81E-04 | 3.22E+01 | 4.34E+02 | BCKDHA;MPST;ALDH6A1;AMDHD2;MCCC1 |
| BRANCHED-CHAIN AMINO ACID CATABOLIC PROCESS | 1.39E-03 | 3.65E+01 | 4.18E+02 | BCKDHA;ALDH6A1;HIBADH;MCCC1 |
| REGULATION OF OXIDATIVE STRESS-INDUCED INTRINSIC APOPTOTIC SIGNALLING PATHWAY | 2.15E-03 | 3.01E+01 | 3.25E+02 | HSPB1;HTRA2;SOD2;PRODH |
| FATTY ACID BETA-OXIDATION | 5.54E-04 | 1.85E+01 | 2.41E+02 | HADHB;HADHA;BDH2;CPT2;ACAD11;HADH |
| GLUTATHIONE METABOLIC PROCESS | 1.39E-03 | 1.95E+01 | 2.21E+02 | GSTK1;GSTM2;GSTA5;GSTA3;GSTP1 |
| FATTY ACID OXIDATION | 6.38E-04 | 1.73E+01 | 2.19E+02 | HADHB;HADHA;BDH2;CPT2;ACAD11;HADH |
| FATTY ACID CATABOLIC PROCESS | 1.25E-03 | 1.44E+01 | 1.69E+02 | HADHB;HADHA;BDH2;CPT2;ACAD11;HADH |

**Table 3.** Gene names, UniProt IDs and protein descriptions for the representative sequence of the protein group referred to in the text.

| Genes | UniProt IDs | Protein Descriptions | Genes | UniProt IDs | Protein Descriptions |
| --- | --- | --- | --- | --- | --- |
| **ACAD11** | Q709F0 | Acyl-CoA dehydrogenase family member 11 | **HADHB** | P31937 | Trifunctional enzyme subunit beta, mitochondria |
| **ACADSB** | P45954 | Short/branched chain specific acyl-CoA dehydrogenase | **HIBADH** | P04792 | 3-hydroxyisobutyrate dehydrogenase, mitochondrial |
| **ACOX1** | Q15067 | Peroxisomal acyl-coenzyme A oxidase 1 | **HSPB1** | O43464 | Heat shock protein beta-1 |
| **Aldh5a1** | P51649 | Succinate-semialdehyde dehydrogenase, mitochondrial | **HTRA2** | P11279 | Serine protease HTRA2, mitochondrial |
| **ALDH6A1** | Q02252 | Methylmalonate-semialdehyde dehydrogenase [acylating], mitochondrial | **LAMP1** | P00338 | Lysosome-associated membrane glycoprotein 1 |
| **AMDHD2** | Q9Y303 | N-acetylglucosamine-6-phosphate deacetylase | **LDHA** | P06858 | L-lactate dehydrogenase A chain |
| **BCKDHA** | P12694 | 2-oxoisovalerate dehydrogenase subunit alpha, mitochondrial | **LPCAT3** | Q5FVN0 | Lysophospholipid acyltransferase 5 |
| **BDH2** | Q9BUT1 | Dehydrogenase/reductase SDR family member 6 | **LPL** | Q96RQ3 | Lipoprotein lipase |
| **CD36** | P16671 | Platelet glycoprotein 4 | **MCCC1** | P33993 | Methylcrotonoyl-CoA carboxylase subunit alpha, mitochondrial |
| **CLU** | P10909 | Clusterin | **MCM2** | P49736 | DNA replication licensing factor MCM2 |
| **COX6B1** | P14854 | Cytochrome c oxidase subunit 6B1 | **MPST** | P25325 | 3-mercaptopyruvate sulfurtransferase |
| **COX6C** | P09669 | Cytochrome c oxidase subunit 6C | **NDUFAB1** | O14561 | Acyl carrier protein, mitochondrial (ACP) |
| **CPT1A** | P50416 | Carnitine O-palmitoyltransferase 1, liver isoform | **NDUFB1** | O75438 | NADH dehydrogenase [ubiquinone] 1 beta subcomplex subunit 1 |
| **CPT2** | P23786 | Carnitine O-palmitoyltransferase 2, mitochondrial | **NDUFB4** | O95168 | NADH dehydrogenase [ubiquinone] 1 beta subcomplex subunit 4 |
| **CROT** | Q9UKG9 | Peroxisomal carnitine O-octanoyltransferase | **NDUFB9** | Q9Y6M9 | NADH dehydrogenase [ubiquinone] 1 beta subcomplex subunit 9 |
| **CYCS** | P99999 | Cytochrome c | **NDUFS6** | O75380 | NADH dehydrogenase [ubiquinone] iron-sulfur protein 6, mitochondrial |
| **FTH1** | P02794 | Ferritin heavy chain | **NQO1** | P15559 | NAD(P)H dehydrogenase [quinone] 1 |
| **GPX4** | P36969 | Phospholipid hydroperoxide glutathione peroxidase | **PCNA** | P12004 | Proliferating cell nuclear antigen |
| **GSS** | P48637 | Glutathione synthetase | **PIPOX** | Q9P0Z9 | Peroxisomal sarcosine oxidase |
| **GSTA3** | Q16772 | Glutathione S-transferase A3 | **PRODH** | O43272 | Proline dehydrogenase 1, mitochondrial |
| **GSTA5** | Q7RTV2 | Glutathione S-transferase alpha-5 | **SC5D** | O75845 | Lathosterol oxidase |
| **GSTK1** | Q9Y2Q3 | Glutathione S-transferase kappa 1 | **SEPSECS** | Q6P6M7 | O-phosphoseryl-tRNA(Sec) selenium transferase |
| **GSTM2** | P28161 | Glutathione S-transferase Mu 2 | **SLC25A20** | O43772 | Mitochondrial carnitine/acylcarnitine carrier protein |
| **GSTM7** | P09211 | Glutathione S-transferase Mu 7 | **SOD1** | P00441 | Superoxide dismutase [Cu-Zn] |
| **GSTP1** | P40939 | Glutathione S-transferase P | **SOD2** | P04179 | Superoxide dismutase [Mn], mitochondrial |
| **HADH** | Q16836 | Hydroxyacyl-coenzyme A dehydrogenase, mitochondrial | **STAT3** | P40763 | Signal transducer and activator of transcription 3 |
| **HADHA** | P55084 | Trifunctional enzyme subunit alpha, mitochondrial | **TGM2** | P21980 | Protein-glutamine gamma-glutamyltransferase 2 |
|  |  |  | **UQCR10** | Q9UDW1 | Cytochrome b-c1 complex subunit 9 |

Lipoprotein lipase (LPL), which hydrolyses intracellular lipid stores to release fatty acids, was significantly upregulated (**Figure 2**A).^47^ Concurrently, the structural machinery of the mitochondrial carnitine shuttle (CPT1A^48^, SLC25A20^40^, CPT2^49^) is upregulated. This was paired with the systematic upregulation of core enzymatic drivers of mitochondrial β-oxidation (ACAD11^50^, HADH^51^, HADHA^51^, HADHB^51^) (**Table 2**). This mitochondrial response was mirrored by the parallel upregulation of proteins controlling peroxisomal fatty acid β-oxidation, including acyl-coenzyme A oxidase 1 (ACOX1)^52,53^ and peroxisomal carnitine O-octanoyltransferase (CROT), indicating that the peroxisome is a complementary source of energy substrates and acyl-carnitines for FAO.

This energetic pivot was supported by a significant enrichment in amino acid catabolism pathways (**Figure 2**D, **Table 2**). Stressed CHO cells selectively overexpressed pathways required to break down amino acids into acyl-CoA intermediates to support mitochondrial function and fatty acid oxidation (ACADSB^54^, ALDH6A1^55^, BCKDHA^56^, BDH2^57^, HIBADH^58^, MCCC1^59^).^60,61^ Peroxisomal sarcosine oxidase (PIPOX)^62,63^ which supports auxiliary amino acid breakdown, was similarly increased, reinforcing the global involvement of the peroxisomal compartment during nutrient starvation. This combined proteomic evidence indicated that late-stage cells undergo extensive metabolic remodelling, focusing their remaining mitochondrial and peroxisomal capacity on the extraction of energy from alternative lipid and amino acid substrates (**Figure 2**D-E). This metabolic remodelling aligns with a transition where CHO cells allocate energy and resources from mAb production to survival and cellular maintenance and survival.

This metabolic redirection toward optimised mitochondrial and peroxisomal FAO machinery acts as a clear bioenergetic constraint. While the upregulation of fatty acid transport machinery and catabolic enzymes supports emergency energy substrate generation, it fundamentally alters the biophysical landscape of the cellular membranes. We hypothesise, that this lipid flux drives a rapid phospholipid remodelling loop via the Lands Cycle.^64^ This is supported by the concurrent proteomic upregulation of LPL (**Figure 2**A), which actively hydrolyses intracellular lipid stores to release a volatile pool of free fatty acids^47^; and LPCAT3, which synthesises phospholipids from acetyl-CoA fatty acids to replenish membrane lipids (**Figure S28**).^65,66^ Remodelling structural membranes to favour fatty acid re-esterification under an accelerated mitochondrial oxidation state facilitates the generation of localised reactive oxygen species (ROS) near lipid bilayers, priming the cellular architecture for membrane peroxidation.^66,67^

This accelerated metabolic state generated high levels of intracellular oxidative strain, as evidenced by the significant enrichment of cellular responses to oxidative stress (**Figure 2**D). To locate the sources of this oxidative burden, we tracked specific proteomic networks involved in ROS generation, superoxide metabolism, and mitochondrial electron leakage (Figure S34). Protein subunits within keys sites for electron leakage^68^ including complex I (NDUFAB1, NDUFB1, NDUFB4, NDUFB9, NDUFS6)^69,70^, complex III (UQCR10)^70^, and complex IV (COX6B1, COX6C and CYCS)^70^ of the electron transport chain (ETC) (as well as proteins from complex II) are upregulated between days 5 – 12 in the stressed cultures (**Figure 2**B, S34). Combined with increased ROS biosynthetic proteins such as ACOX1 and superoxide response proteins such as mitochondrial superoxide dismutase 1 (SOD1)^71^ these findings suggest that high-flux alternative substrate oxidation and structural electron leakage along the ETC are the primary drivers of oxidative stress in late-stage CHO operations.

This cellular oxidation environment directly correlated with the activation of ferroptosis, an iron dependent, non-apoptotic regulated cell death mechanism characterised by the catastrophic accumulation of lipid peroxides across structural membranes.^17 72^ By day 10, a critical checkpoint where control cultures enter growth arrest and the stressed IFB culture undergoes rapid decline, both microenvironments demonstrated strong upregulation of pro-ferroptotic signalling proteins (**Figure 2**C). However, the high-stress IFB condition demonstrated an accelerated and far more pronounced response: overexpressing key negative regulators of ferroptosis, including NAD(P)H quinone dehydrogenase 1 (NQO1)^73^, ferritin (FTH1) and lathosterol oxidase (SC5D). Additionally, there is significant upregulation of the enzyme for the biosynthesis of selenocysteine (SEPSEC) (**supplementary data**).^74^ Selenocysteine is the essential catalytic amino acid within the active site of the anti-ferroptotic protein glutathione peroxidase 4 (GPX4).^75,76^ This early response indicated that highly seeded cells experience chronic, long-term ferroptotic stress forcing them to upregulate defensive iron-storage and antioxidant pathways well before macroscopic cell death is observed. Collectively, these proteomic signatures indicate that the high-stress phenotype is driven by an energetic crisis, cells undergo extensive metabolic remodelling to fuel mitochondrial FAO, creating an electron-leakage microenvironment that drives ROS generation and triggers terminal ferroptotic pathways (**Figure S28**).

### Extended Glycolysis and Redox Control Prevent Decline

In sharp contrast to the high-stress IFB model, cultures utilising enriched basal media (H6-B and H6-C) show significant down regulation of pro-ferroptotic signalling networks paired with a strong enrichment of glutathione-mediated redox defence systems (**Figure 3A-C, S6-15**). Both the H6-B and H6-C proteomes retained low baseline expression of alternative lipid transport and FAO proteins compared to the control. Both these higher viability phenotypes are grown within an enriched amino acid environment, so they can maintain robust intracellular amino acid pools. Glutathione is the core substrate required to neutralise lipid hyperoxidases, a process catalysed by GPX4. Both the H6-B and H6-C cultures utilise the enriched amino acid pool (specifically glycine) to actively drive glutathione synthesis via the upregulation of glutathione synthetase (GSS) on day 7 (**Figure 3D, S7, S12**). No concurrent increase was observed in glutamate cysteine ligase, which produces the key intermediate γ-glutamylcysteine. However, the media also features enriched glutamate and cysteine, suggesting that these enriched amino acids increase ferroptotic control through enhanced glutathione production.

**Figure 3:**
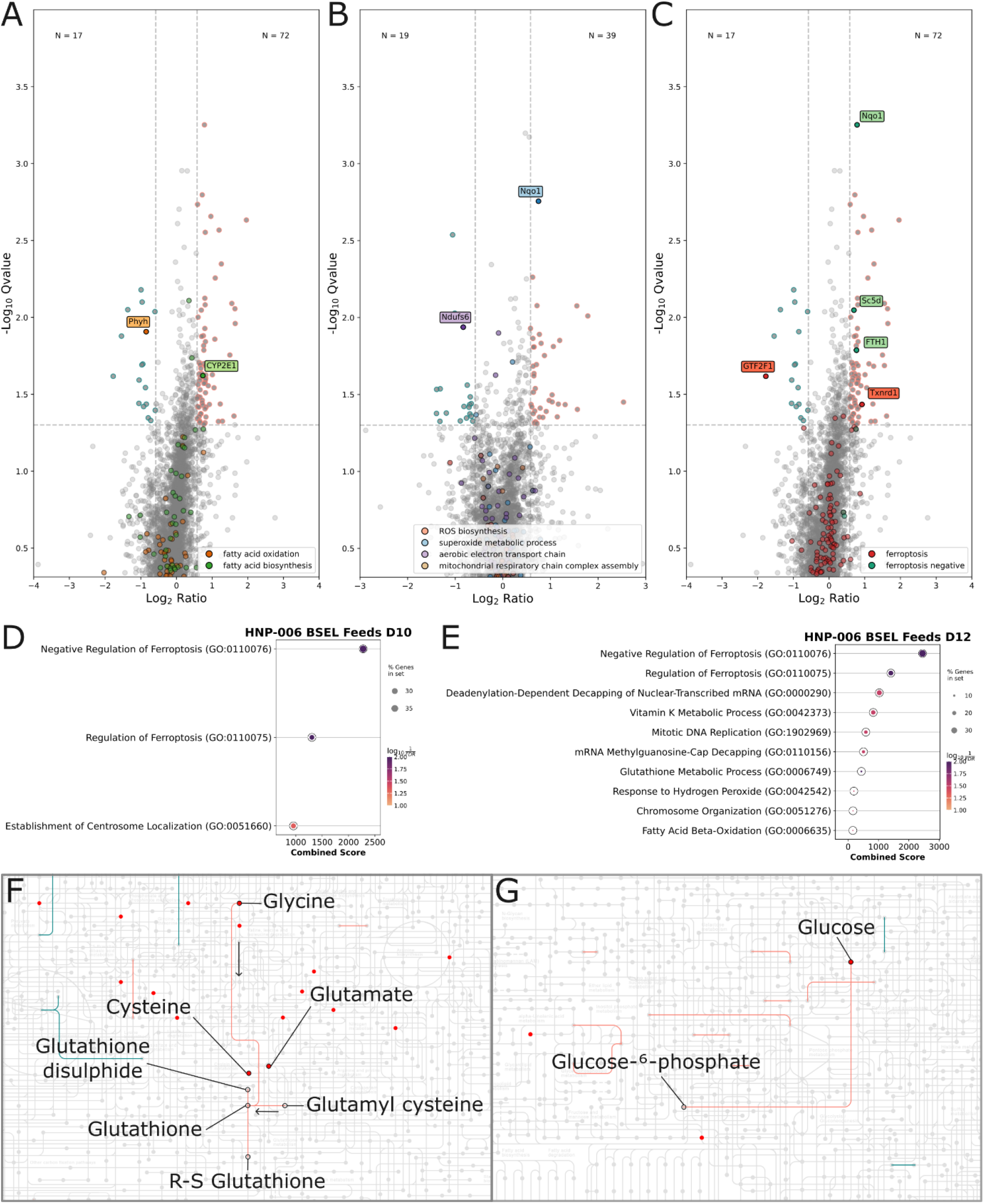
Differentially abundant hits between the control (blue) and high viability (pink) phenotypes on day 10, with GO biological processes pathways **A)** fatty acid biosynthesis (green) and fatty acid oxidation (orange), day 10 **B)** ROS biosynthesis (red), superoxide metabolic process (blue), aerobic electron transport chain (purple) and mitochondrial respiratory chain complex (tan), day 12 **C)** positive (red) and negative (green) ferroptotic control, day 10 **D)** gene set enrichment for H6-B on day 10 and **E)** day 12. **F)** KEGG map showing upregulated glutathione biosynthesis and **G)** higher glucose utilisation, with metabolites present at higher concentrations within the enriched feed marked in red, and metabolites of interest labelled.

Futhermore, the high-viability state was characterised by a distinct absence of FAO associated oxidative enzymes and mitochondrial electron-leakage markers (**Figures S29-30, S31-32**). The H6 basal medium contains higher baseline levels of glucose, and in the early culture days, the H6-B condition demonstrated an increased utilisation of glucose to produce glucose-6-phosphate (G6P), the first step in production of energy via glycolysis. By sustaining glycolysis longer into the cultivation lifecycle, these enriched cultures defer the necessity of switching to alternative, ROS-heavy lipid catabolism, therby minimising mitochondrial electron leakage and maintaining low ferroptotoic susceptibility.

Feed enrichment also appears to have an impact on viability, but only if enriched media is supplied. H6-B has enhanced viability compared to H6-C, however there is no significant difference between J5-B or J5-C (**Figure 1**A). The BSEL feed has higher levels of arginine, asparagine, histidine and lysine, and lower levels of aspartate. There are no differences in proteins involved in the metabolism of any of these amino acids between medias with different feeds (**Figures S35-36**), so it is unclear why feed enrichment, but only when combined with media enrichment, leads to higher viability.

It should be noted that while suppression of lipid metabolism and continuation of glycolytic flux leads to increased viability, biomass accumulation does not translate to increased mAb production. That suggests that controlling the metabolic switch occuring on day 10 offers candidate strategies for the continuation of productive CHO cell states.

### Lipid Remodelling Sensitises the Cells to Ferroptosis

To validate our proteomic model of lipid-driven cellular stress, untargeted intracellular metabolomics was performed to identify the small-molecule features driving the separation between the H6-B, IFB and control bioreactors (**Figure 4**).

**Figure 4:**
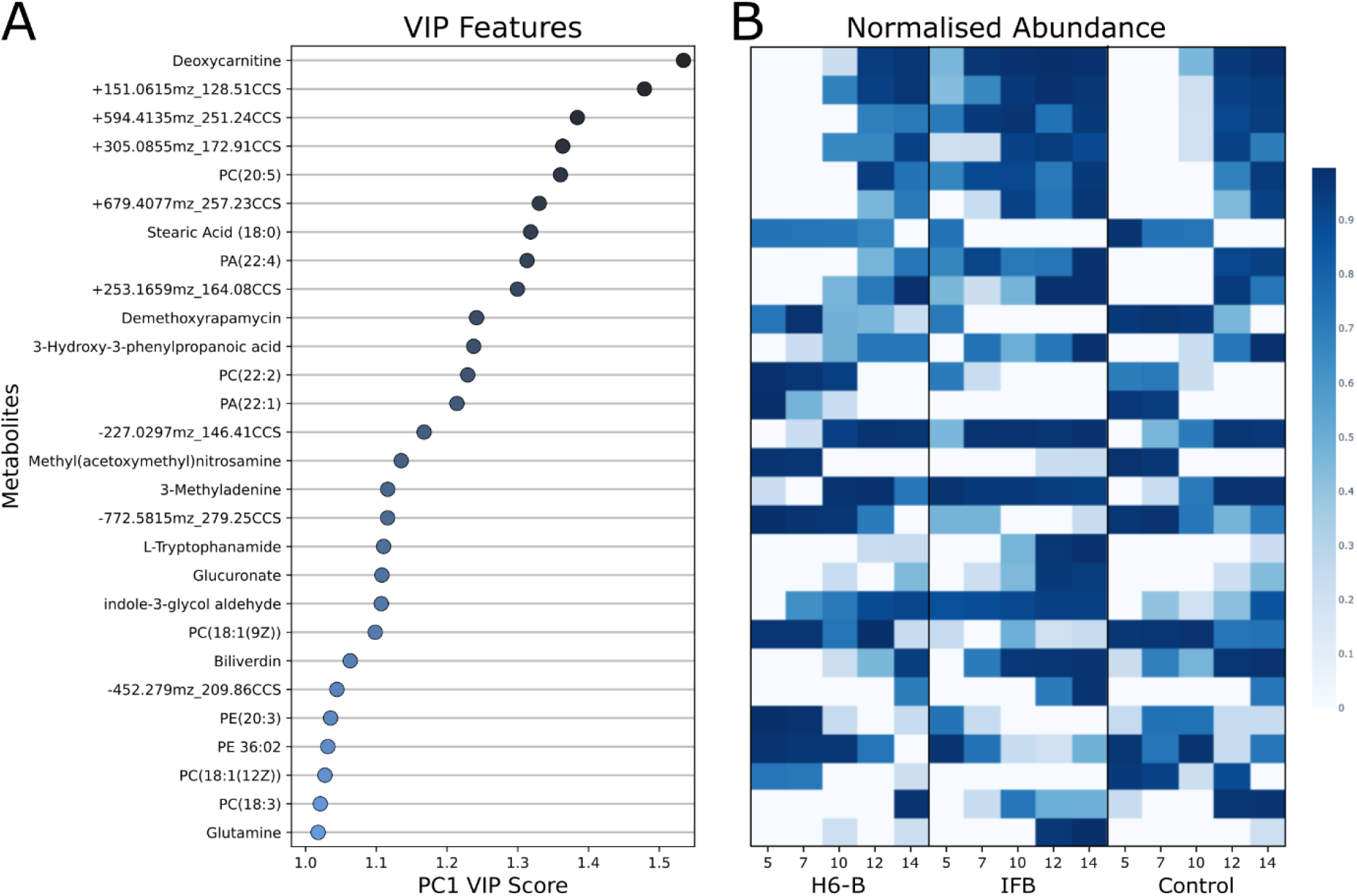
Selected intracellular metabolomics data **A)** Features with a VIP score > 1 **B)** normalised abundance of the VIP features between H6-B, IFB and control bioreactors. Where features could not be annotated, they are named according to their ionisation polarity, *m/z* and CCS values.

The specific intracellular metabolite driving the largest separation across the datasets was deoxycarnitine, which increased in abundance over time across all bioreactors, with the high-stress IFB condition accumulating the highest levels (**Figure 4**). As a direct baseline precursor in the carnitine biosynthetic pathway, the accumulation of deoxycarnitine provides further evidence that stressed cells aggressively expand their carnitine pools to facilitate the mitochondrial import of long-chain fatty acids for FAO^77,78^, matching the observed upregulation of mitochondrial carnitine shuttle proteins (**Table 2**).

Intracellular lipid profiling revealed a significant shift in membrane composition during the late-stage stress phase. Nine fatty acids were identified with significant VIP scores, including long chain phosphatidylcholines (PC), phosphatidic acids (PA), phosphatidylethanolamines (PE) and stearic acid (SA) (**Figure 4**). A variety of unsaturated, monounsaturated and polyunsaturated fatty acid (UFA, MUFA, PUFA) containing phospholipids are observed to decrease in abundance (PA(22:1), PC(18:1(9Z)), PC(18:3), PC(22:2), PE(20:3), PE(36:02) and stearic acid (18:0)). Highly polyunsaturated fatty acid (HPUFA) and PUFA containing phospholipids (PA(22:4), PC(20:5) and PC(18:3)) increase in abundance. This structural lipidomic remodelling aligns with the selective beta-oxidation of lower-saturation fats during accelerated FAO, causing a relative enrichment of HPUFAs in the remaining membrane pool via the Land’s Cycle driven by LPCAT3.^64,66,67^ These HPUFAs are highly susceptible to radical-mediated peroxidation therefore their disproportionate accumulation may drive ferroptotic susceptibility of lipid bilayers. Separate from the longitudinal trends, the high-stress IFB condition maintained higher levels of HPUFA containing phospholipids and lower levels of lower saturation fatty acid containing phospholipids, than control and H6-B conditions. This suggests that the switch to FAO from glycolysis sensitises the cells to ferroptosis through the enrichment of peroxidation sensitive lipids in the membranes and the production of a highly oxidative environment.

We also observed an increase in intracellular glutamine throughout the cultivation lifecycle across all tested bioreactor conditions (**Figure 4**B). Glutamine depletion acts as a known driver of the lactate metabolic switch in CHO cells.^79^ However, as glutamine was replenished in these fed-batch feeding strategies, it accumulated inside the cytoplasm without being fully utilised by the primary biosynthetic machinery. Accumulation of glutamine (and transferrin) have been shown to induce ferroptosis.^80^ This accumulation profile indicated that persistent nutrient overfeeding under uncoupled glycolytic flux inadvertently contributes to the induction of late-stage ferroptotic stress.

## Conclusion

Increasing the productivity of CHO cells remains a critical bottleneck for reducing manufacturing costs and improving process sustainability of biopharmaceuticals. By leveraging parallel proteomic and metabolomic profiling across highly divergent fed-batch microenvironments, we have demonstrated that the late-stage transition into a stressed, low-production state is fundamentally governed by a metabolic redirection toward mitochondrial and peroxisomal fatty acid β-oxidation. This energetic pivot fuels an active membrane remodelling cascade via the Lands Cycle. The subsequent accumulation of peroxidation-susceptible HPUFAs within structural bilayers primes the cellular architecture for ROS attack, propagated by ETC electron leakage this triggers terminal ferroptotic cell death pathways (**Figure 5**).

**Figure 5:**
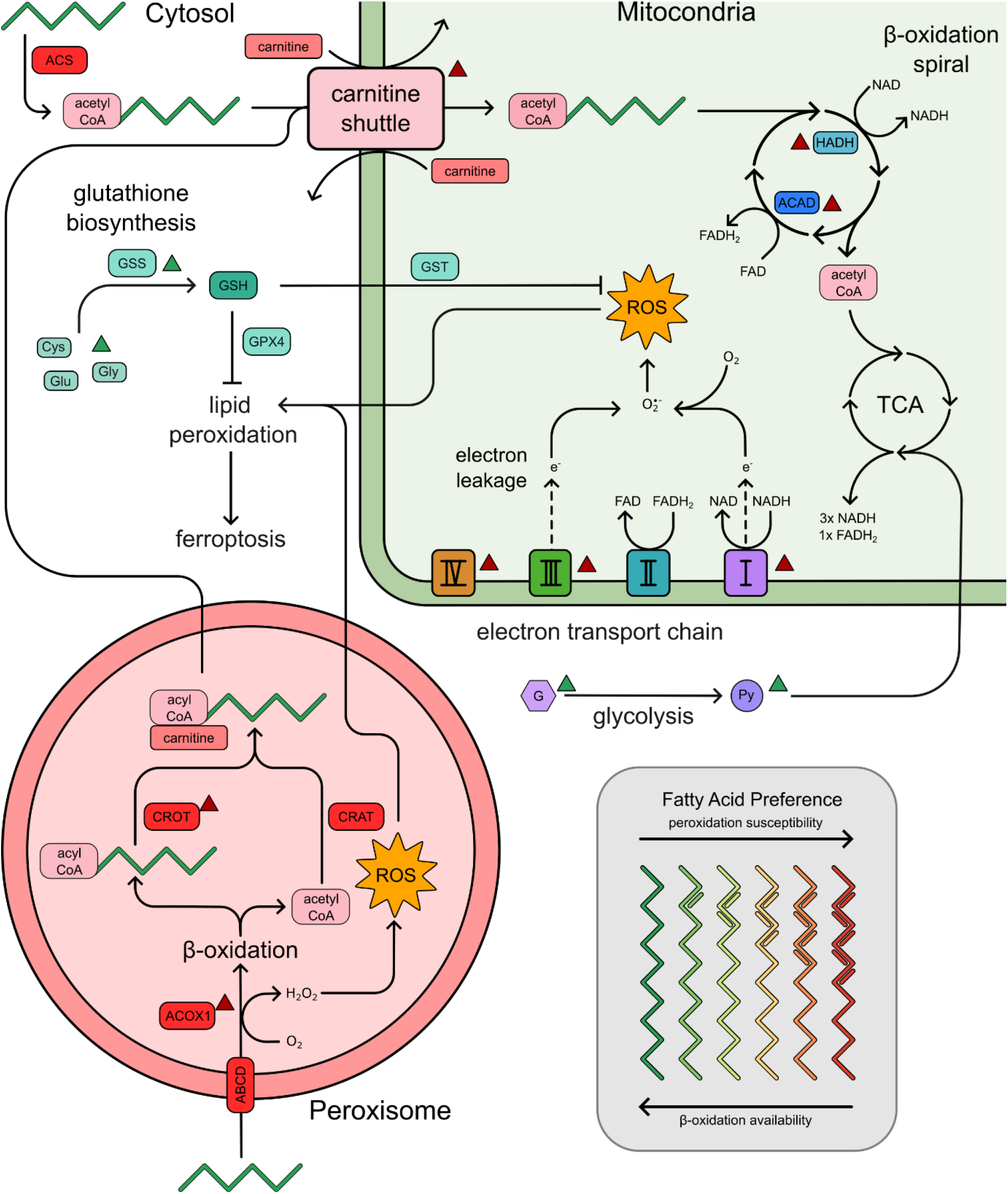
Mechanistic figure linking the connections between cytosolic, mitochondrial and peroxisomal metabolism and the induction of lipid peroxidation via ROS. Proteins or metabolites upregulated in either the high viability or stressed states are marked with a green or red arrow respectively. Link between fatty acid oxidation and the production of mitochondrial ROS, and the lower reliance on oxidative metabolism for energy production.

While advanced, nutrient-enriched cultivation frameworks (such as H6-B) successfully delay this phenotypic decline, they expose a stark biological trade-off. These specialised formulations leverage an alternative amino acid metabolic bypass to sustain the TCA cycle via the malate-aspartate shuttle and bolster glutathione-mediated redox networks (GPX4). However, the considerable metabolic energy demanded to fuel these protective defence frameworks heavily depletes the cellular resources required to sustain productivity. Consequently, volumetric product yields stagnate despite improved biomass accumulation and viability.

Our work here offers insights into candidate strategies for increasing the productivity of biotherapeutic production within CHO cells. Productivity and product quality within the end stage of bioreactors can be improved through limiting the reliance on the transition from glycolytic flux towards fatty acid metabolism. Feed manipulation to balance this energetic shift, such as supplying feeding with selenocysteine or other amino acids required for the control of the redox environment through GPX4. One might think that supplying further glucose would allow cells to maintain glycolysis preventing this transition, however this is not the case. The enriched feed bioreactors still transition to a lower productivity state where energy is shifted towards survival away from productivity even when additional glucose and amino acids are present. The work here highlights that this conserved metabolic transition is a key point where productivity declines. Identification of the metabolic constraints could allow for the continuation of the productive state, delaying the transition into where energetic resources are allocated for the maintenance of survival rather than mAb production.

We acknowledge that this study is inherently association-based and bounded by the current structural limitation of commercial CHO reference genomes: protein identities were established using consensus mammalian databases and assume strict functional conservation with *Cricetulus griseus* orthologues. Future investigation will focus on executing targeted gene knockouts or chemical inhibition studies of the key enzymatic gatekeepers regulating lipid fluxes and alternative amino acid routing to definitively validate this regulatory model. Additionally, deep coverage lipidomic tracking will be required to fully map dynamic phospholipid fluxes during the FAO transition. Ultimately, our findings establish that suppressing alternative lipid catabolism and stabilising glycolytic flux are paramount to maintaining prolonged biomanufacturing efficiency. Decoupling cellular longevity from recombinant protein production will require next-generation cell line engineering and feed formulations designed specifically to insulate host networks against the carnitine/fatty acid-ferroptosis axis.

## Supplementary Information & Data

Supplementary figures showing the volcano plots, GSEA and KEGG maps for HNP-006 BSEL, HNP-006 Cytiva, Intensified fed-batch, JM-05B BSEL and JM-05B Cytiva compared to Platform, and HNP-006 BSEL compared to HNP-006 Cytiva. Supplementary data includes tables containing all significant hits within volcano plots from all comparisons, all Enrichr reports for biological processes and cellular compartments, the mapped pathways used to produce the KEGG maps, and the spreadsheet used to map UniProt IDs to KEGG KO numbers.

## Data Availability

The raw mass spectrometry data, spectral libraries, and analysis software parameters have been deposited to the ProteomeXchange consortium via the MassIVE repository with the dataset identifier MSV000102138.

## Acknowledgements

We thank Tessa Moses and the technical staff at Edinomics for running and analysing the metabolomics samples, and the support from the Mass Spectrometry and Separations Facility of the Faculty of Science and Engineering and the Future Biomanufacturing Research Hub at the University of Manchester. This work was supported by a Prosperity Partnership grant (EP/V038095/1) funded by the Engineering and Physical Research Council (EPSRC) and FUJIFILM Biotechnologies, the University of Manchester, and instrumentation was funded by the BBSRC (BB/W019892/1).

