## Supplementary figures and images for "Fatty Acid β-oxidation and Ferroptosis Define a Survival-Productivity Trade Off in CHO fed-Batch Bioreactors"

### GO_Biological_Process_2025.Mouse.enrichr.reports.pdf

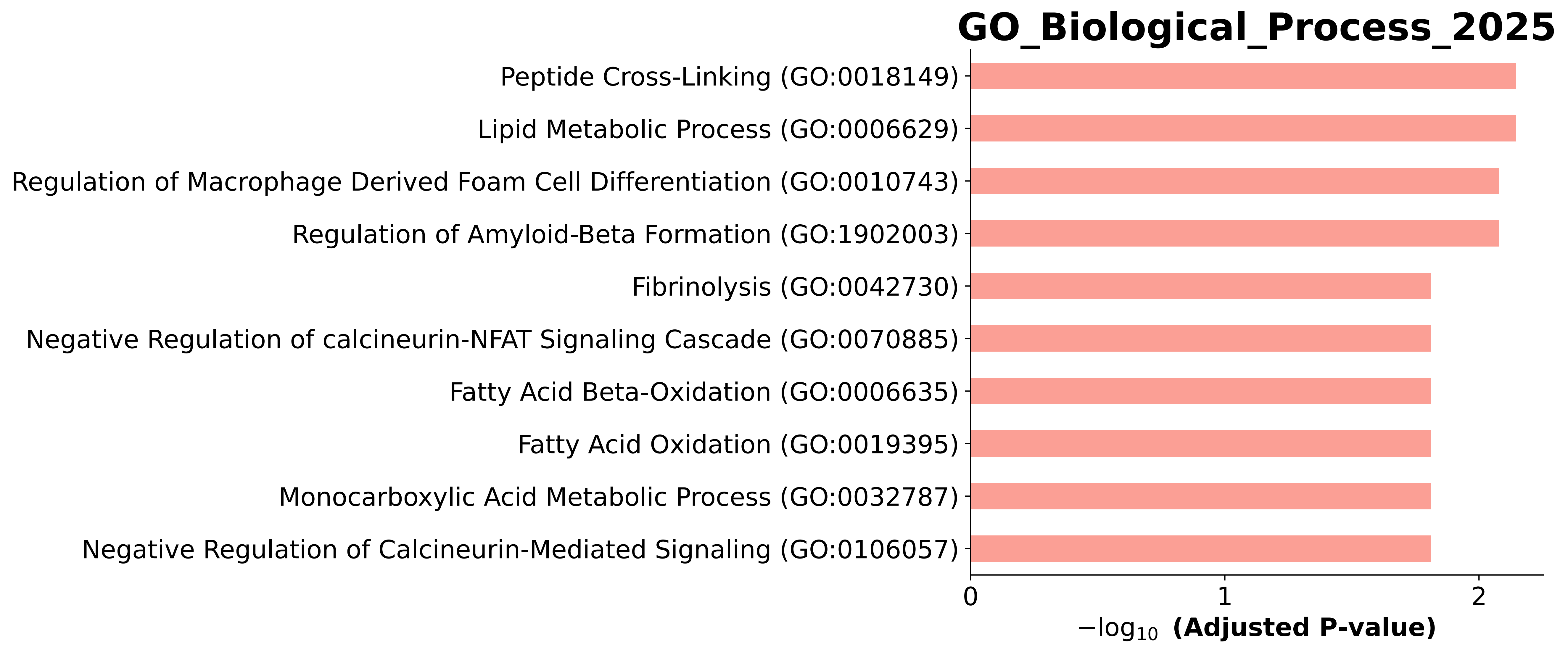

### GO_Biological_Process_2025.Mouse.enrichr.reports.pdf

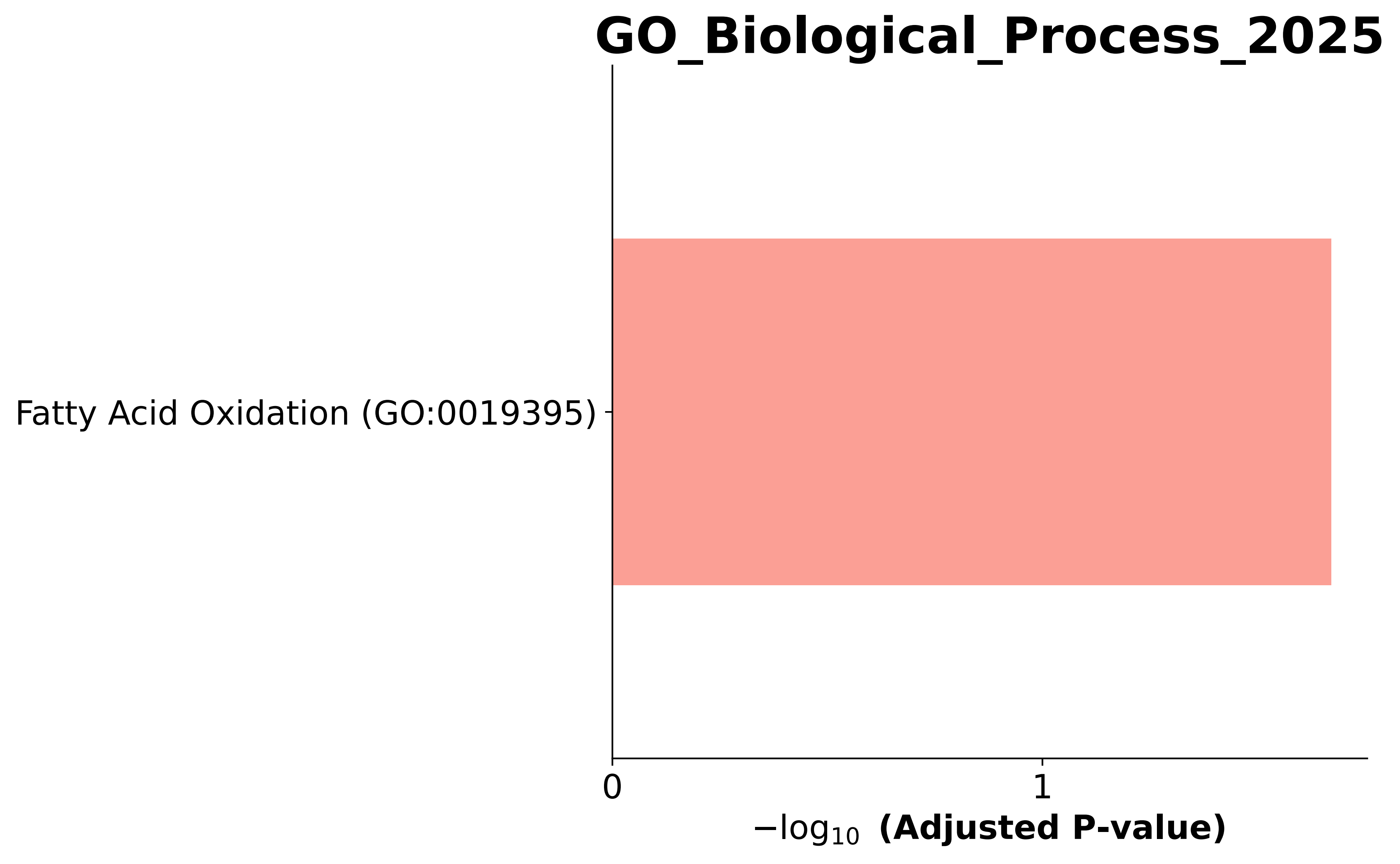

### GO_Biological_Process_2025.Mouse.enrichr.reports.pdf

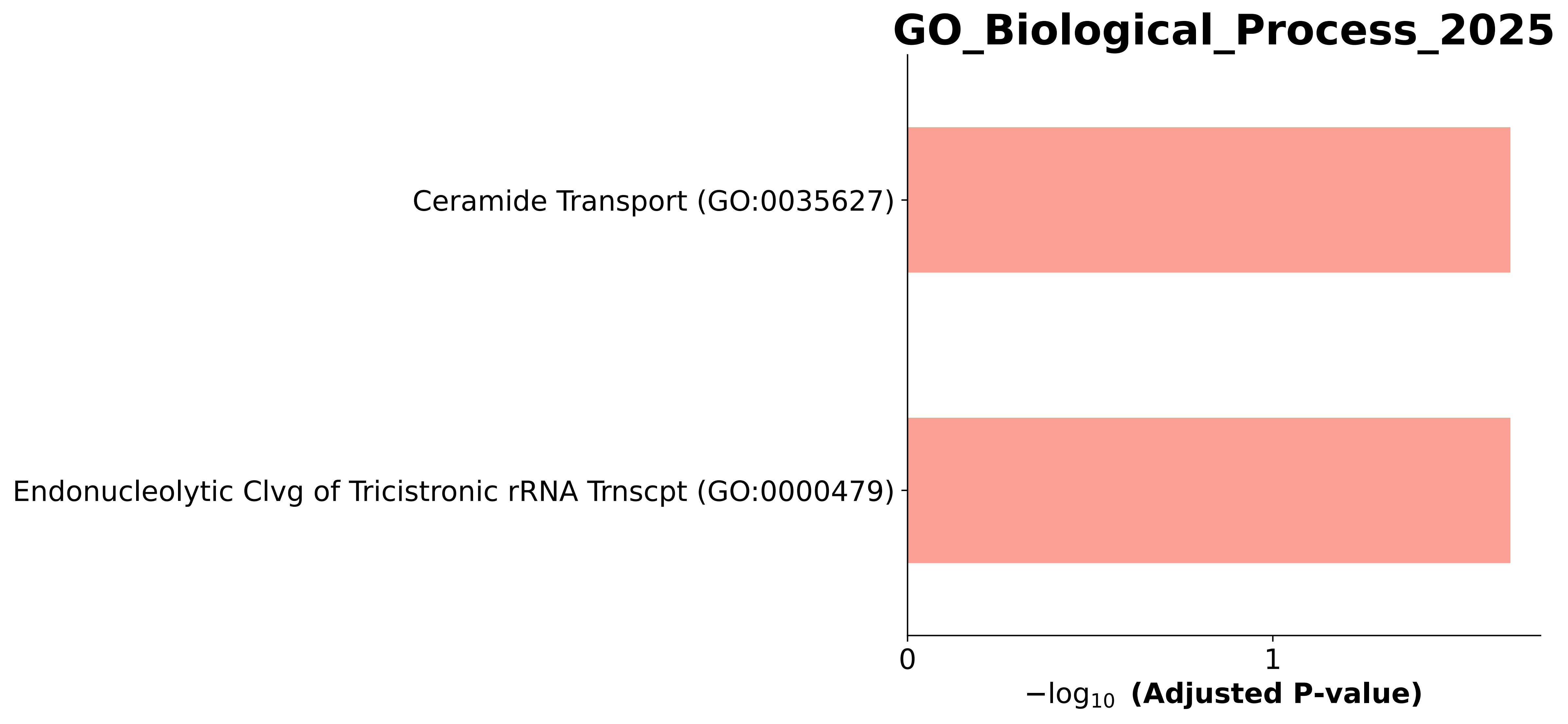

### GO_Biological_Process_2025.Mouse.enrichr.reports.pdf

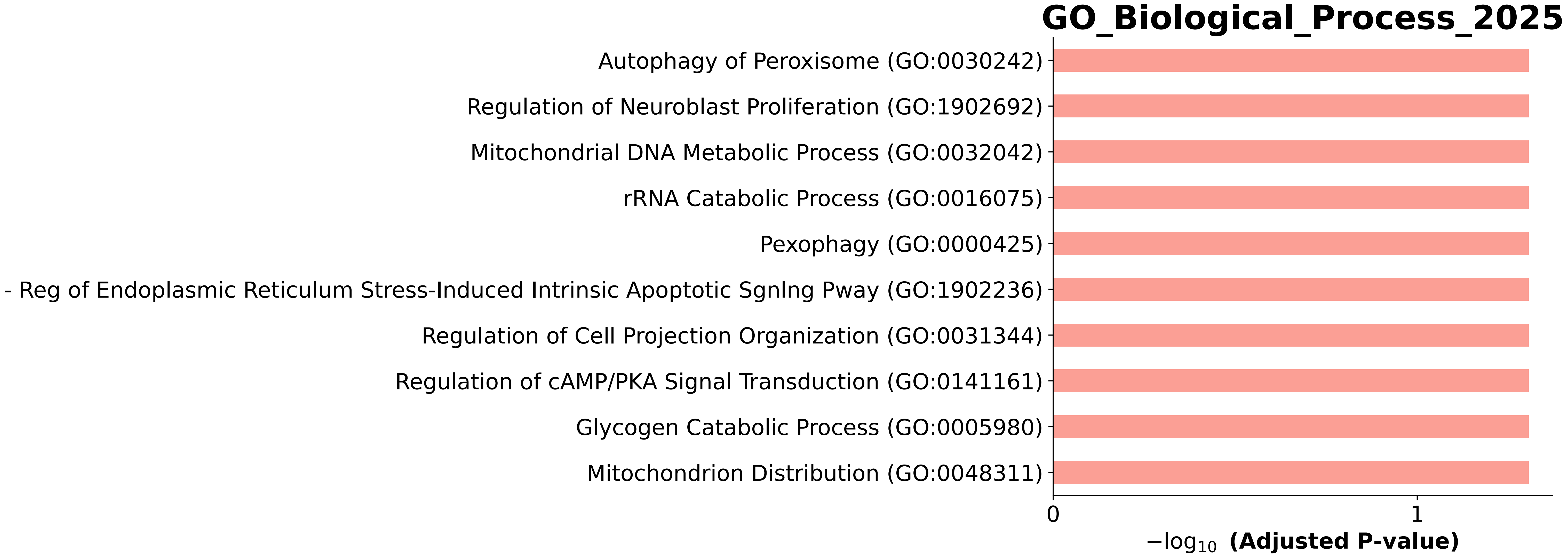

### GO_Biological_Process_2025.Mouse.enrichr.reports.pdf

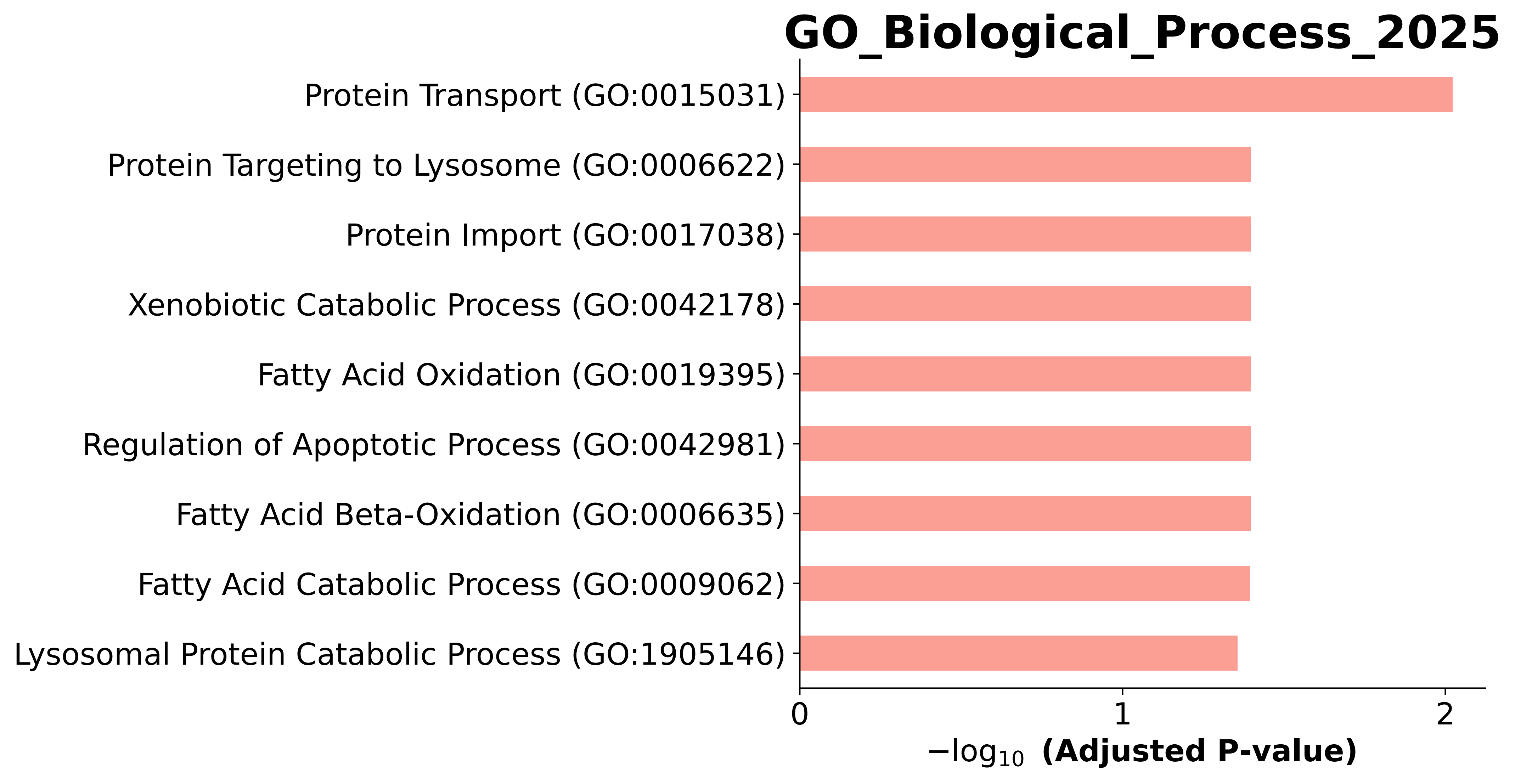

### GO_Biological_Process_2025.Mouse.enrichr.reports.pdf

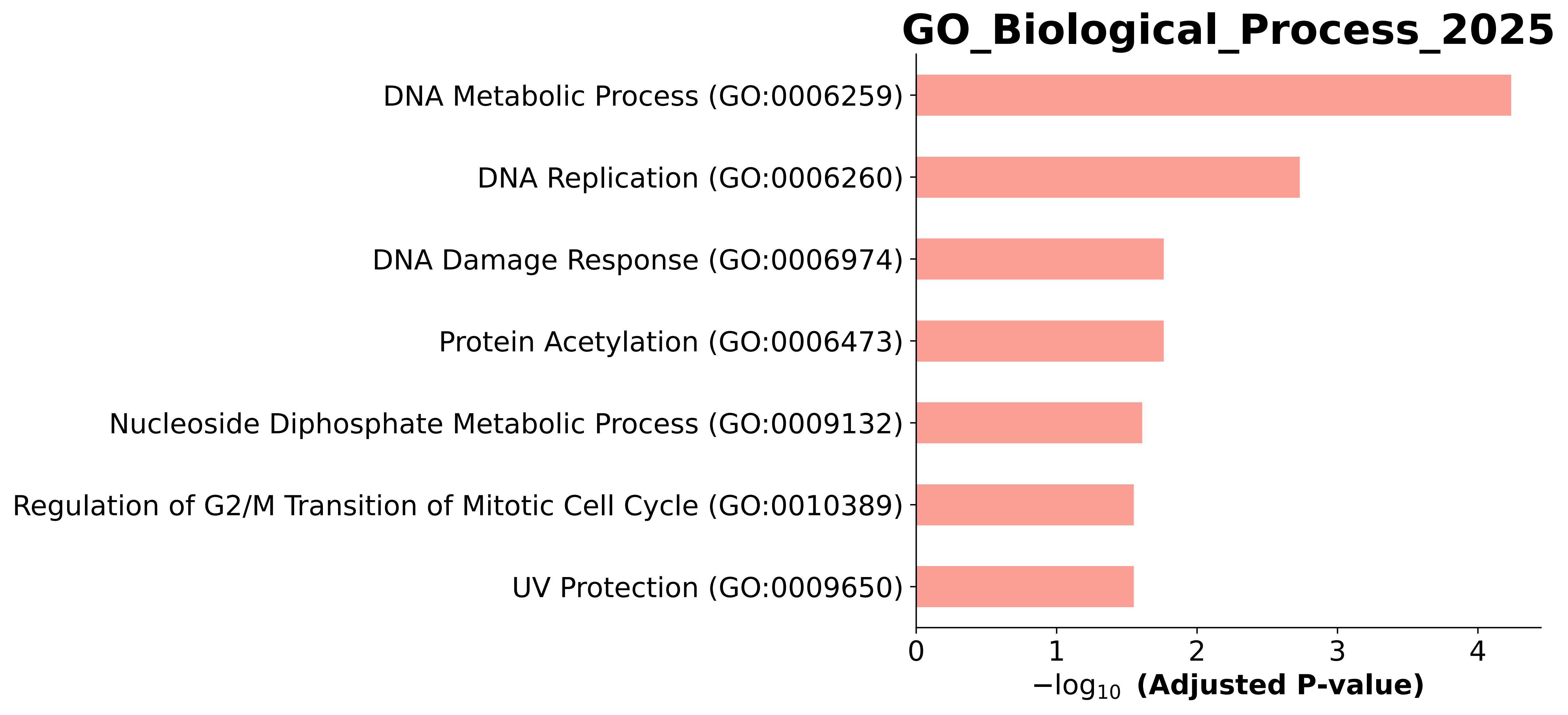

### GO_Biological_Process_2025.Mouse.enrichr.reports.pdf

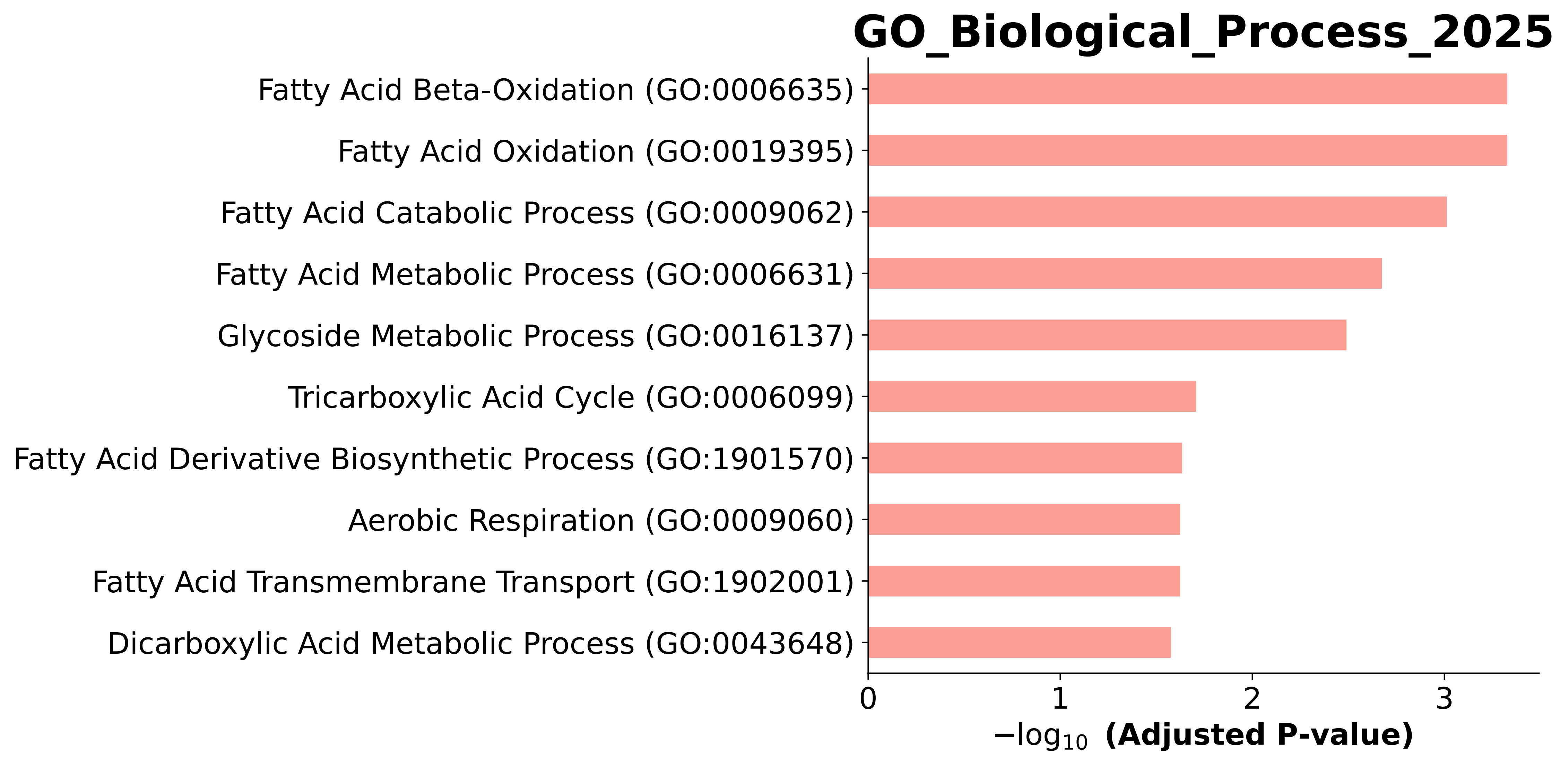

### GO_Biological_Process_2025.Mouse.enrichr.reports.pdf

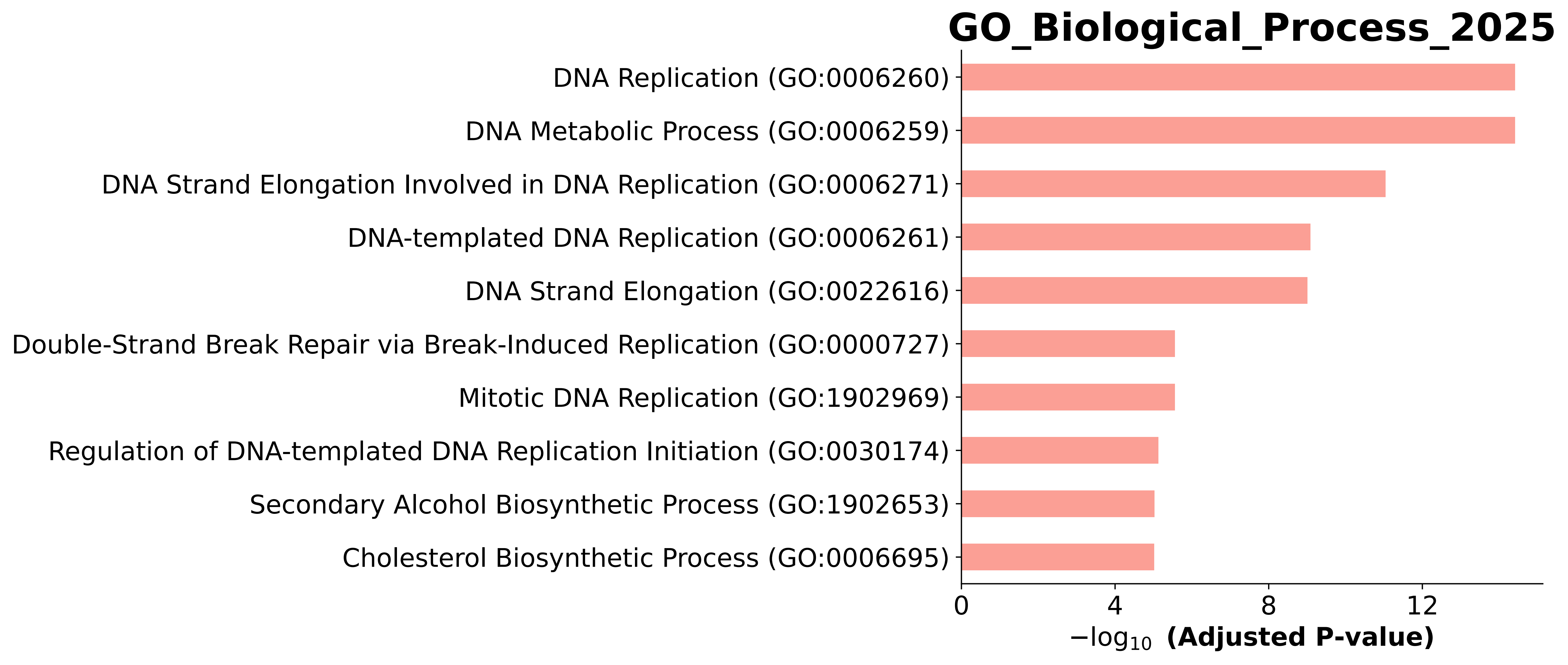

### GO_Biological_Process_2025.Mouse.enrichr.reports.pdf

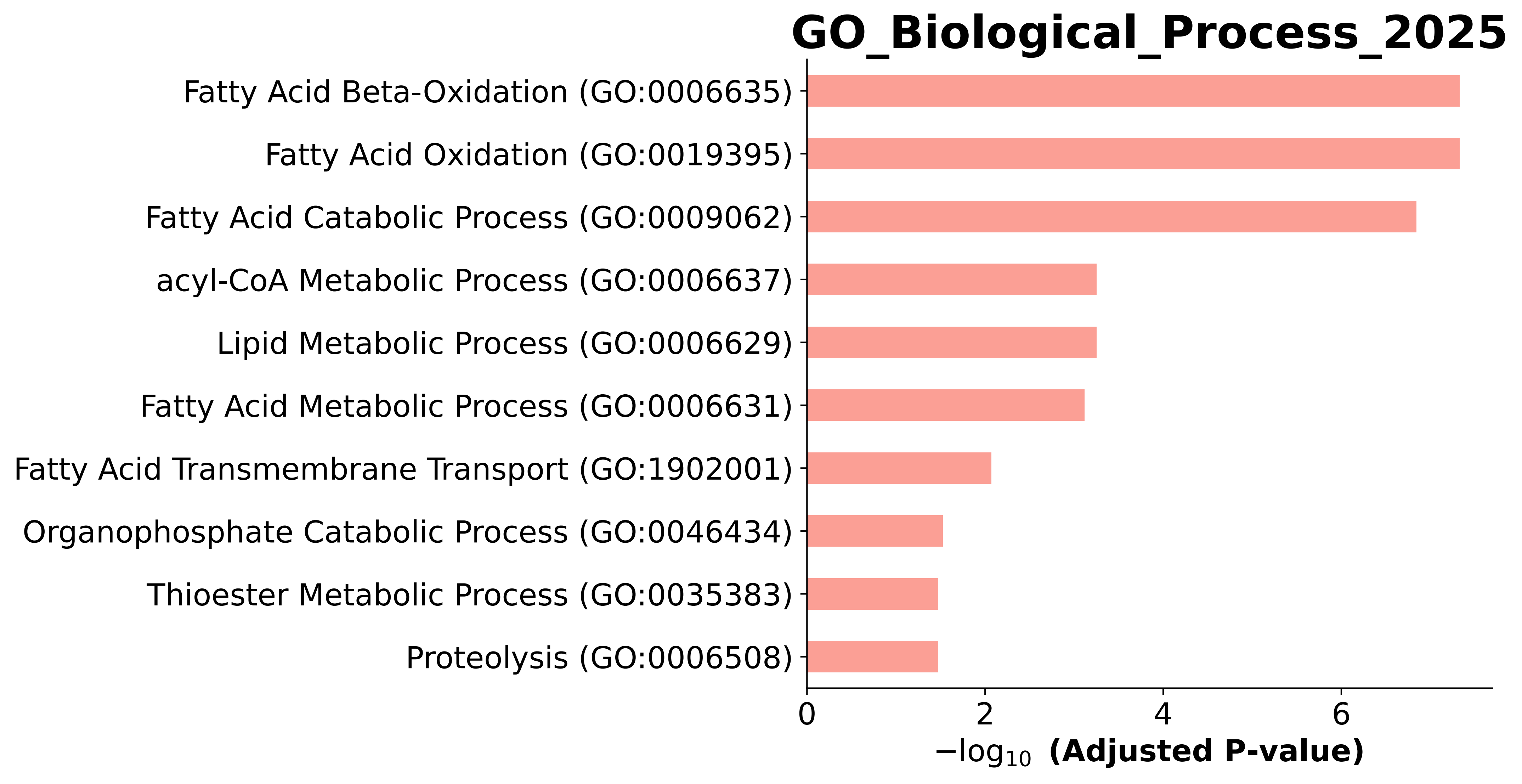

### GO_Biological_Process_2025.Mouse.enrichr.reports.pdf

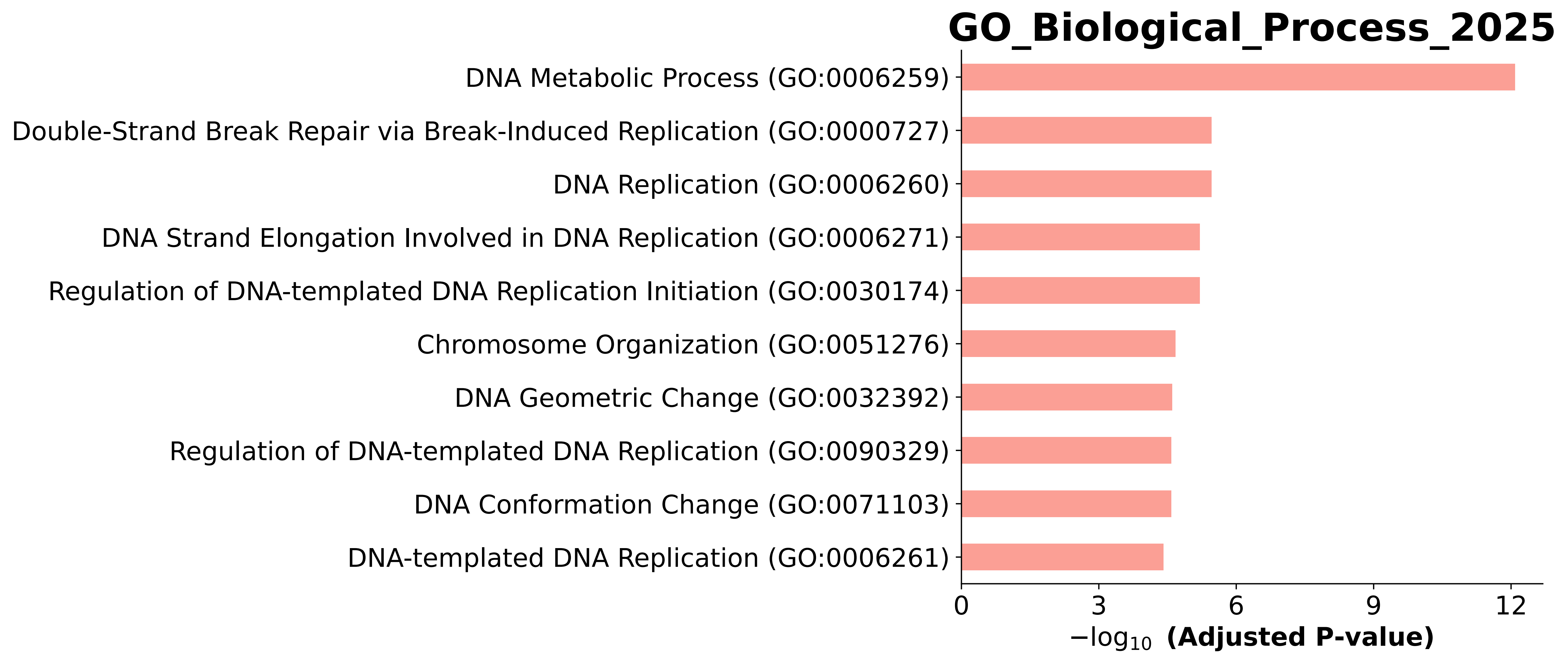

### GO_Biological_Process_2025.Mouse.enrichr.reports.pdf

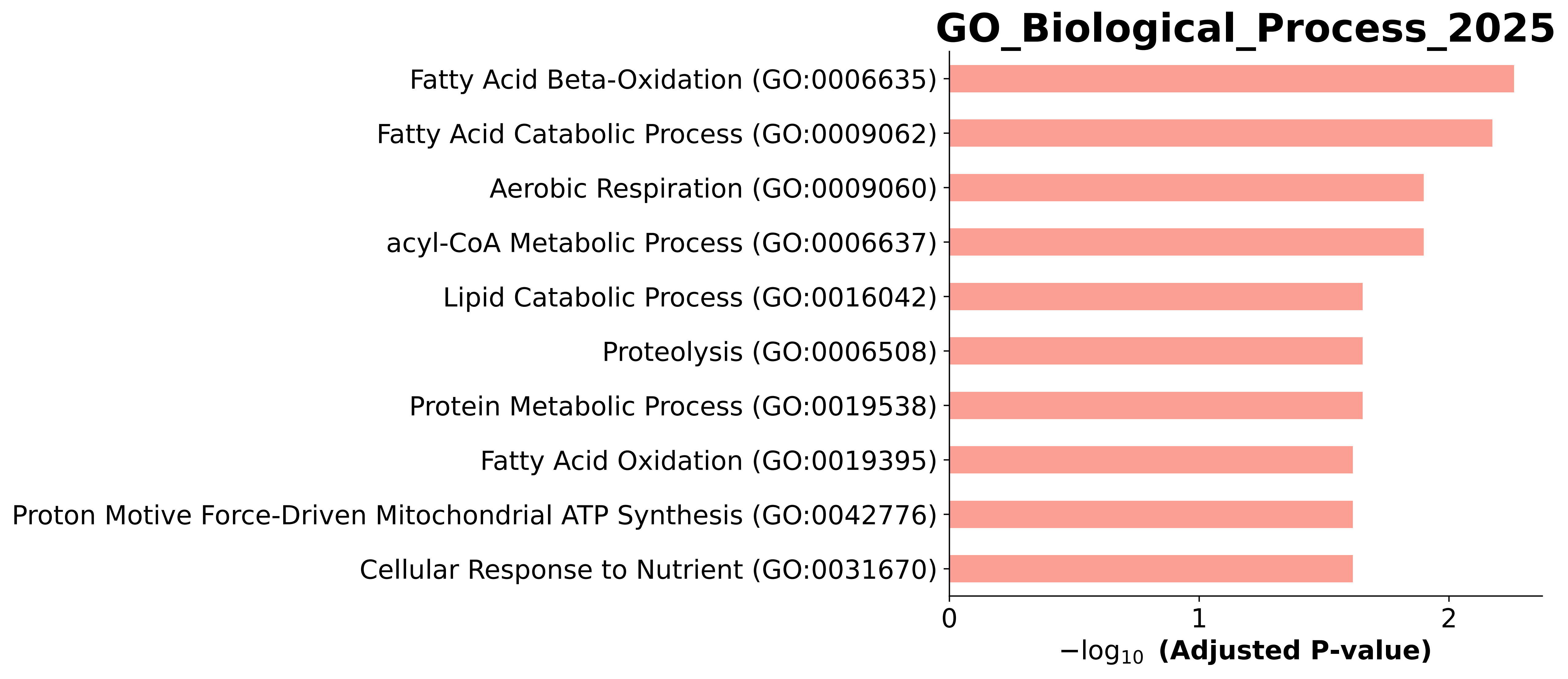

### GO_Biological_Process_2025.Mouse.enrichr.reports.pdf

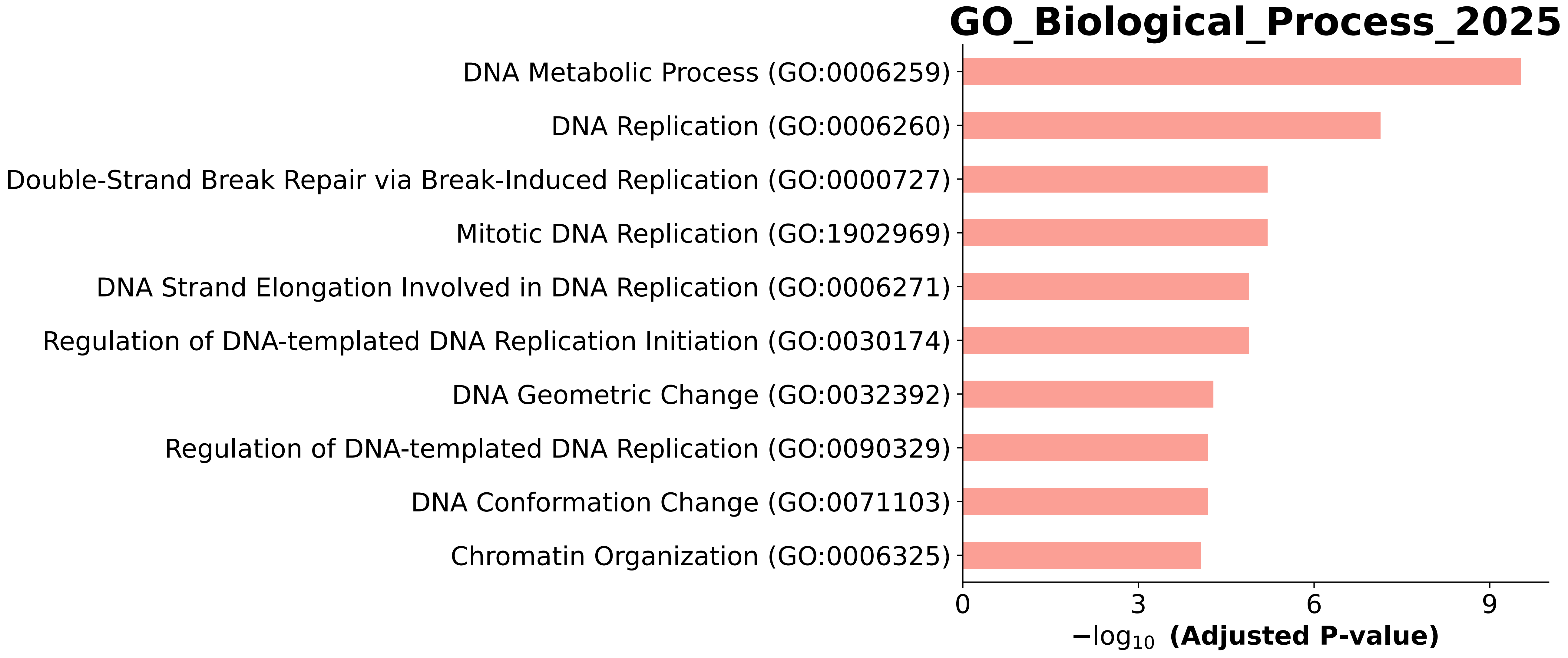

### GO_Biological_Process_2025.Mouse.enrichr.reports.pdf

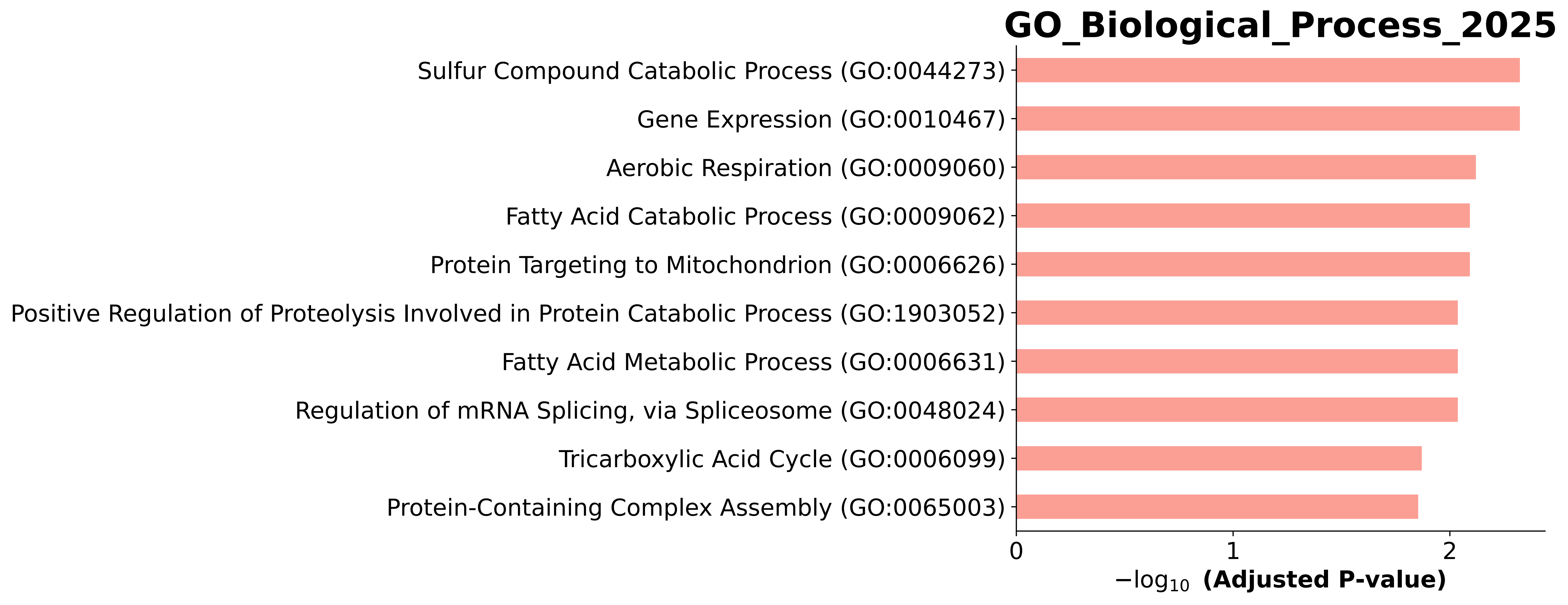

### GO_Biological_Process_2025.Mouse.enrichr.reports.pdf

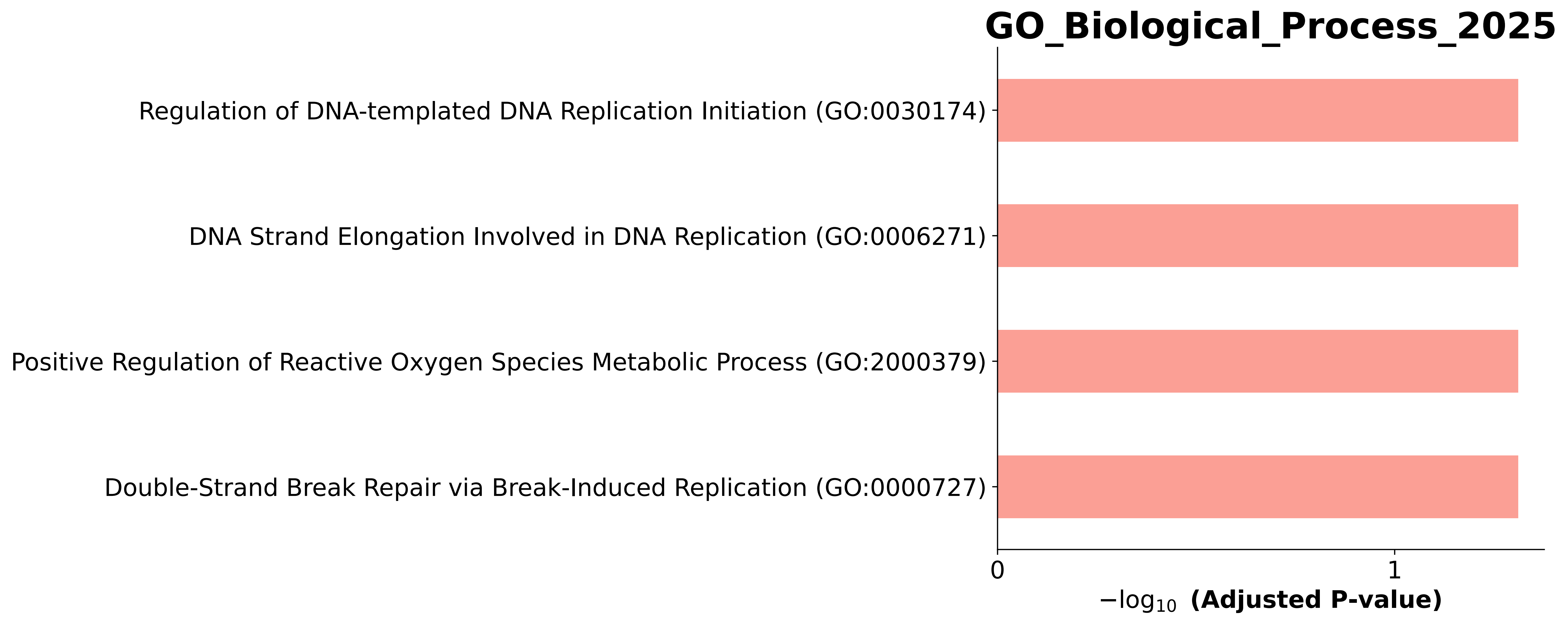

### GO_Biological_Process_2025.Mouse.enrichr.reports.pdf

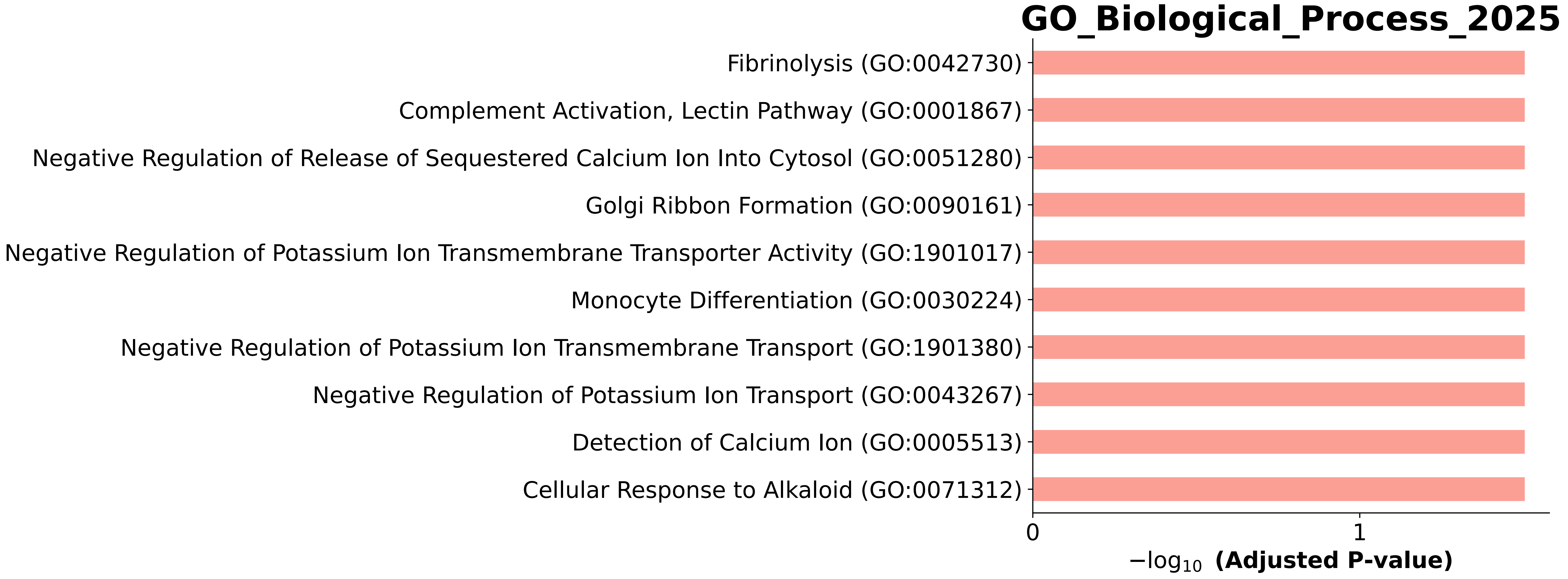

### GO_Biological_Process_2025.Mouse.enrichr.reports.pdf

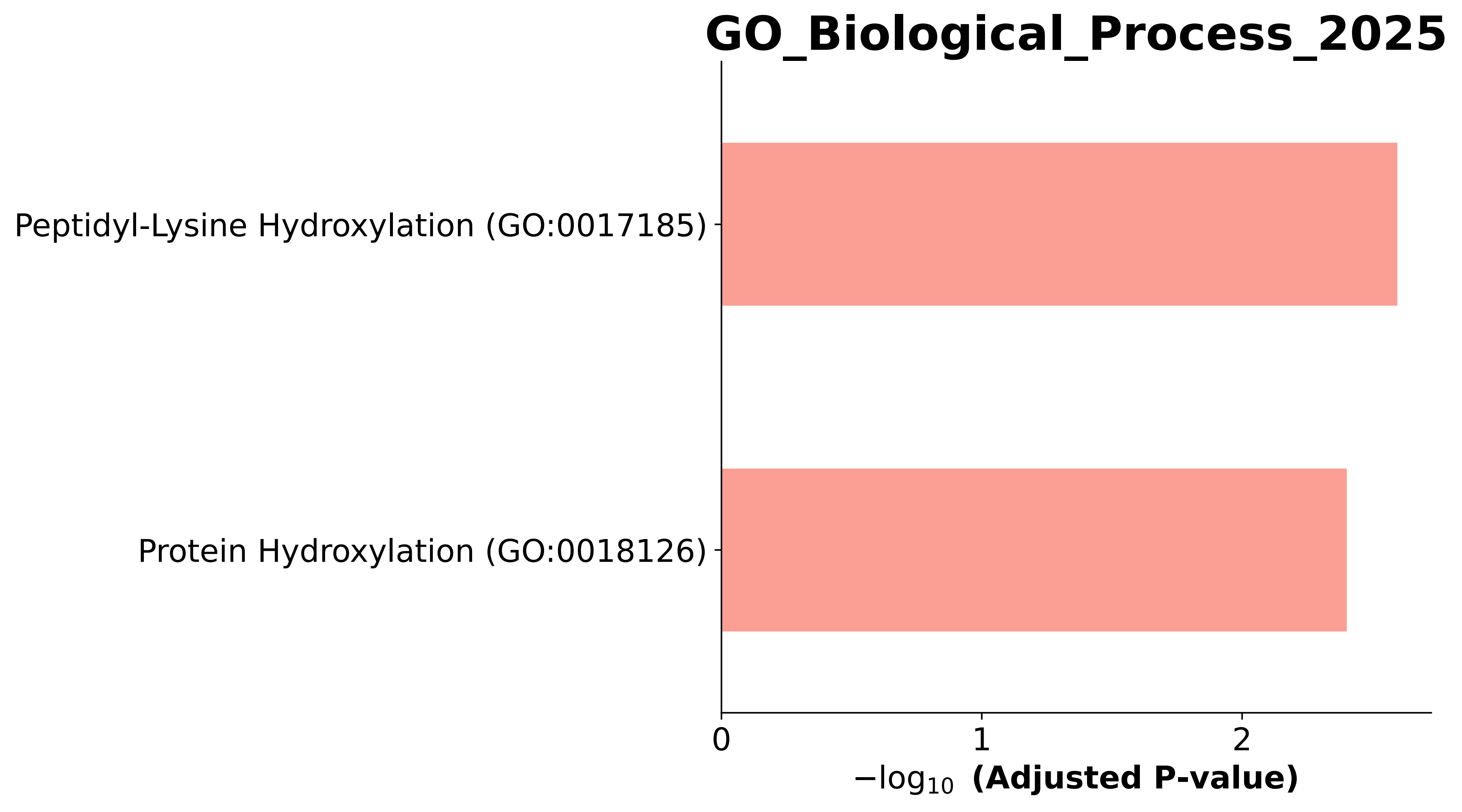

### GO_Biological_Process_2025.Mouse.enrichr.reports.pdf

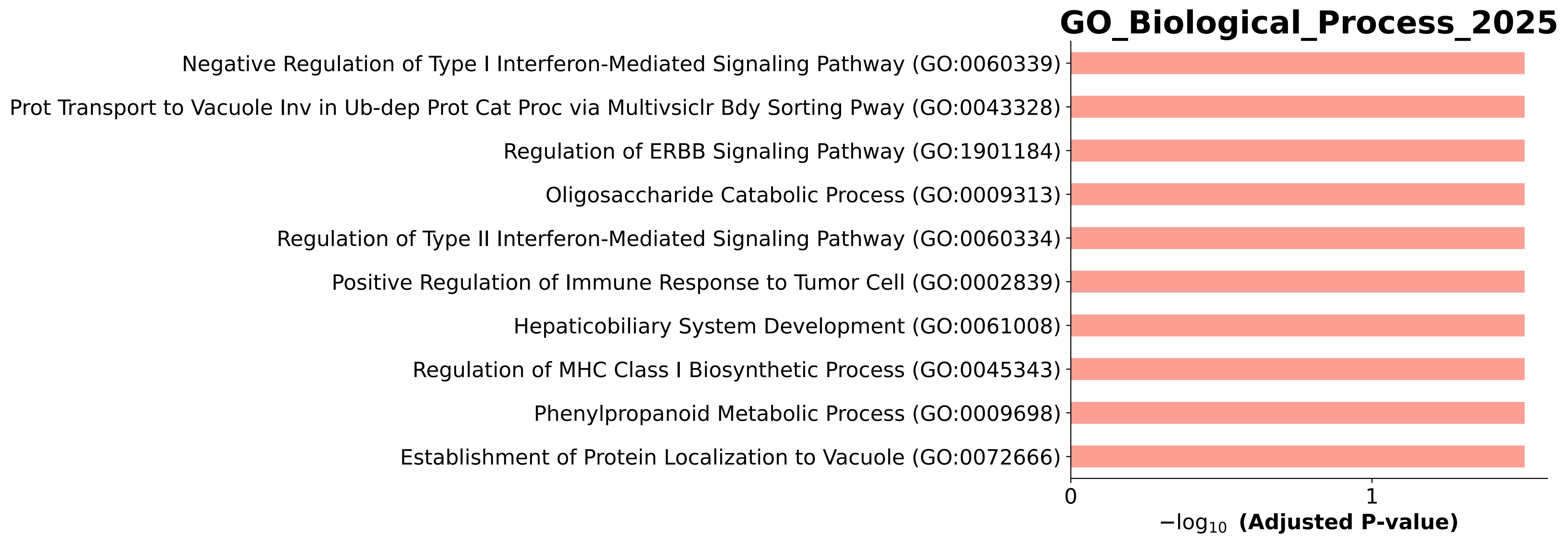

### GO_Biological_Process_2025.Mouse.enrichr.reports.pdf

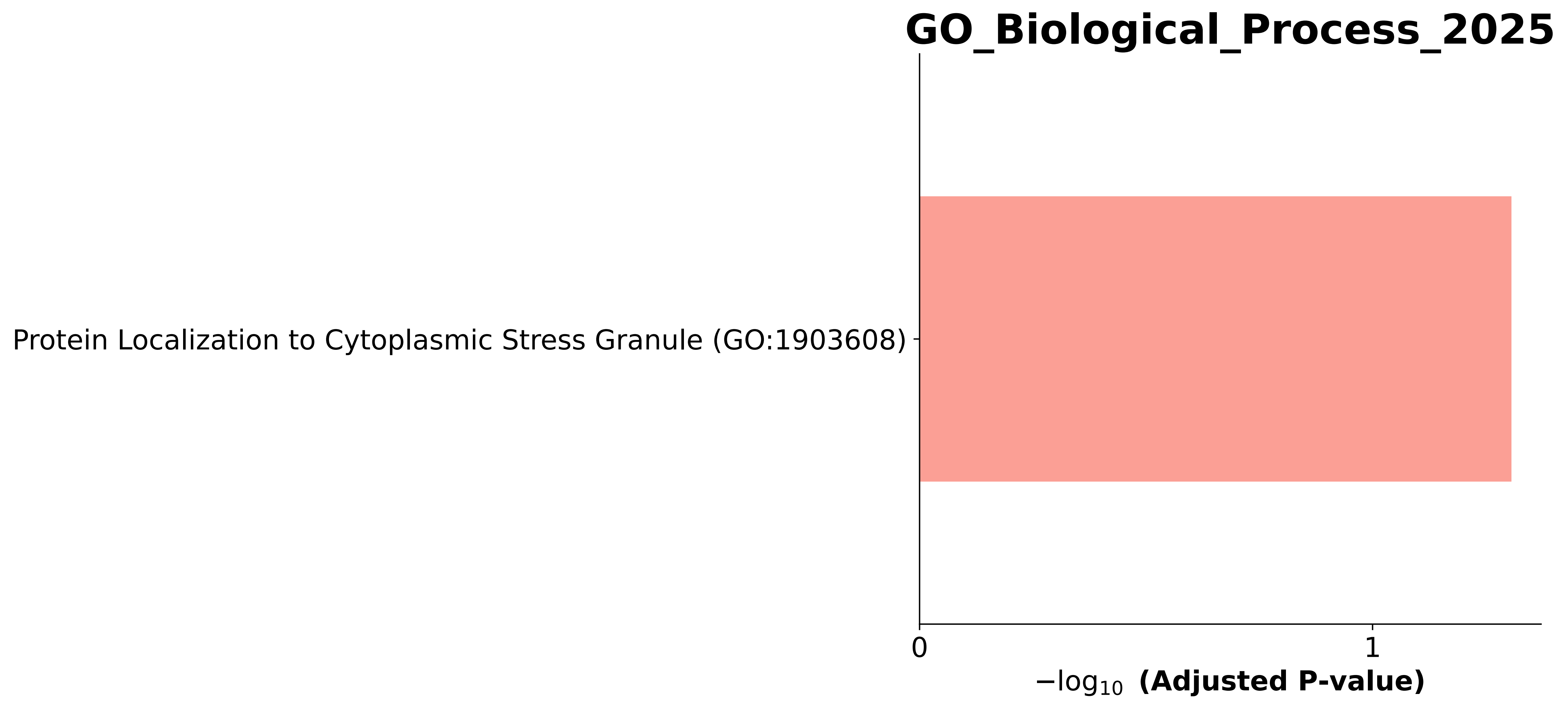

### GO_Biological_Process_2025.Mouse.enrichr.reports.pdf

#### GO_Biological_Process_2025

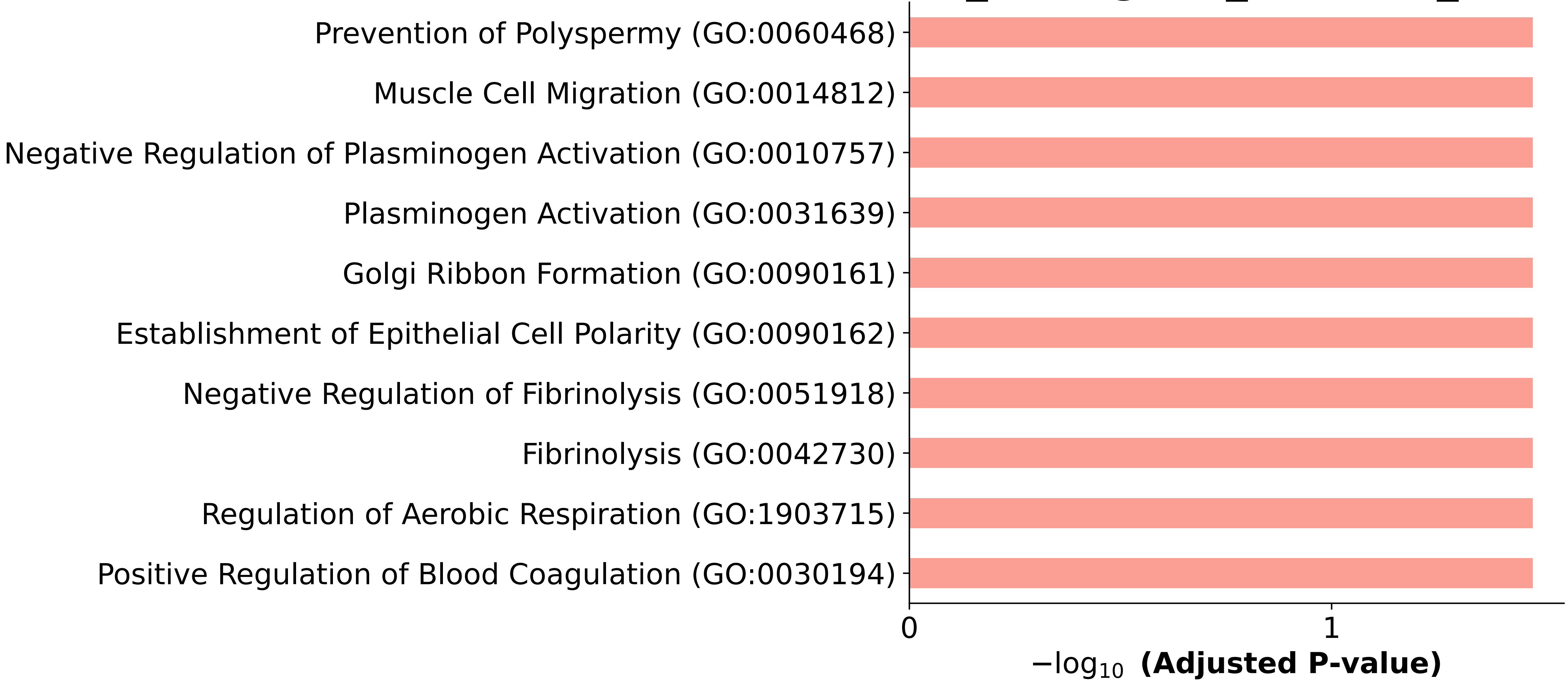

### GO_Biological_Process_2025.Mouse.enrichr.reports.pdf

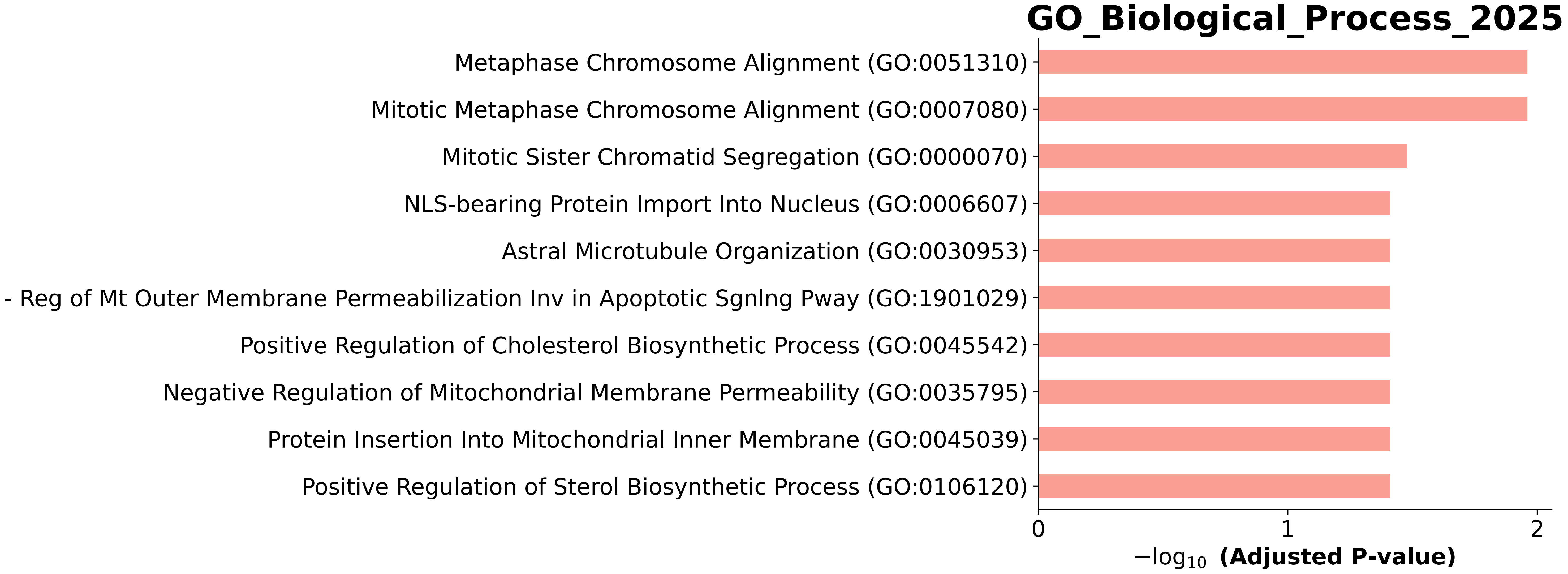

### GO_Biological_Process_2025.Mouse.enrichr.reports.pdf

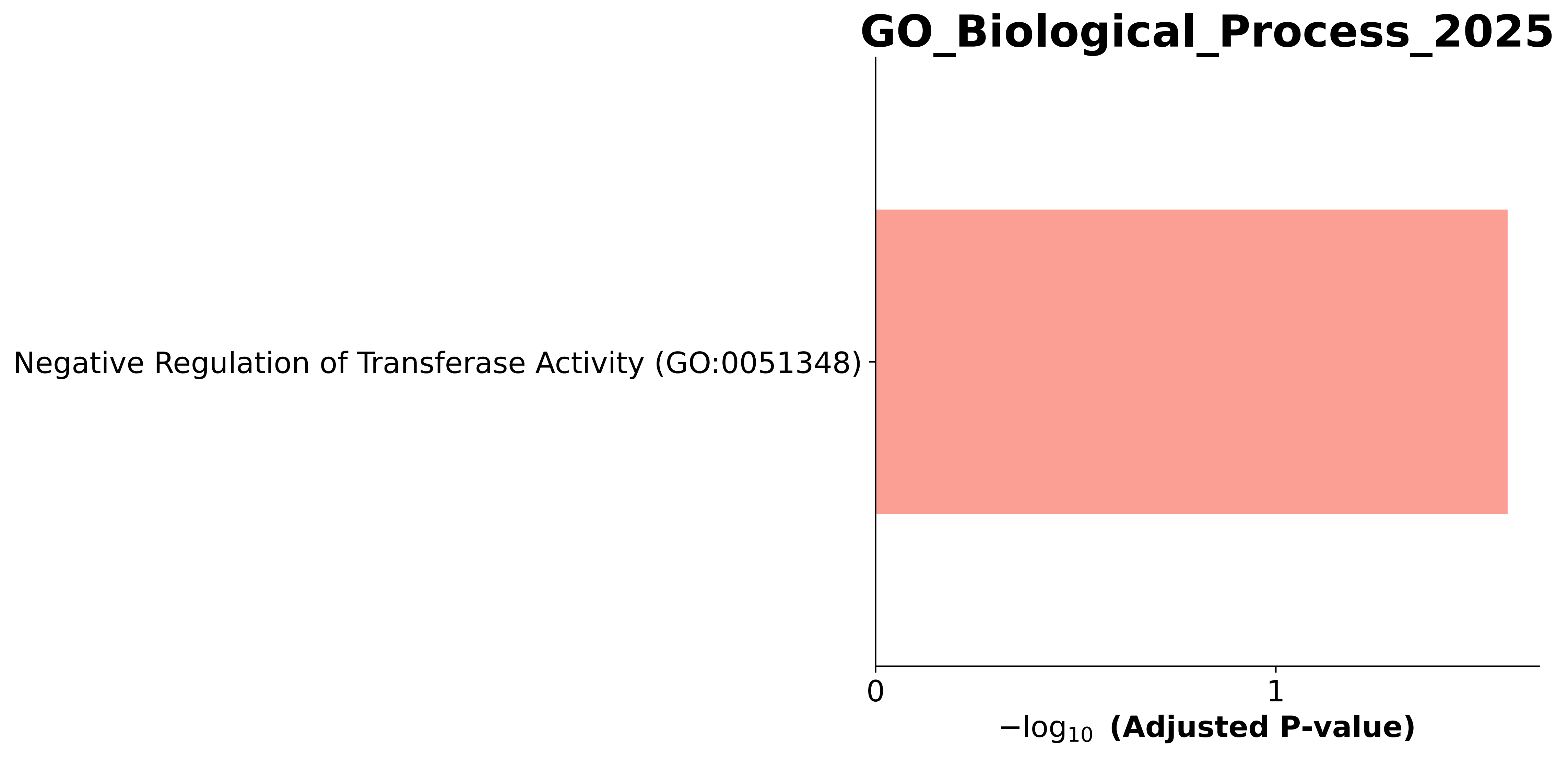

### GO_Biological_Process_2025.Mouse.enrichr.reports.pdf

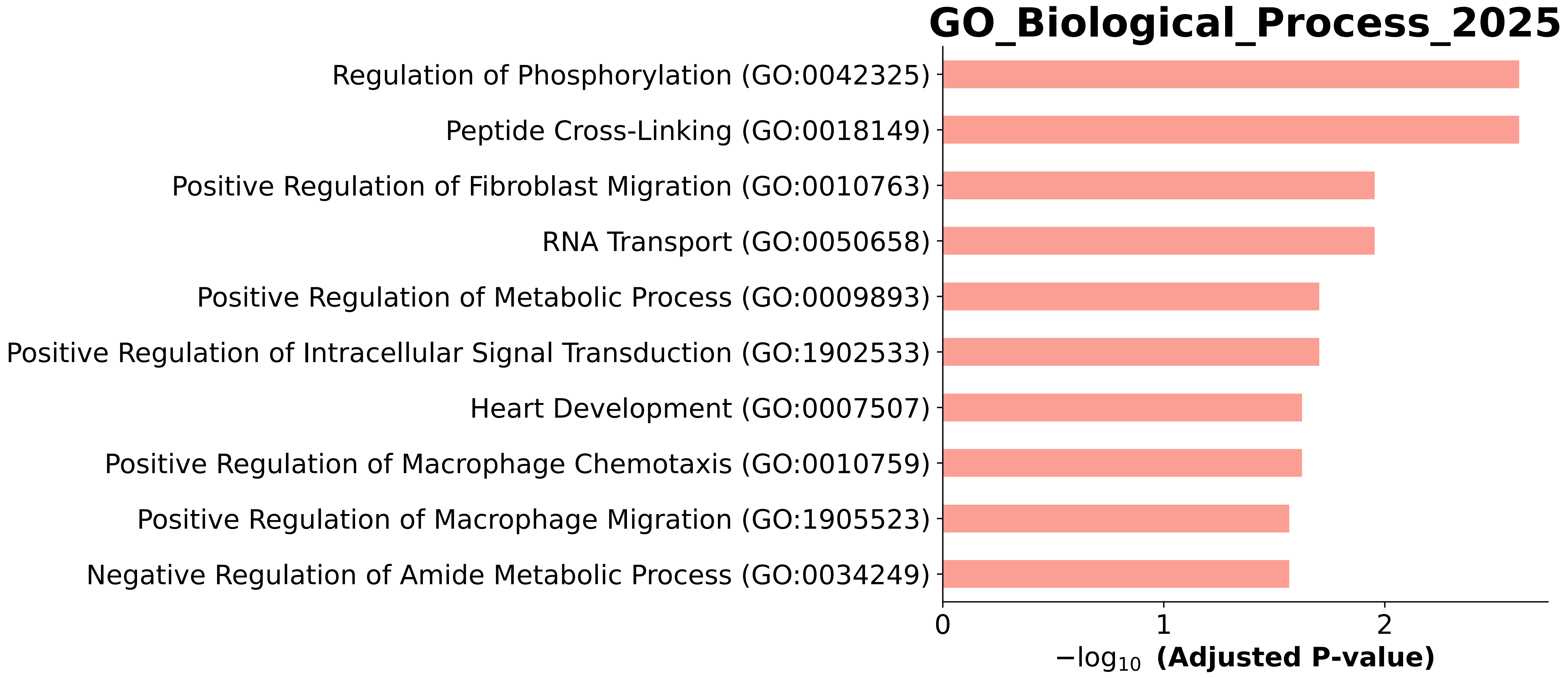

### GO_Biological_Process_2025.Mouse.enrichr.reports.pdf

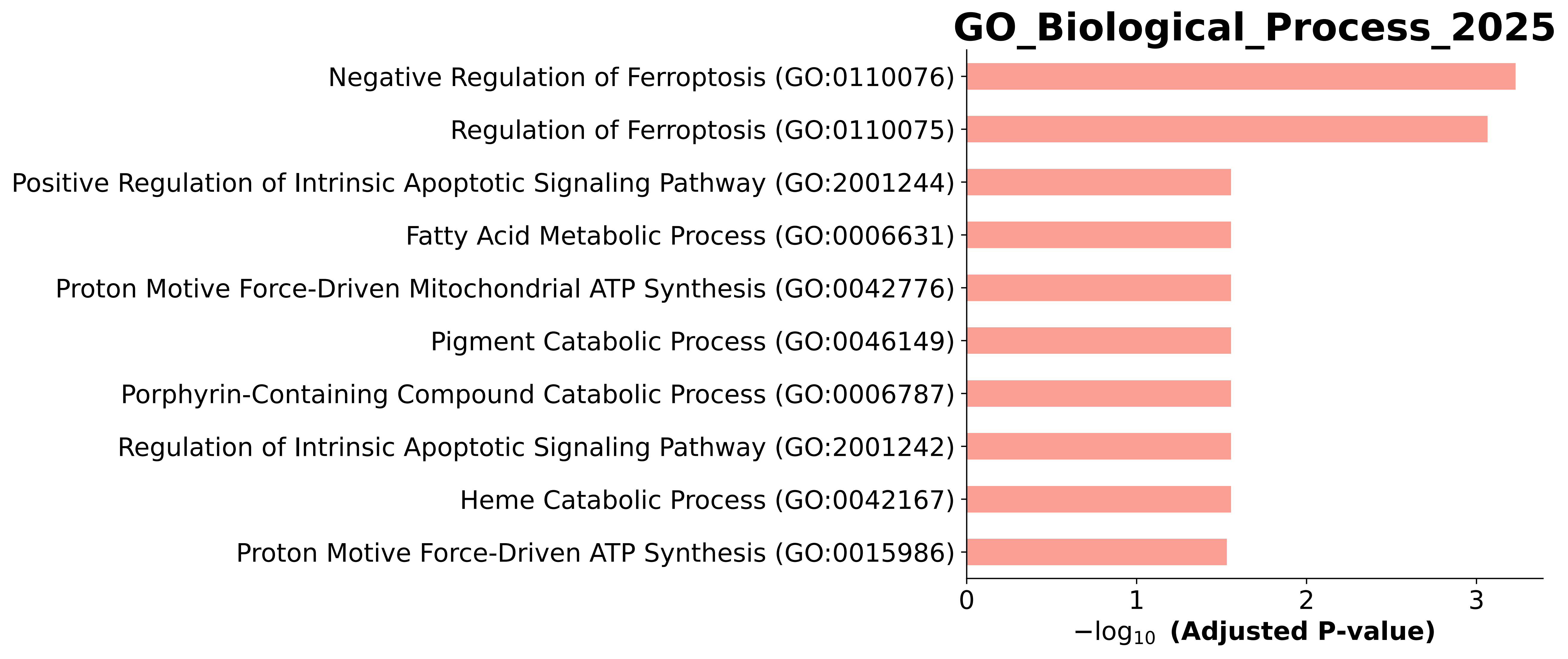

### GO_Biological_Process_2025.Mouse.enrichr.reports.pdf

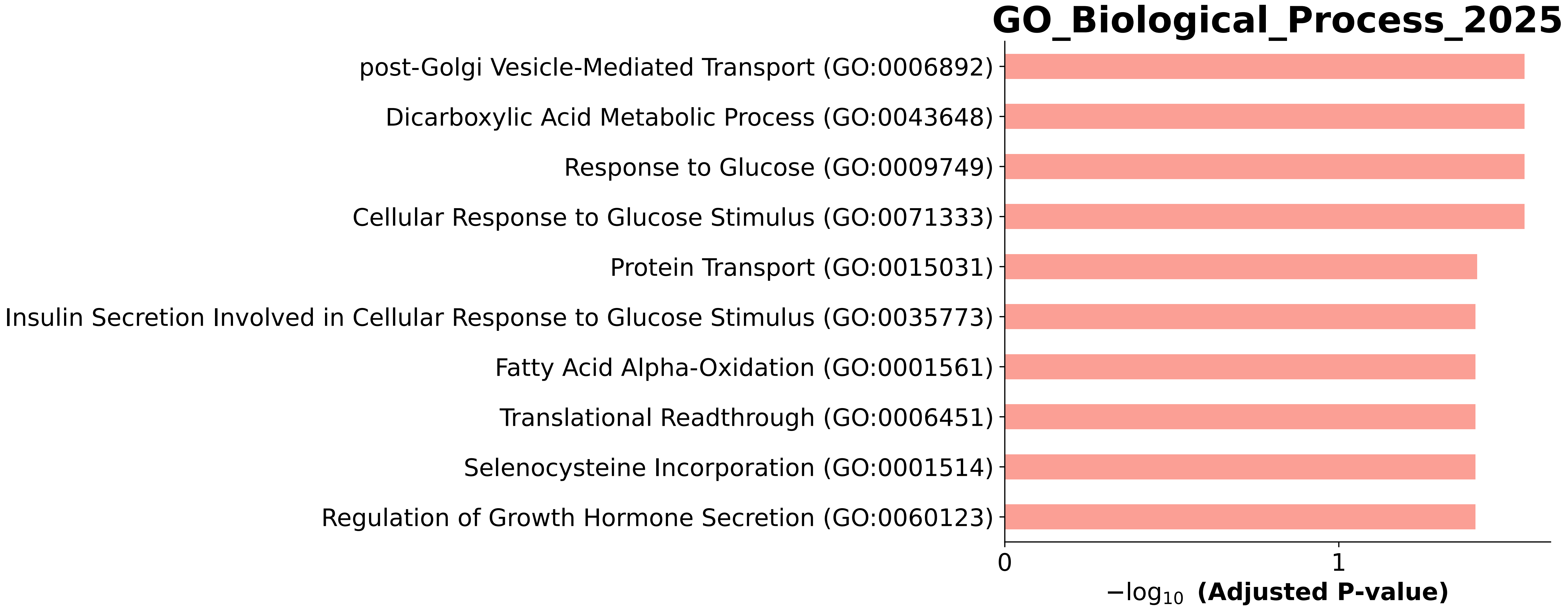

### GO_Biological_Process_2025.Mouse.enrichr.reports.pdf

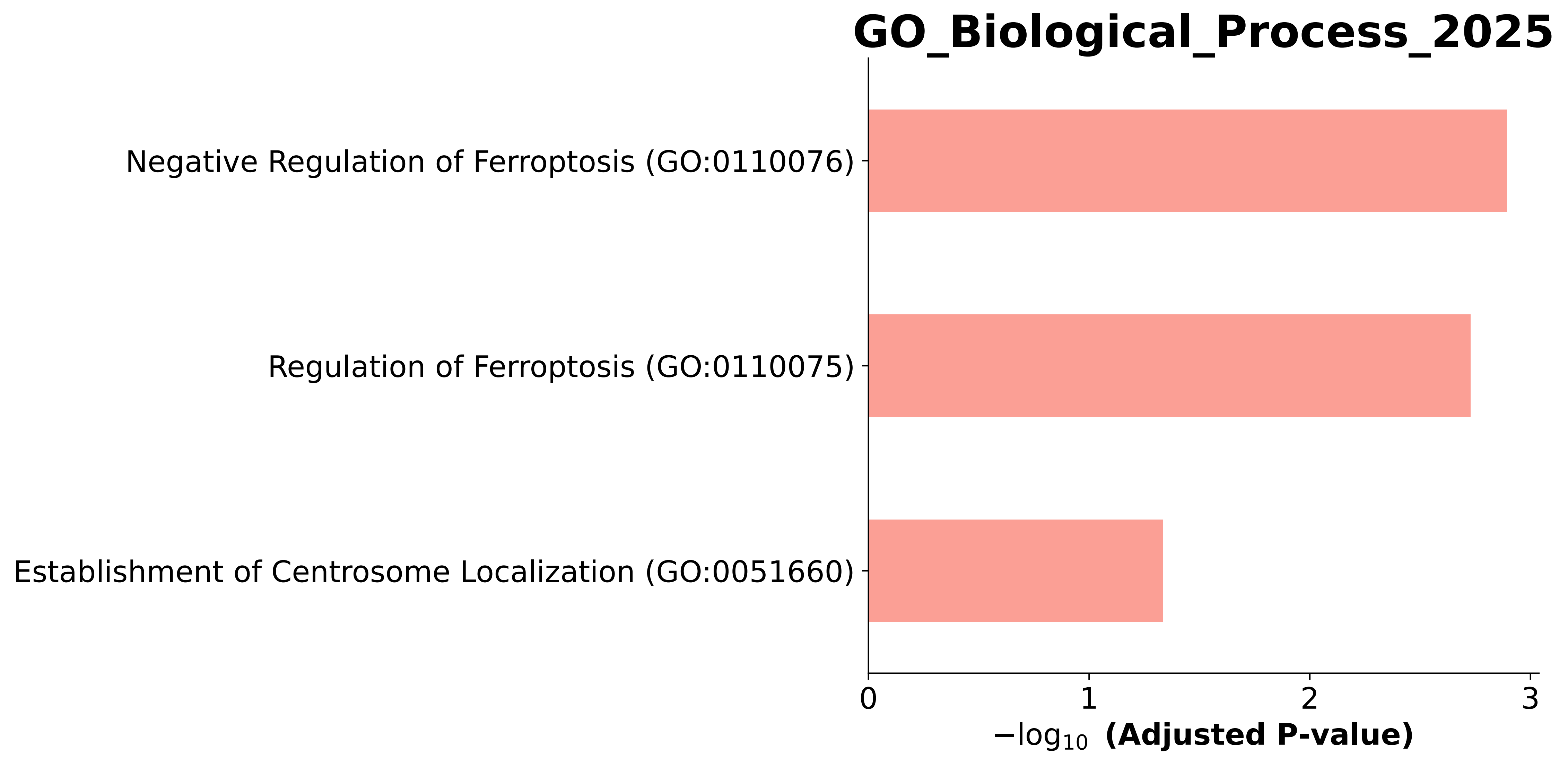

### GO_Biological_Process_2025.Mouse.enrichr.reports.pdf

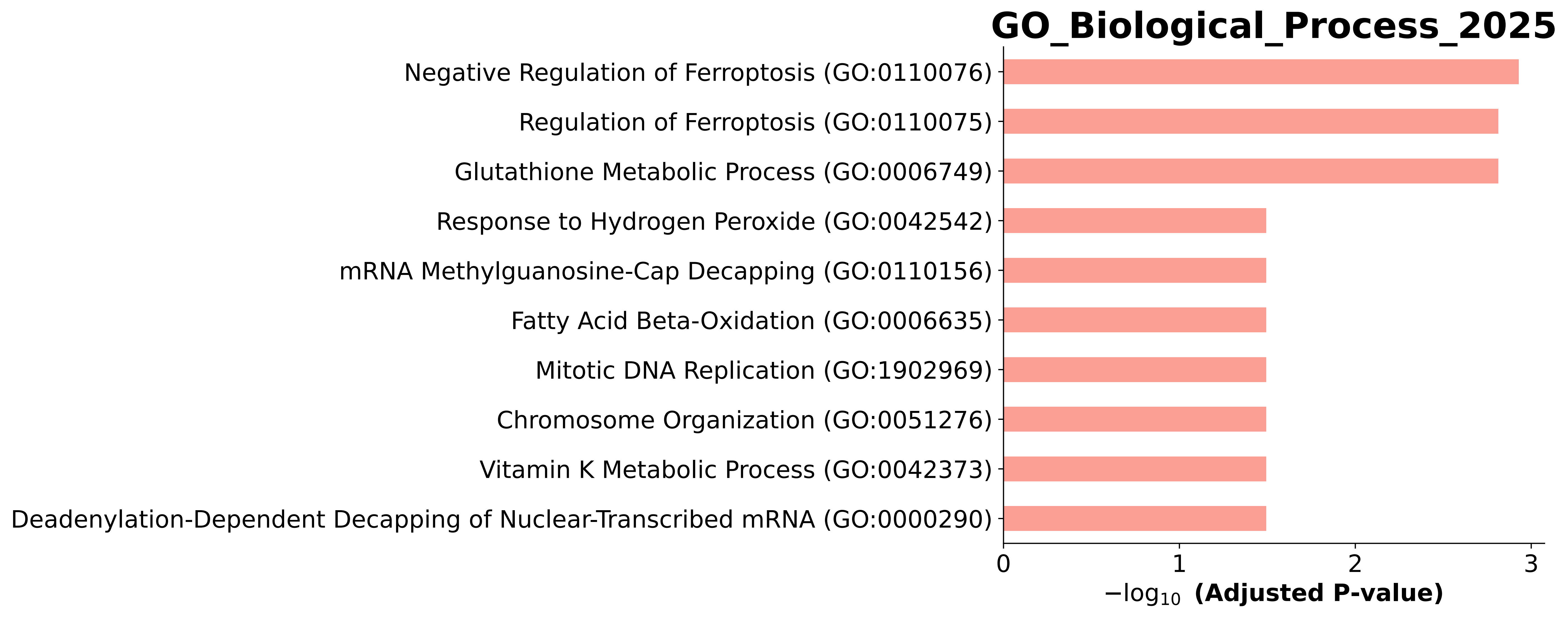

### GO_Biological_Process_2025.Mouse.enrichr.reports.pdf

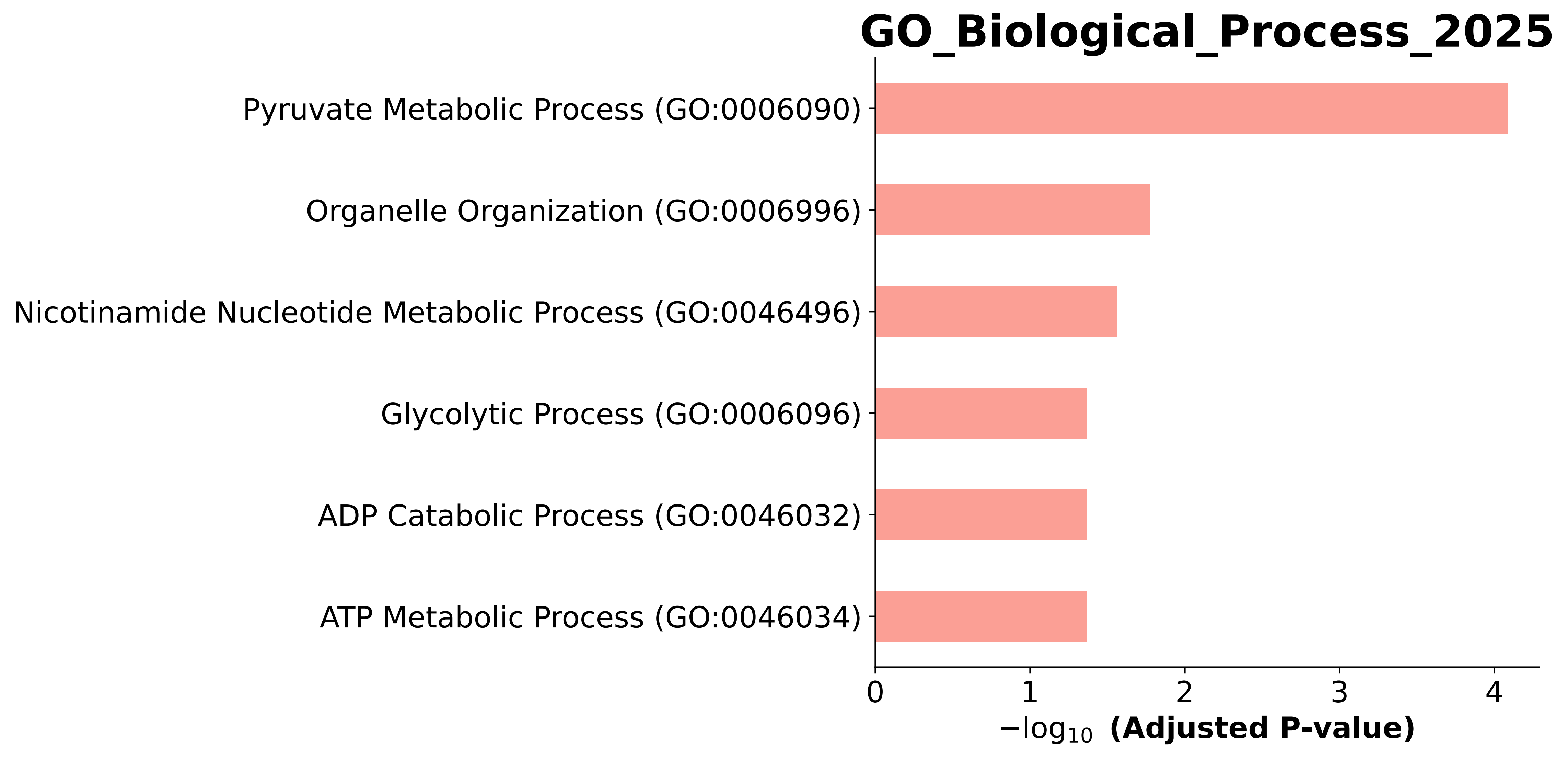

### GO_Biological_Process_2025.Mouse.enrichr.reports.pdf

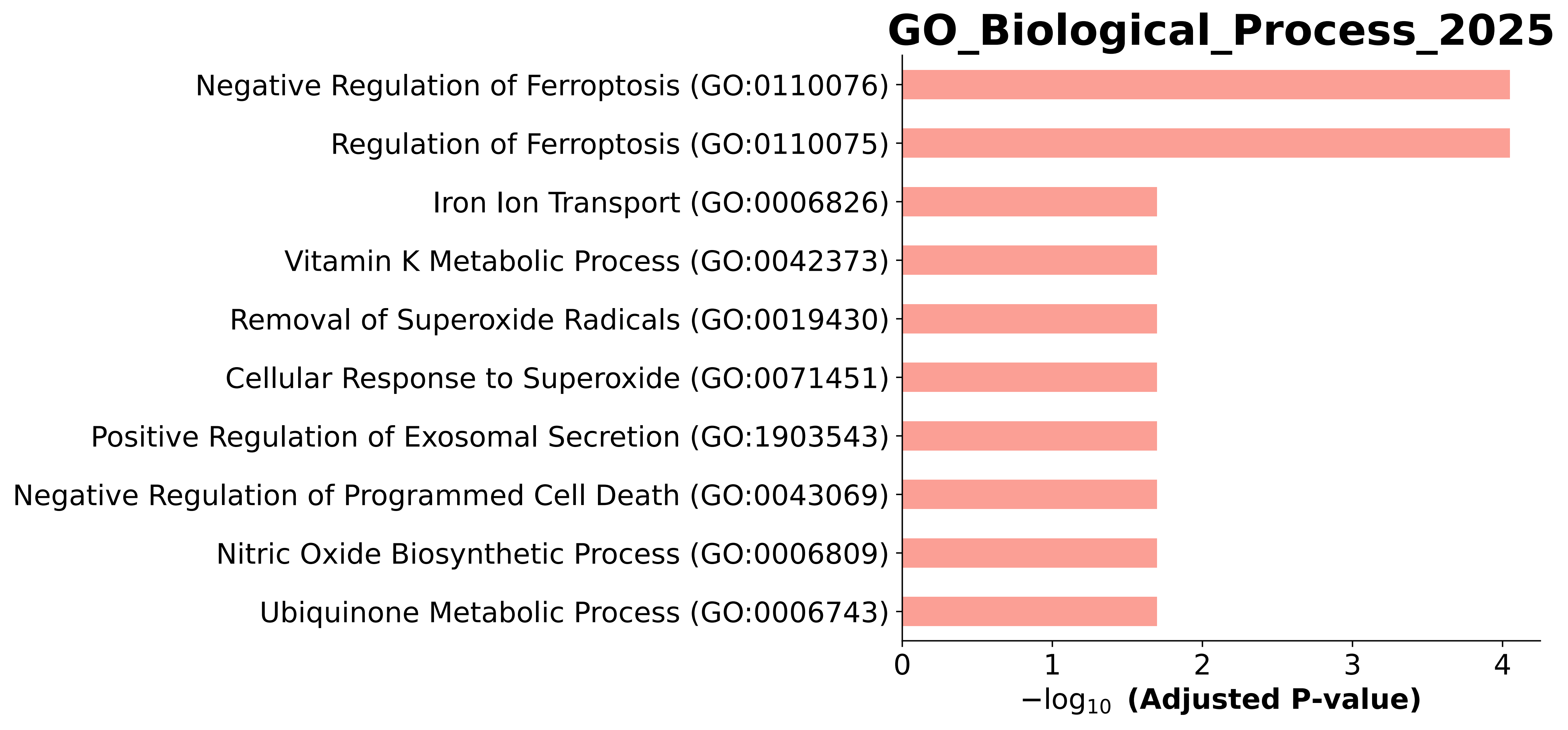

### GO_Biological_Process_2025.Mouse.enrichr.reports.pdf

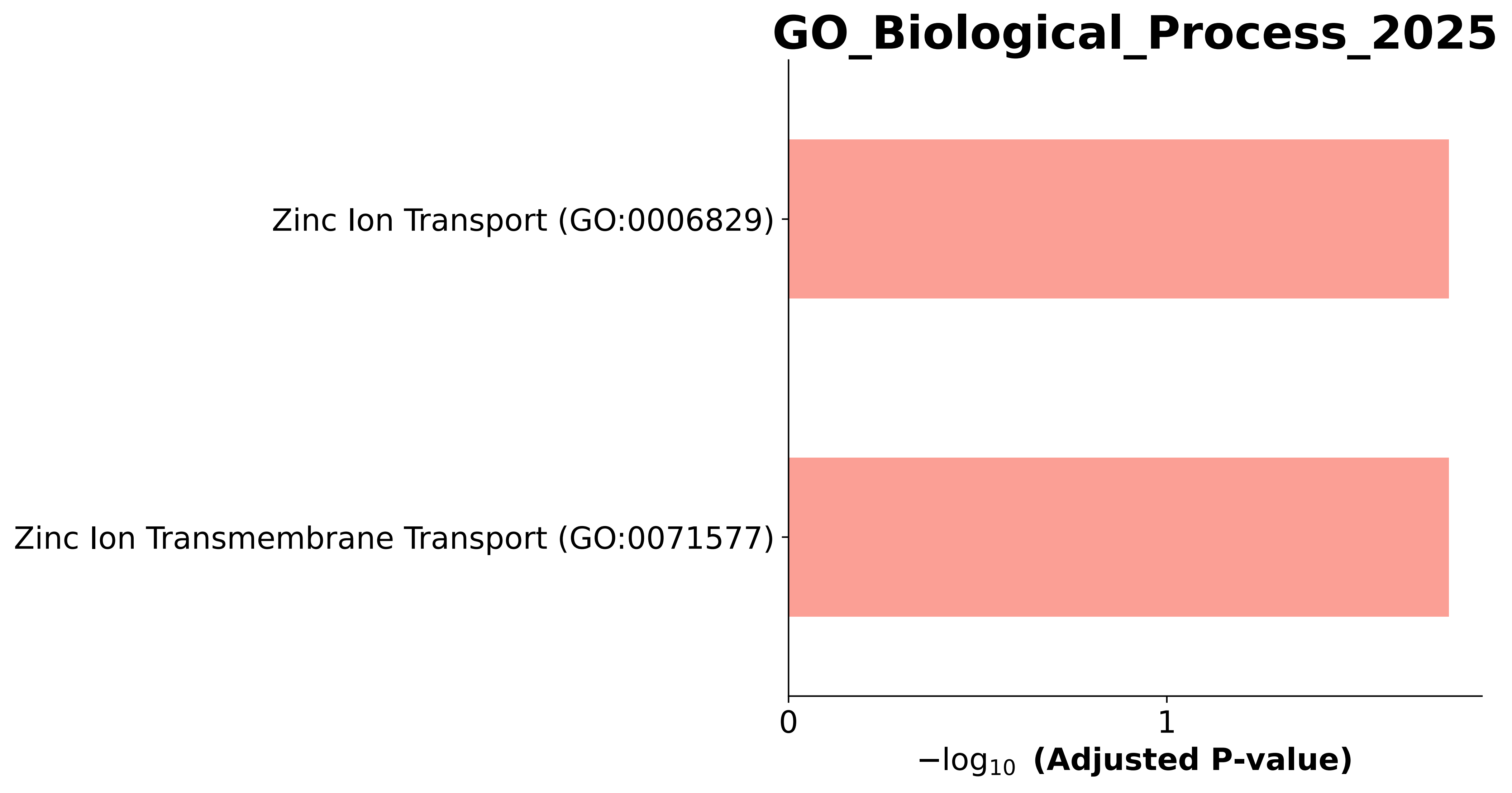

### GO_Biological_Process_2025.Mouse.enrichr.reports.pdf

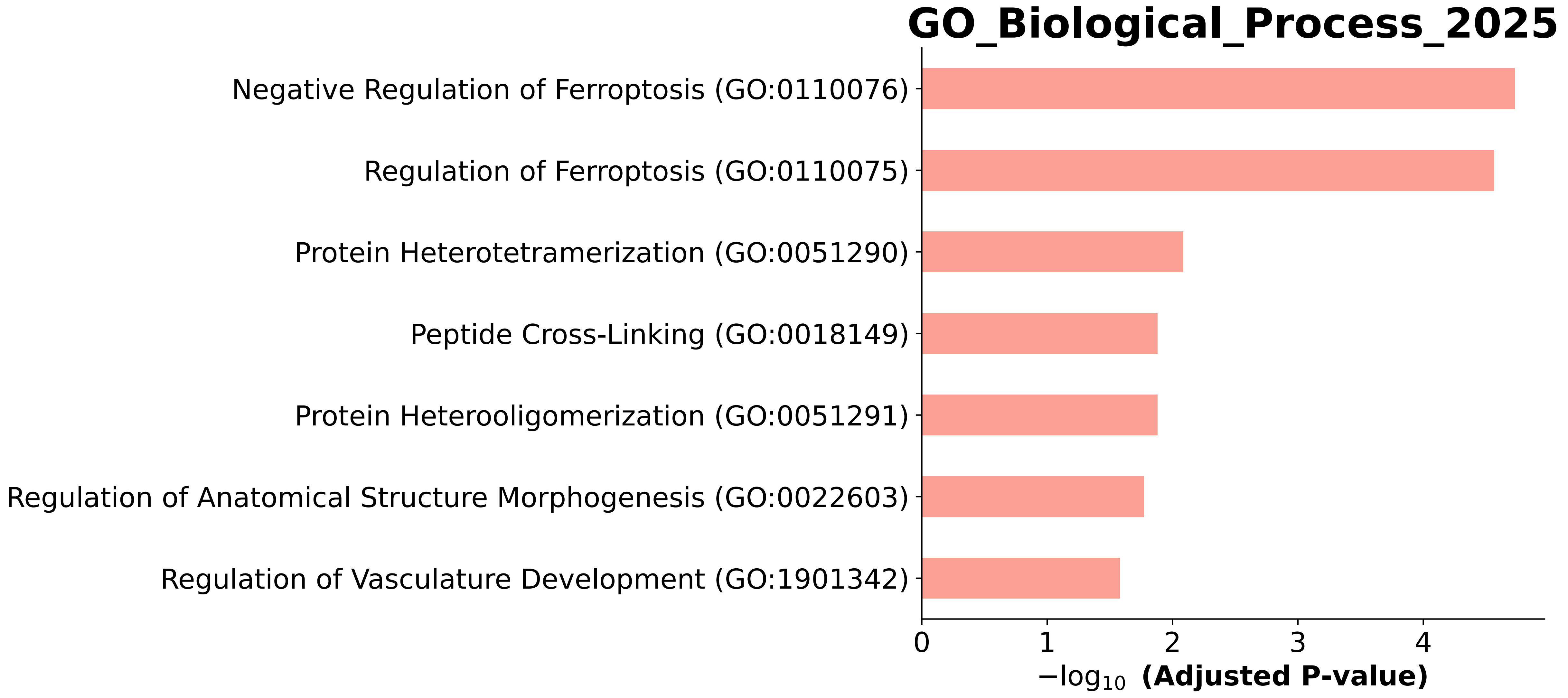

### GO_Biological_Process_2025.Mouse.enrichr.reports.pdf

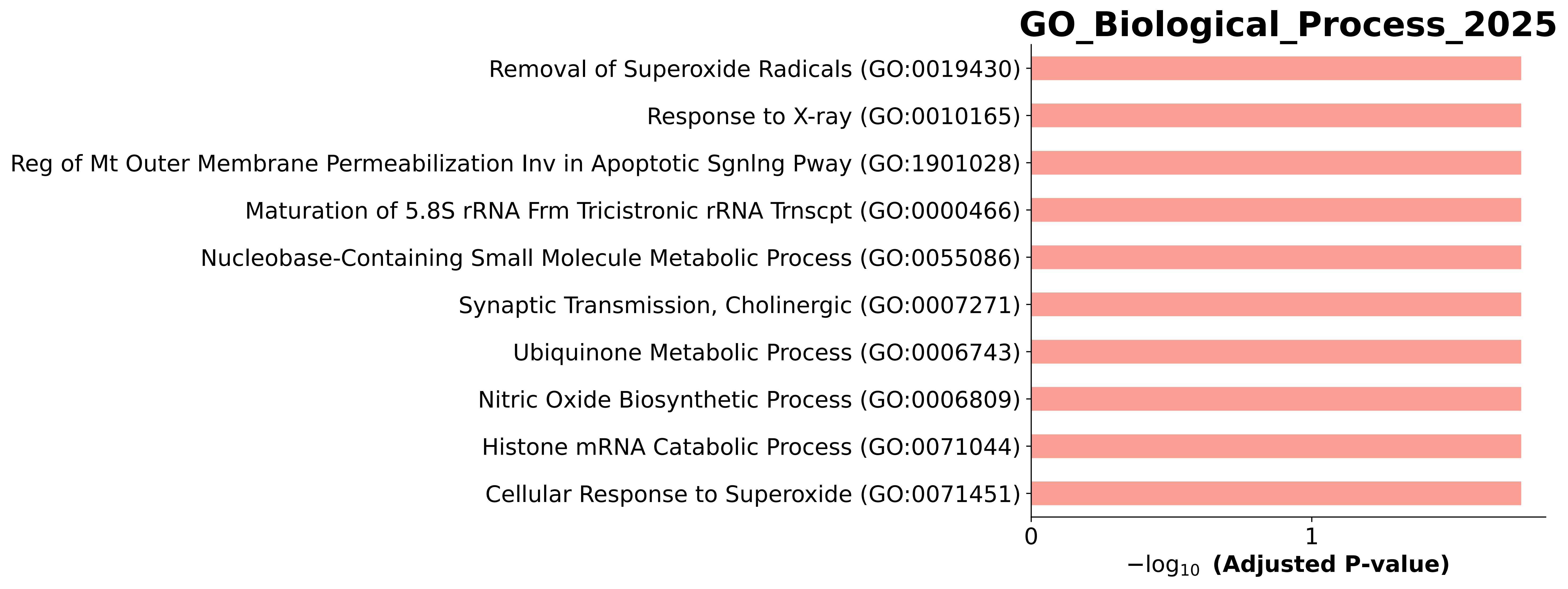

### GO_Biological_Process_2025.Mouse.enrichr.reports.pdf

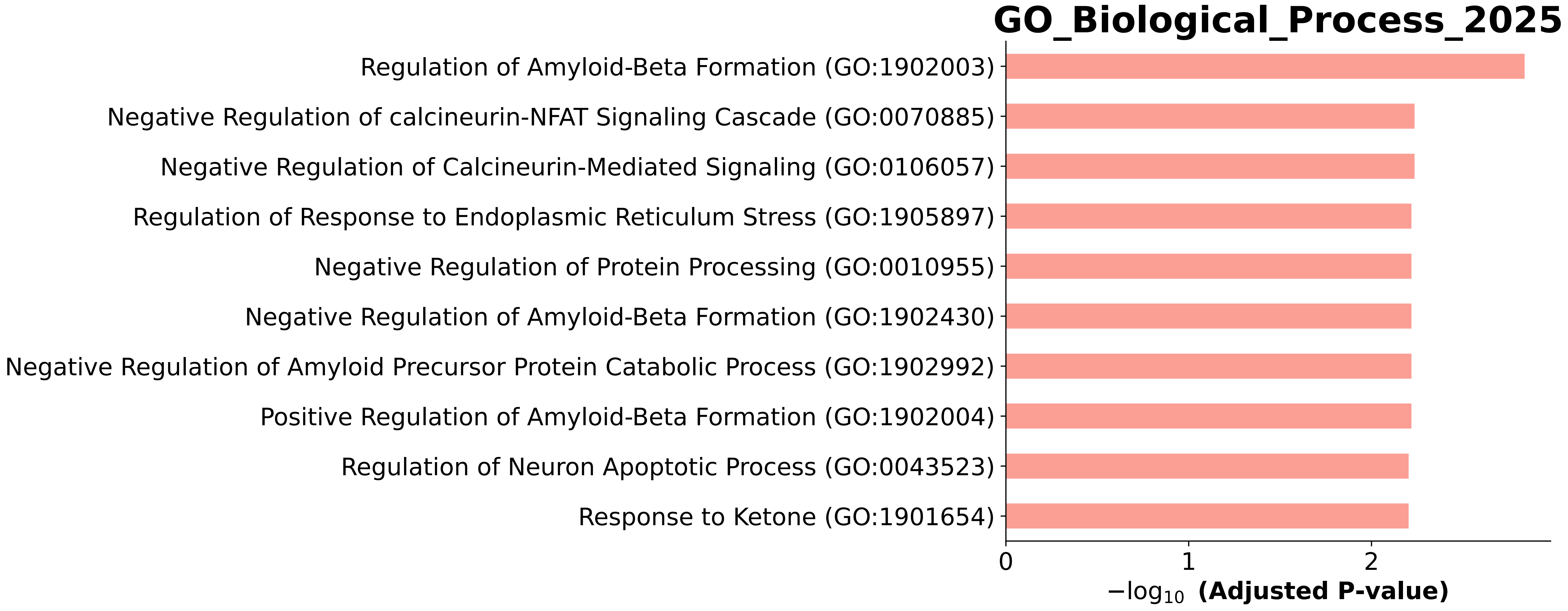

### GO_Biological_Process_2025.Mouse.enrichr.reports.pdf

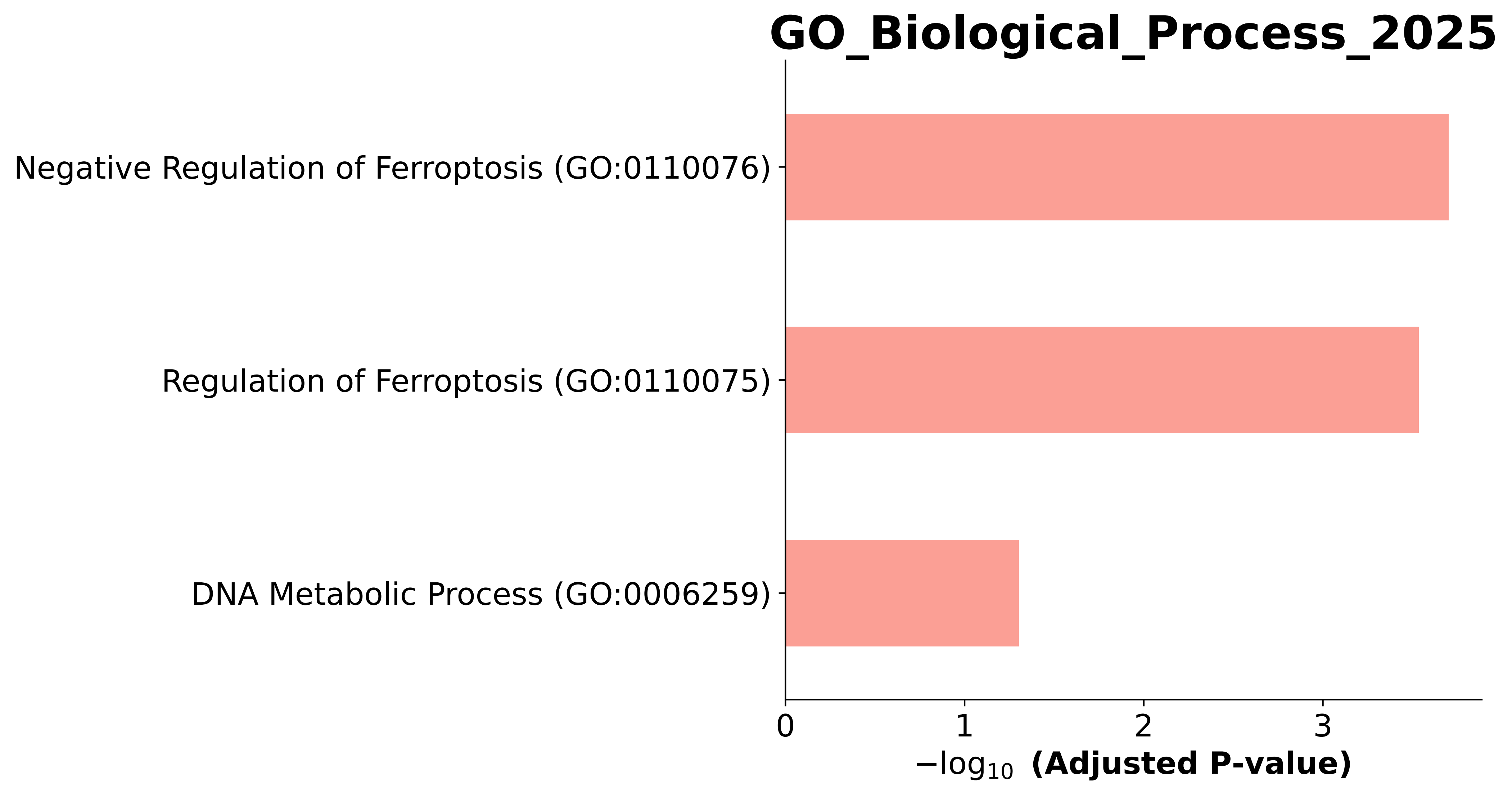

### GO_Biological_Process_2025.Mouse.enrichr.reports.pdf

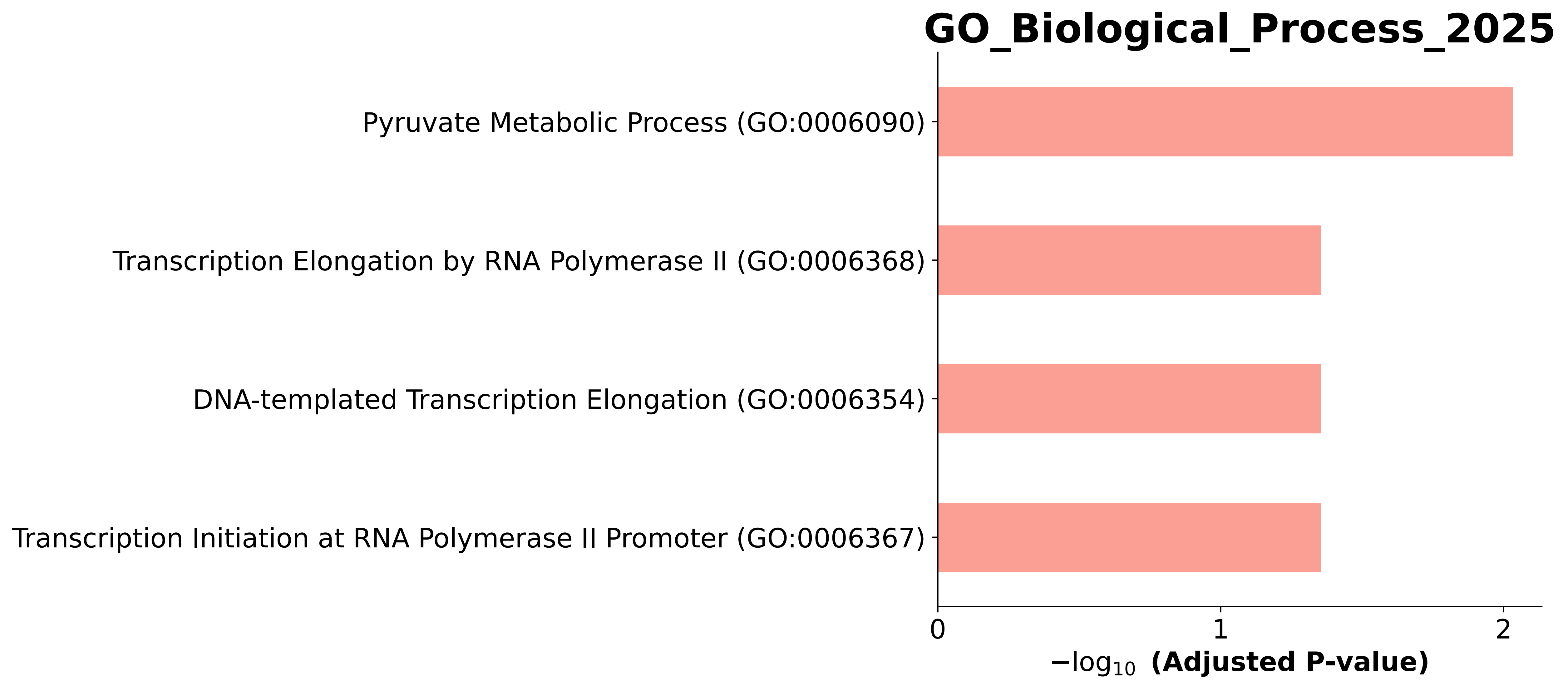

### GO_Biological_Process_2025.Mouse.enrichr.reports.pdf

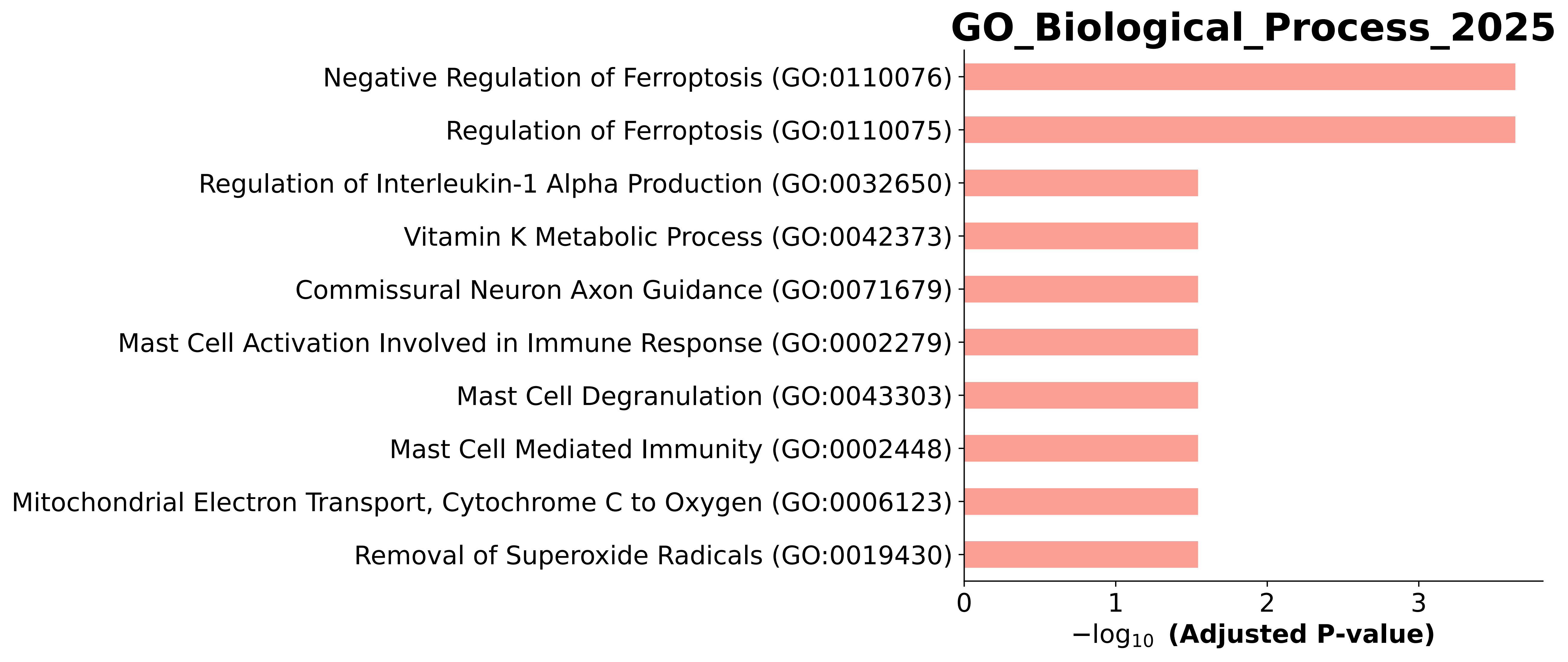

### GO_Biological_Process_2025.Mouse.enrichr.reports.pdf

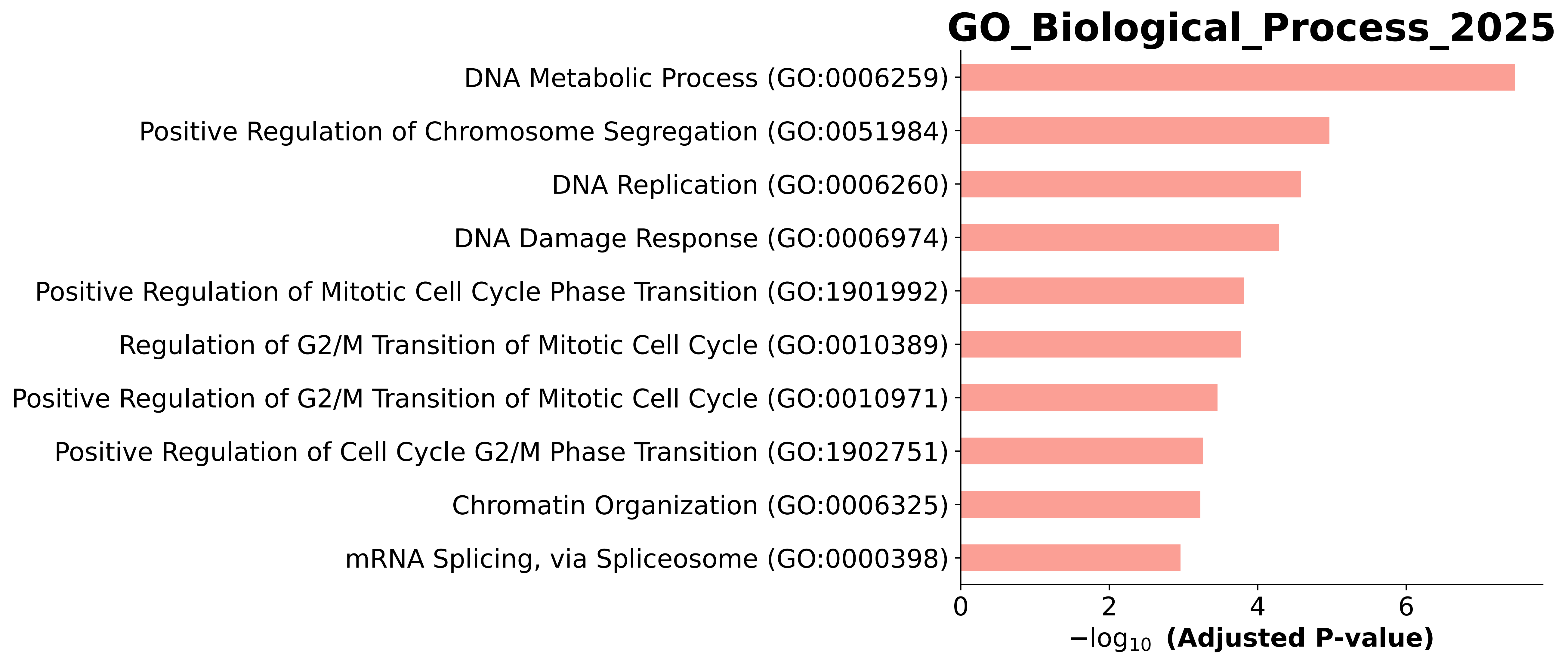

### GO_Biological_Process_2025.Mouse.enrichr.reports.pdf

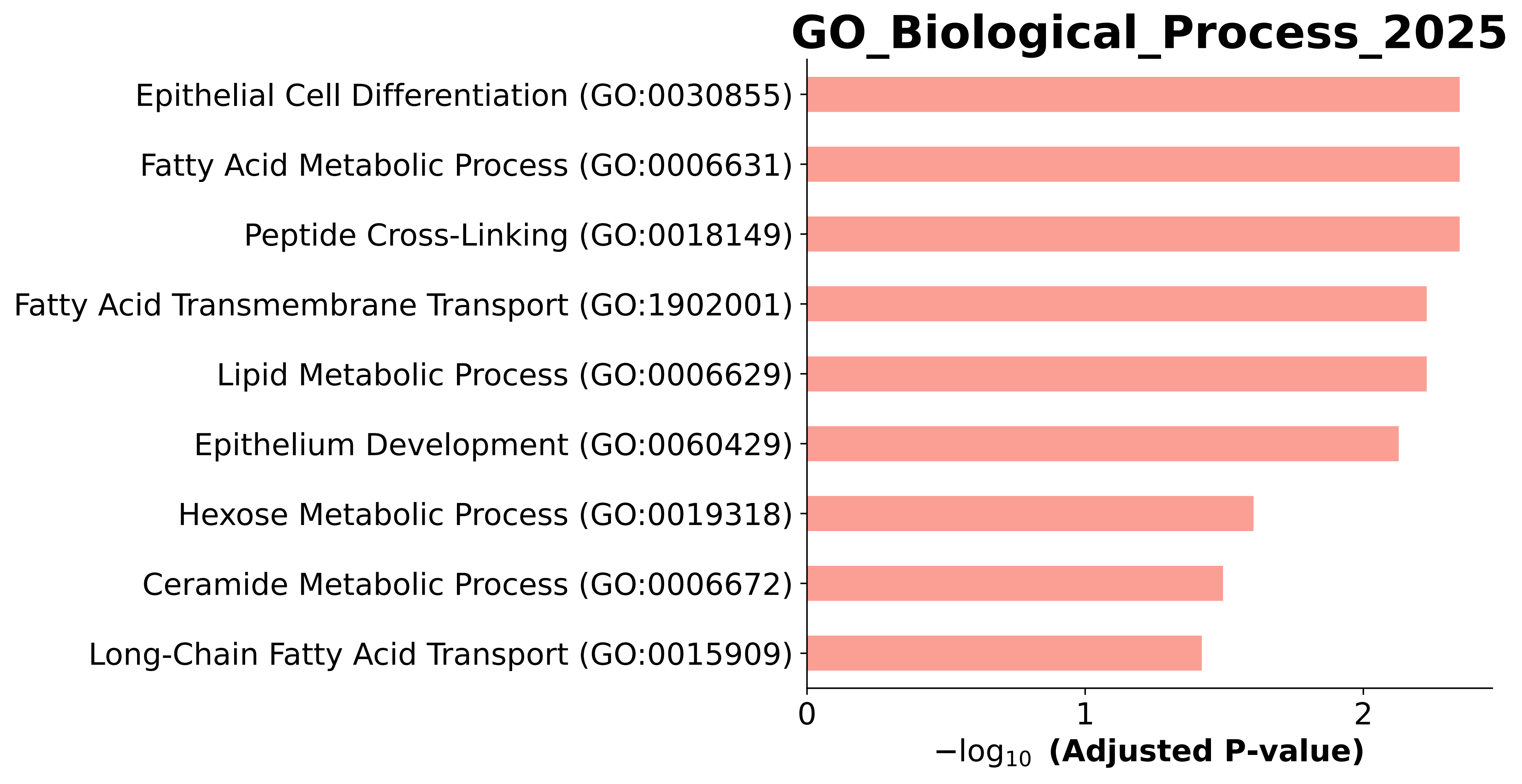

### GO_Biological_Process_2025.Mouse.enrichr.reports.pdf

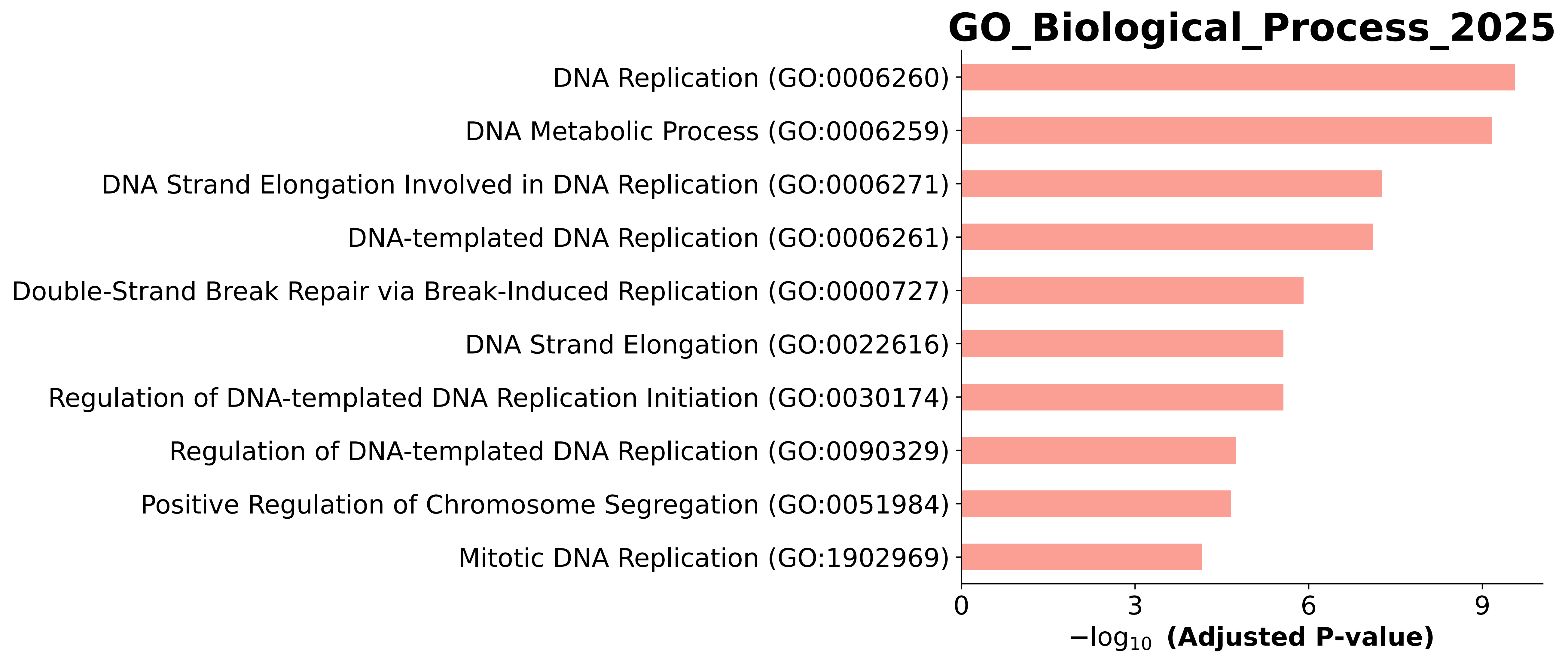

### GO_Biological_Process_2025.Mouse.enrichr.reports.pdf

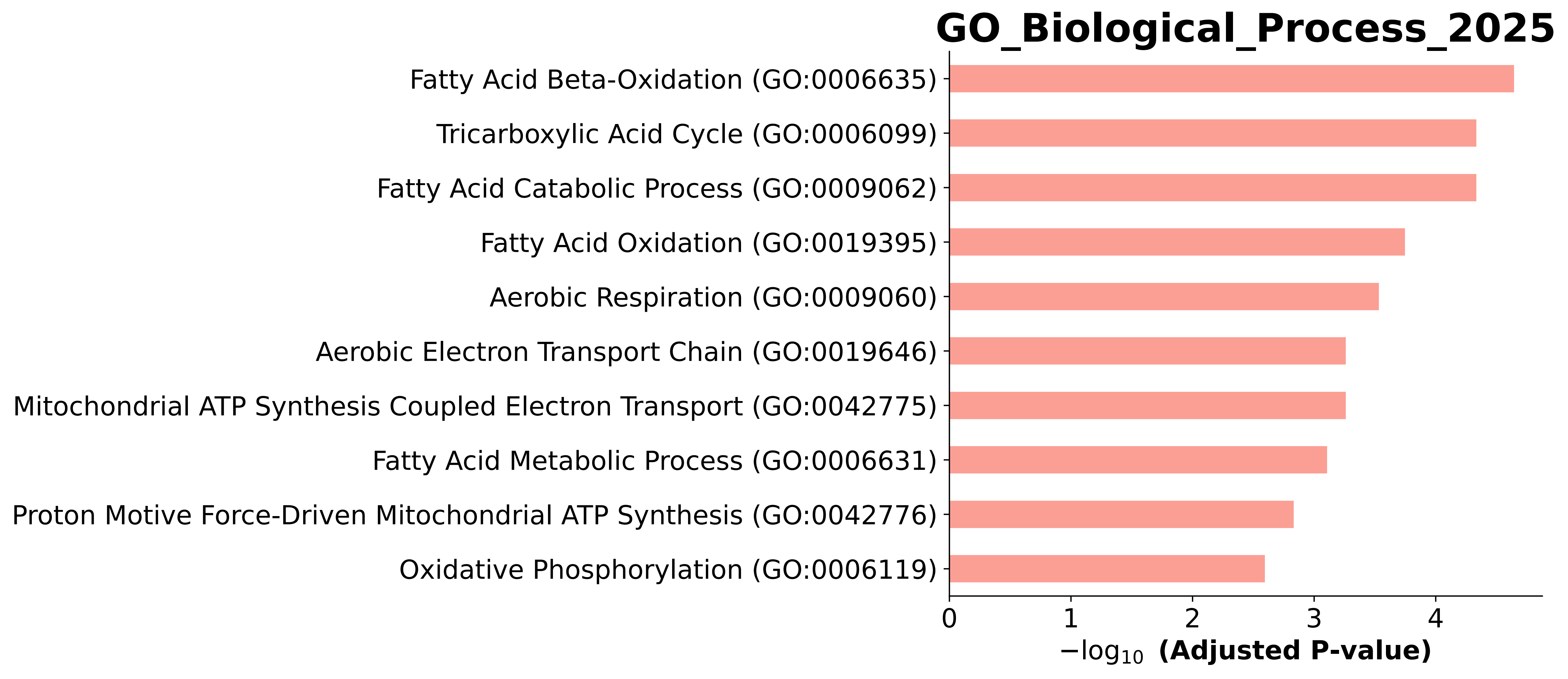

### GO_Biological_Process_2025.Mouse.enrichr.reports.pdf

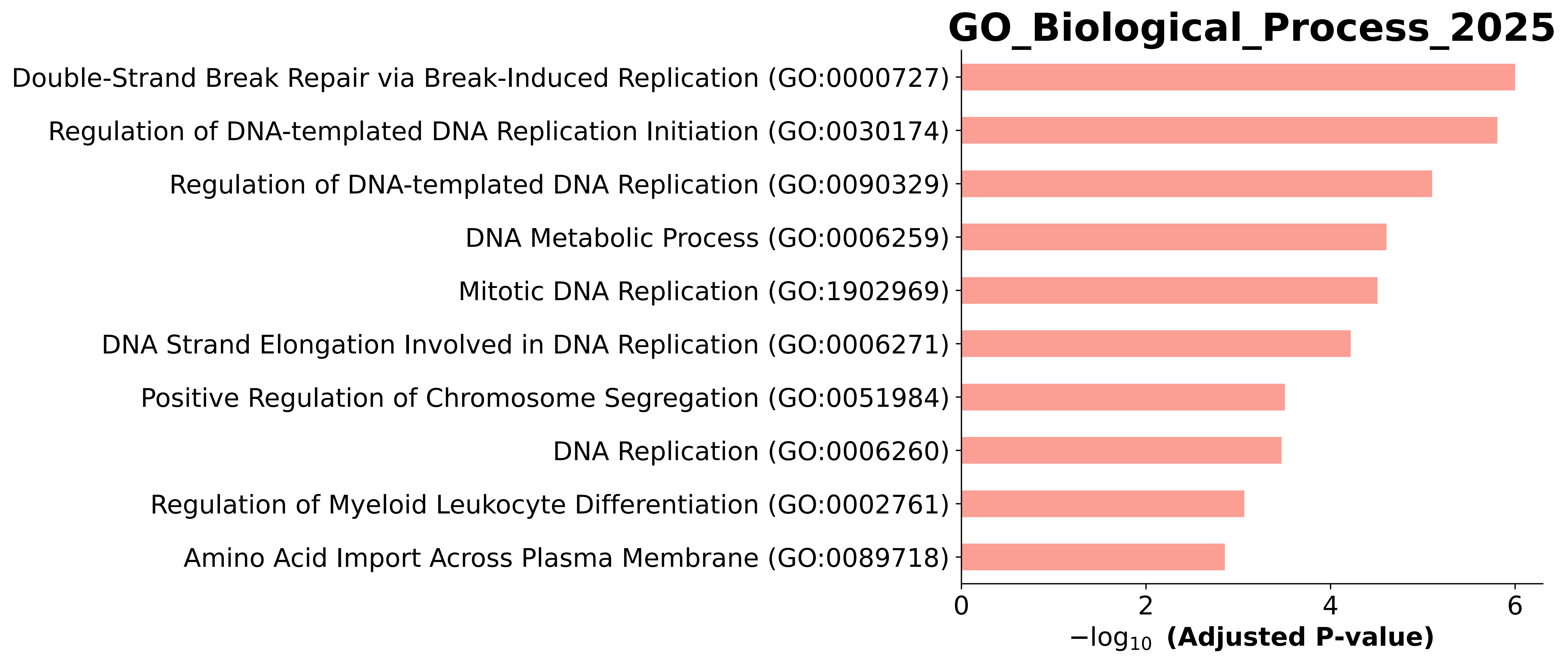

### GO_Biological_Process_2025.Mouse.enrichr.reports.pdf

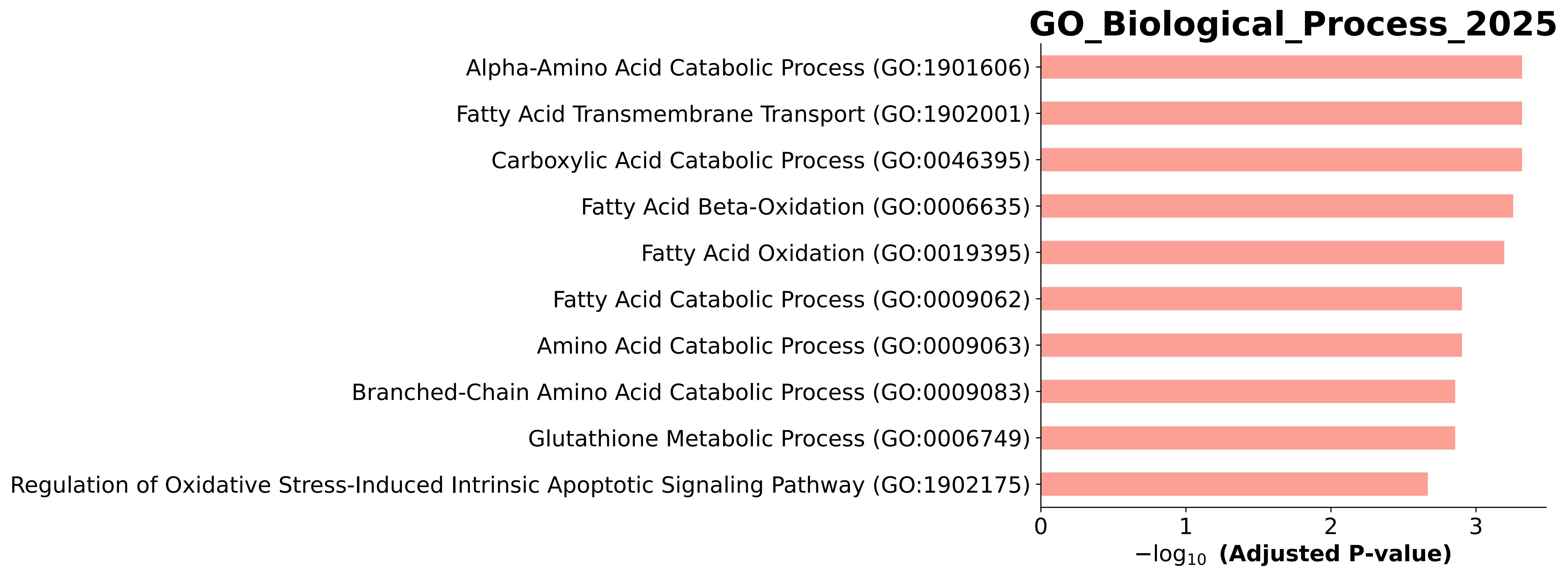

### GO_Biological_Process_2025.Mouse.enrichr.reports.pdf

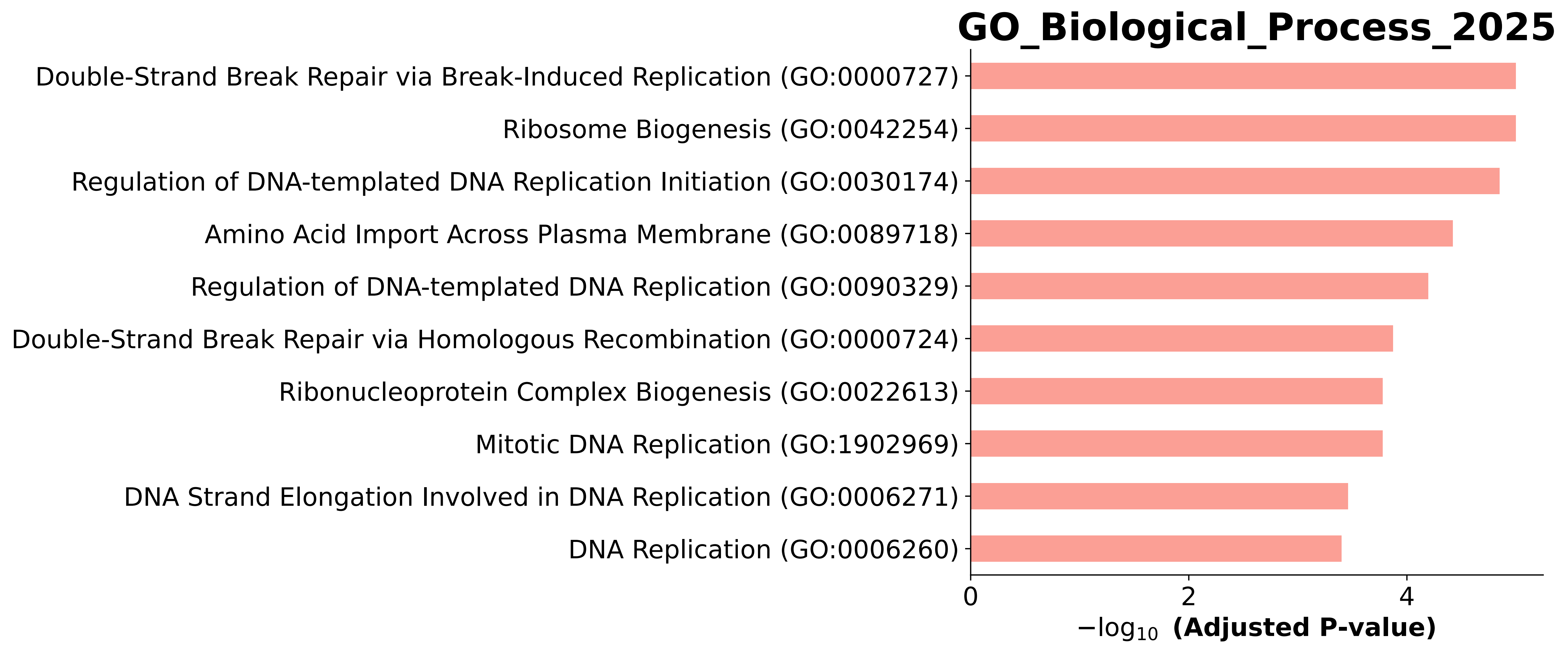

### GO_Biological_Process_2025.Mouse.enrichr.reports.pdf

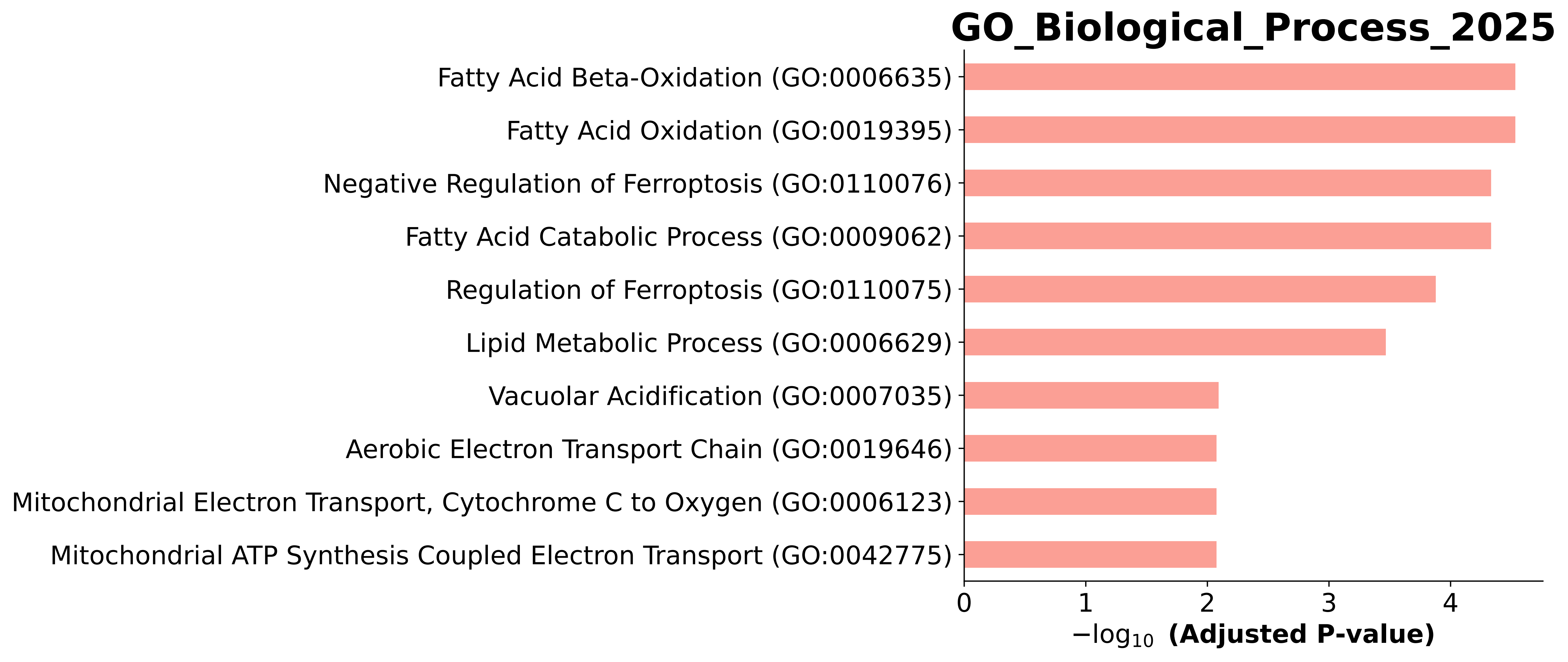

### GO_Biological_Process_2025.Mouse.enrichr.reports.pdf

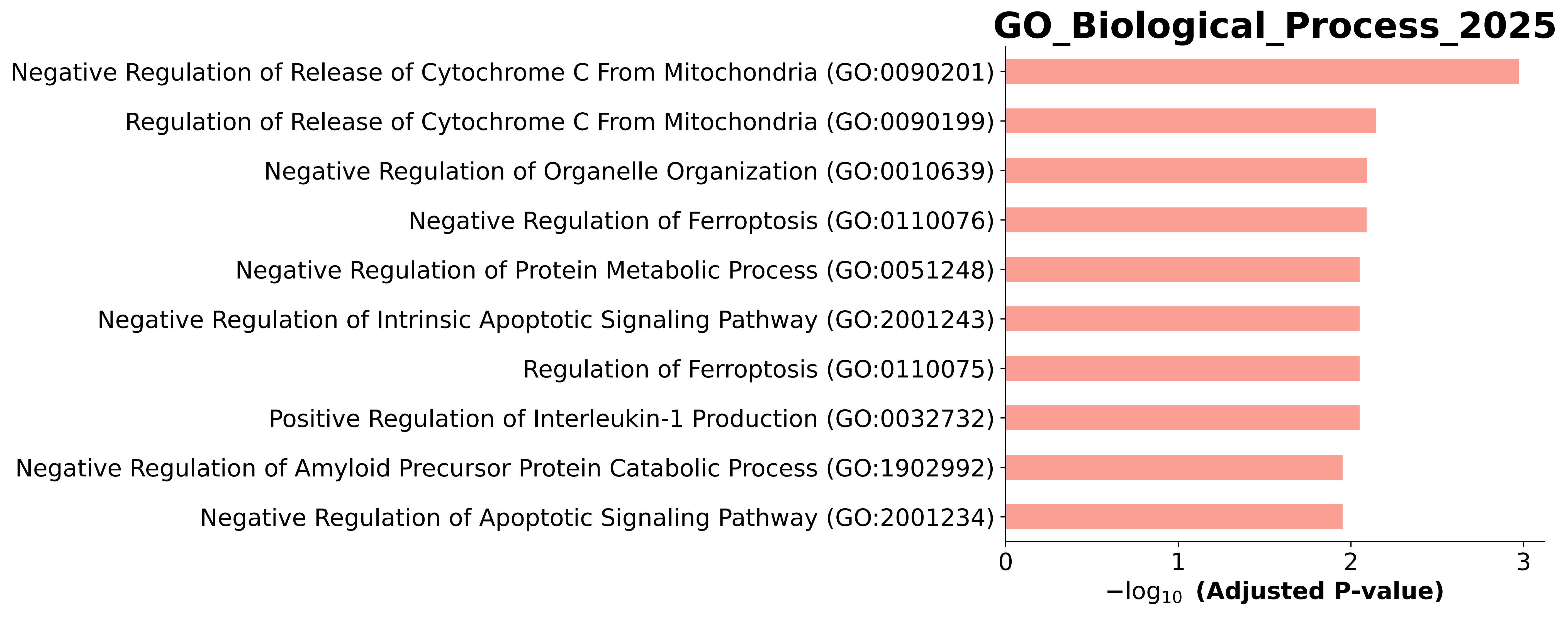
