## supplementary information and figures for "Fatty Acid β-oxidation and Ferroptosis Define a Survival-Productivity Trade Off in CHO fed-Batch Bioreactors"

**Supplementary Methods**

| Time (min) | A% | B% | Flowrate (uL/min) |
|---|---|---|---|
| 0 | 97 | 3 | 2 |
| 3 | 97 | 3 | 2 |
| 32.5 | 75 | 25 | 2 |
| 40 | 62.5 | 40 | 2 |
| 41 | 5 | 95 | 2 |
| 43 | 5 | 95 | 2 |
| 43.5 | 97 | 3 | 2 |
| 50 | 97 | 3 | 2 |
| 51 | 5 | 95 | 3 |
| 52 | 5 | 95 | 2 |
| 55 | 97 | 3 | 2 |
| 60 | 97 | 3 | 2 |

**Table S1**: LC gradient used on the Zeno ToF 7600, where A is 0.1% formic acid, and B is acetonitrile with 0.1% formic acid.

| Parameter | Value | Parameter | Value |
|---|---|---|---|
| Curtain Gas | 35 | TOF End Mass | 2050 |
| CAD gas | 7 | TOF MS Accumulation time | 0.1 s |
| Ion source gas 1 | 20 | Declustering potential | 80 V |
| Ion gas source 2 | 60 | Fragmentation mode | CID |
| Temperature | 225 °C | TOF MSMS start mass | 100 |
| Column Temperature | 50 °C | TOF MSMS end mass | 2000 |
| Spray Voltage | 5000 V | TOF MSMS Accumulation time | 0.018 s |
| TOF Start Mass | 400 | Total scan time | 2.394 s |

**Table S2** DIA instrument parameters for the ZenoTOF 7600

**Supplementary Data**

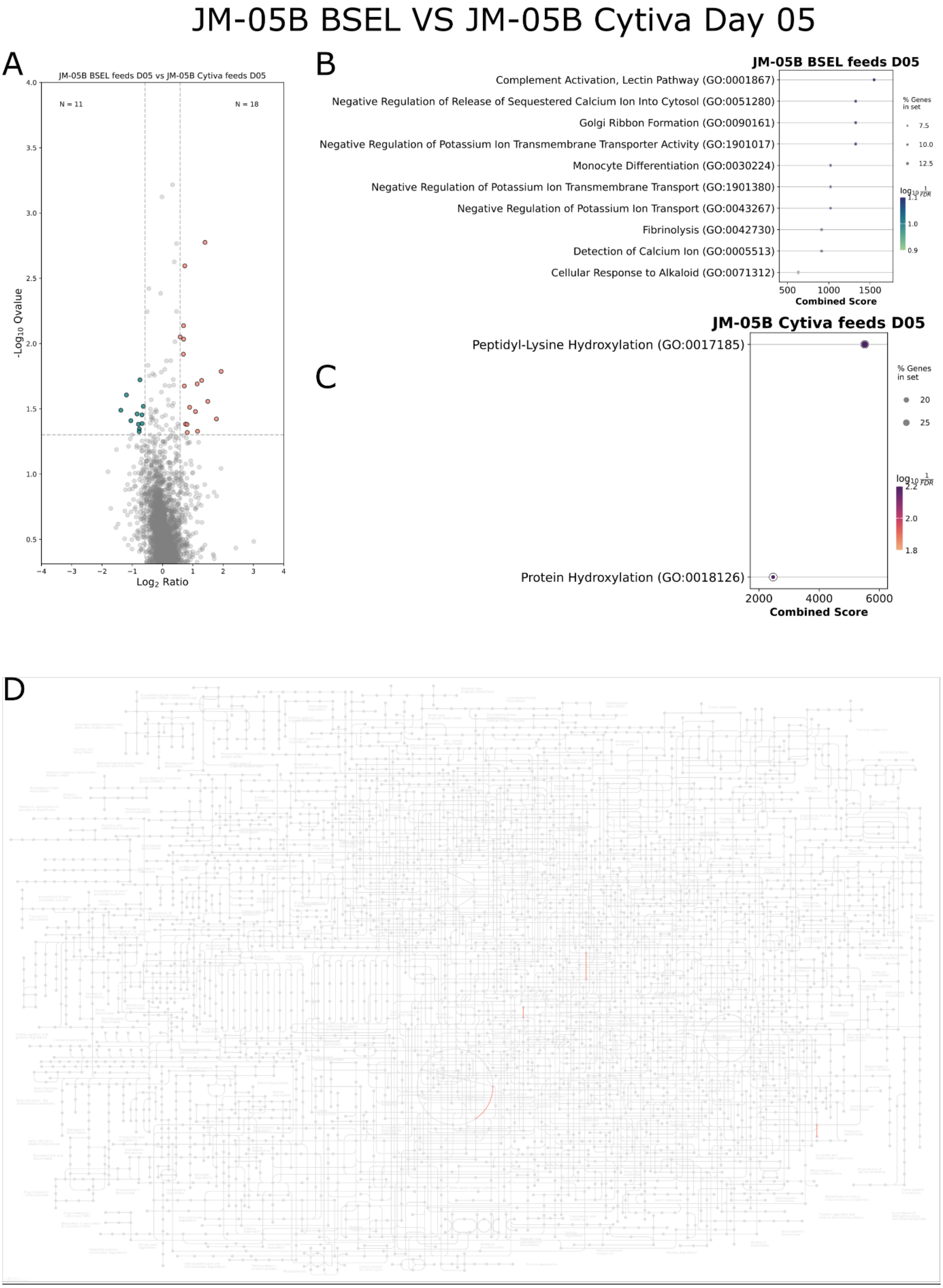


**Figure S1**: JM-05B BSEL (blue) vs JM-05B Cytiva (pink) D05 A) Volcano plot, B-C) gene set enrichment analysis D) KEGG mapped pathways

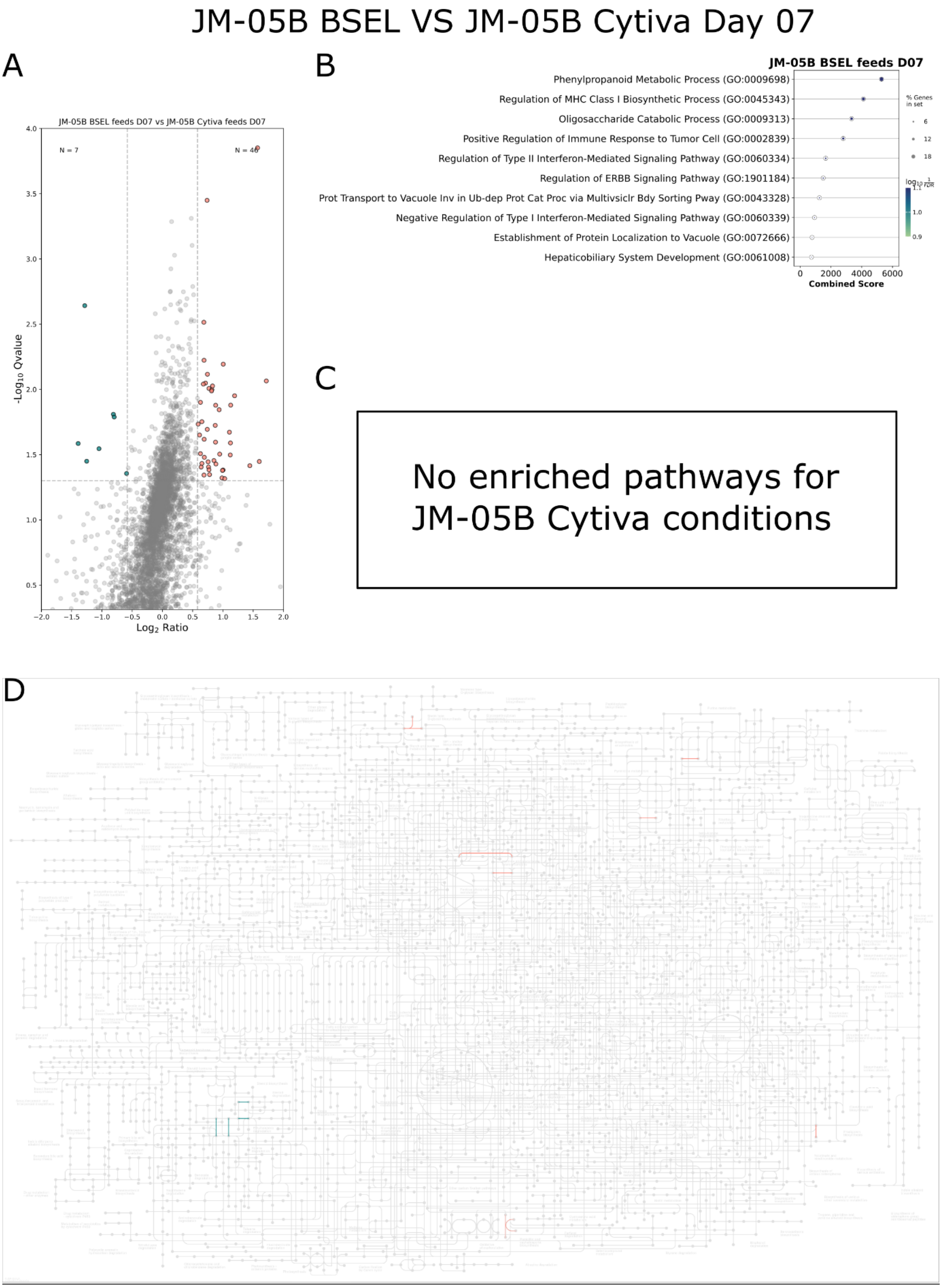


**Figure S2:** JM-05B BSEL (blue) vs JM-05B Cytiva (pink) D07 A) Volcano plot, B-C) gene set enrichment analysis D) KEGG mapped pathways

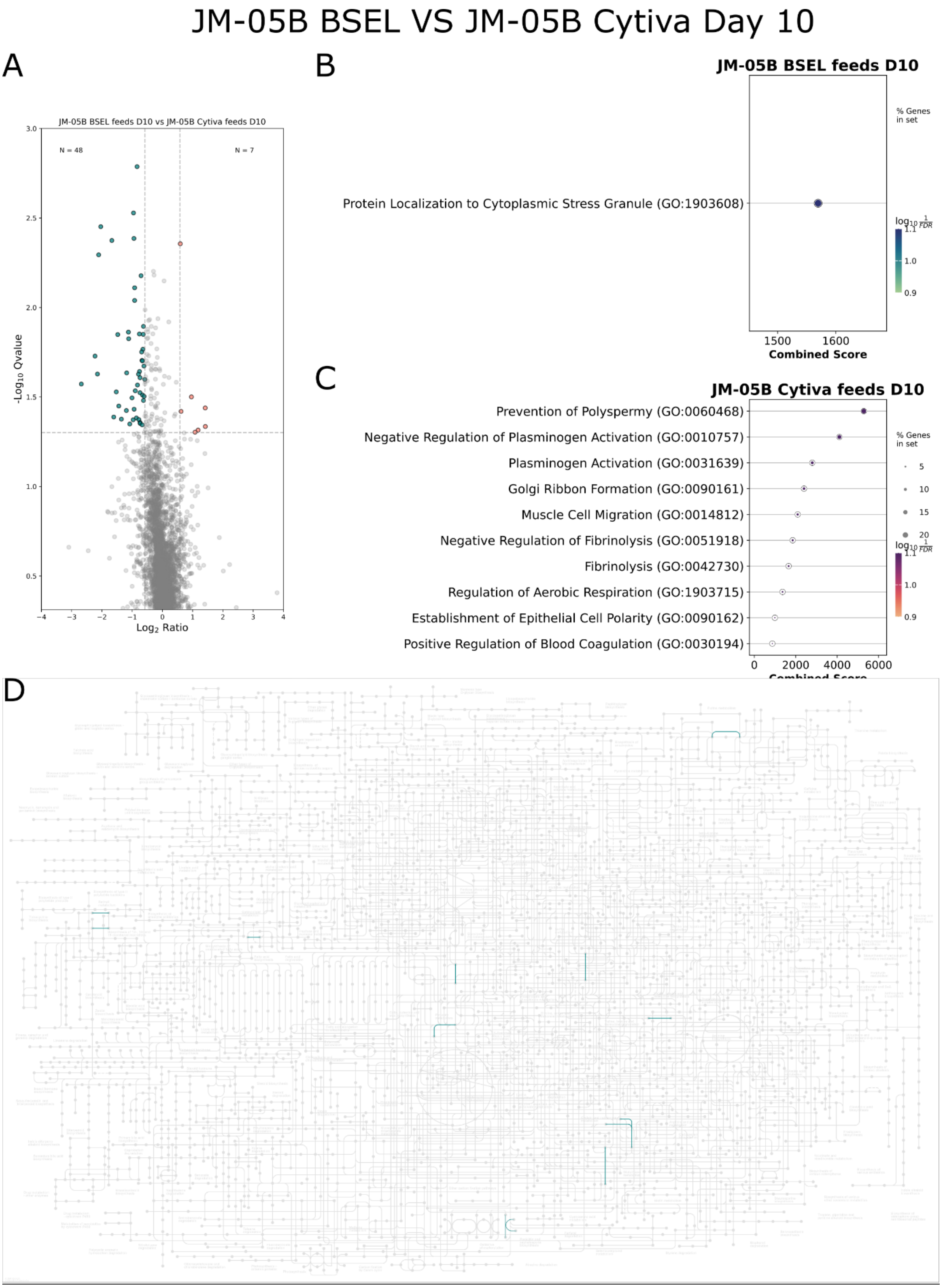


**Figure S3:** JM-05B BSEL (blue) vs JM-05B Cytiva (pink) D05 A) Volcano plot, B-C) gene set enrichment analysis D) KEGG mapped pathwa10

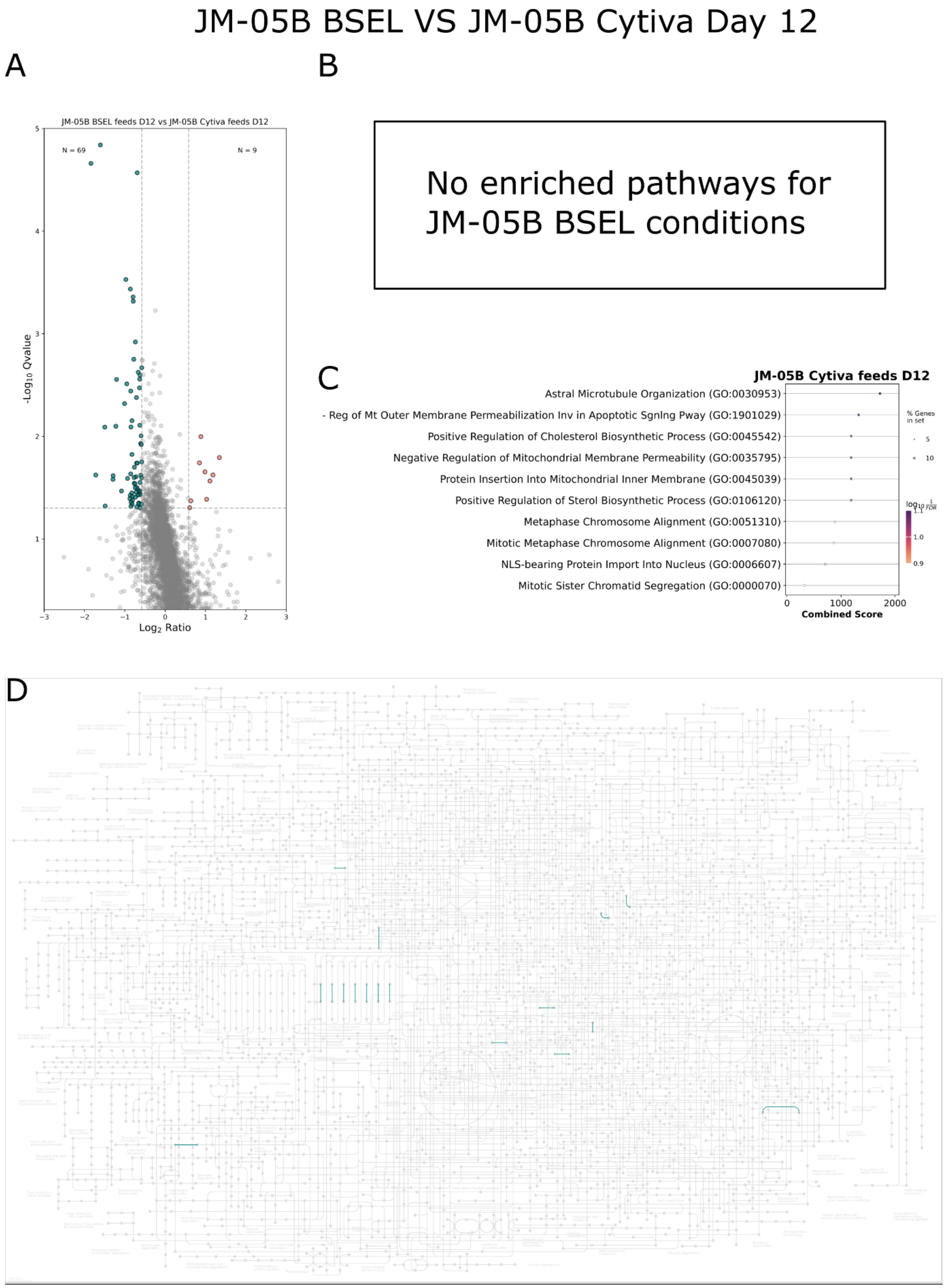


**Figure S4:** JM-05B BSEL (blue) vs JM-05B Cytiva (pink) D12 A) Volcano plot, B-C) gene set enrichment analysis D) KEGG mapped pathways

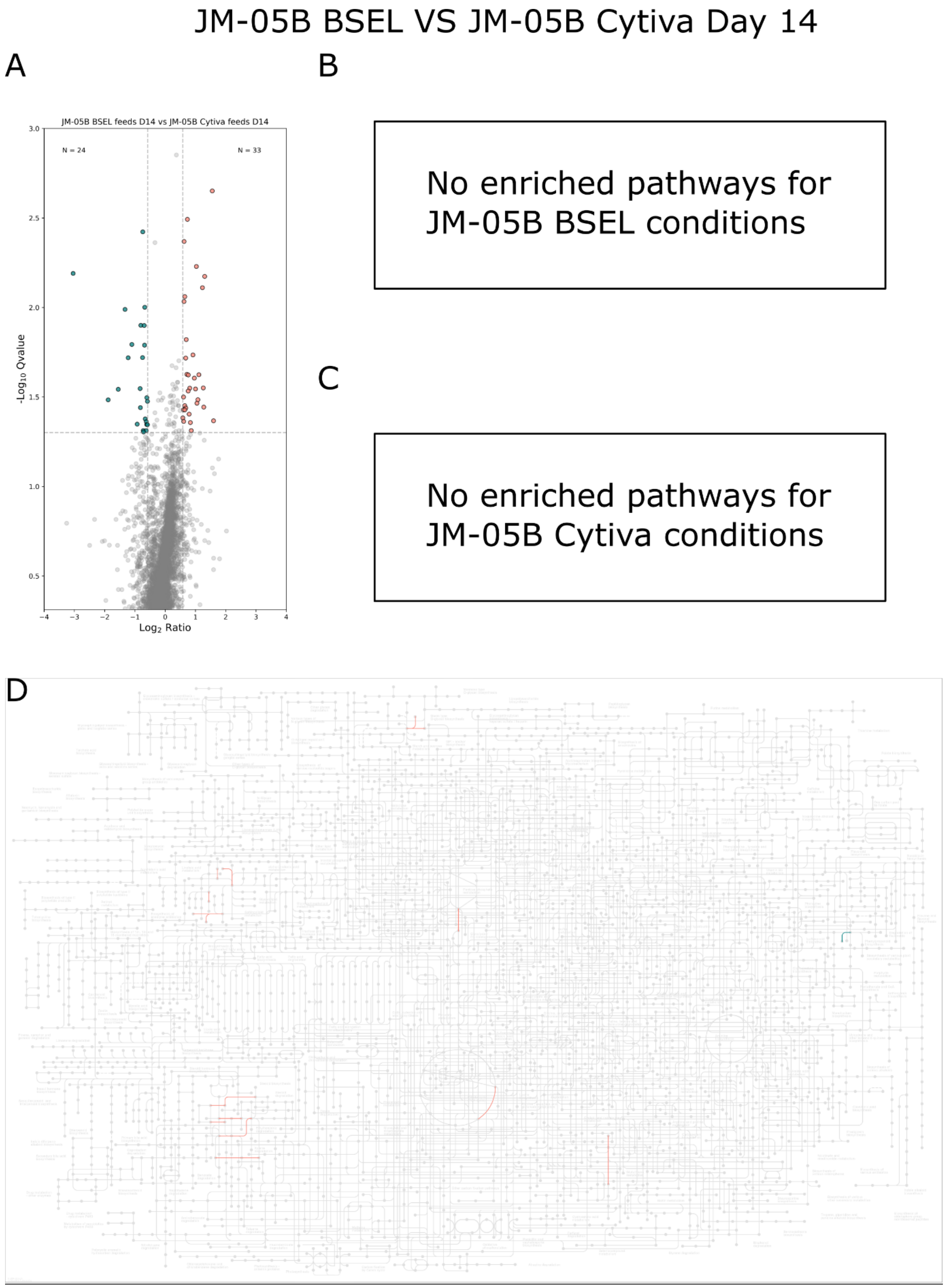


**Figure S5** JM-05B BSEL (blue) vs JM-05B Cytiva (pink) D14 A) Volcano plot, B-C) gene set enrichment analysis D) KEGG mapped pathways

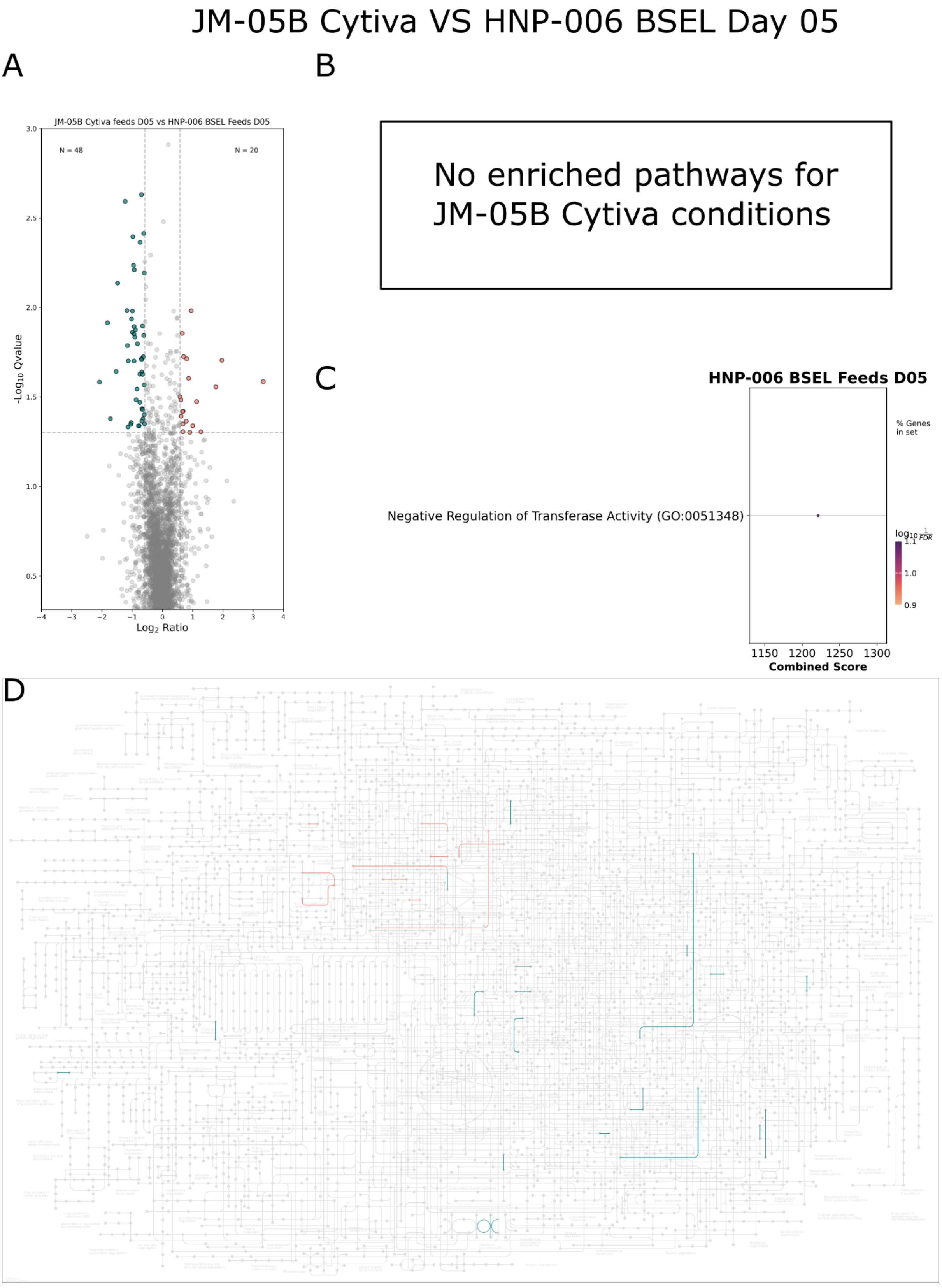


**Figure S6** JM-05B Cytiva (blue) vs HNP-006 BSEL (pink) D05 A) Volcano plot, B-C) gene set enrichment analysis D) KEGG mapped pathways

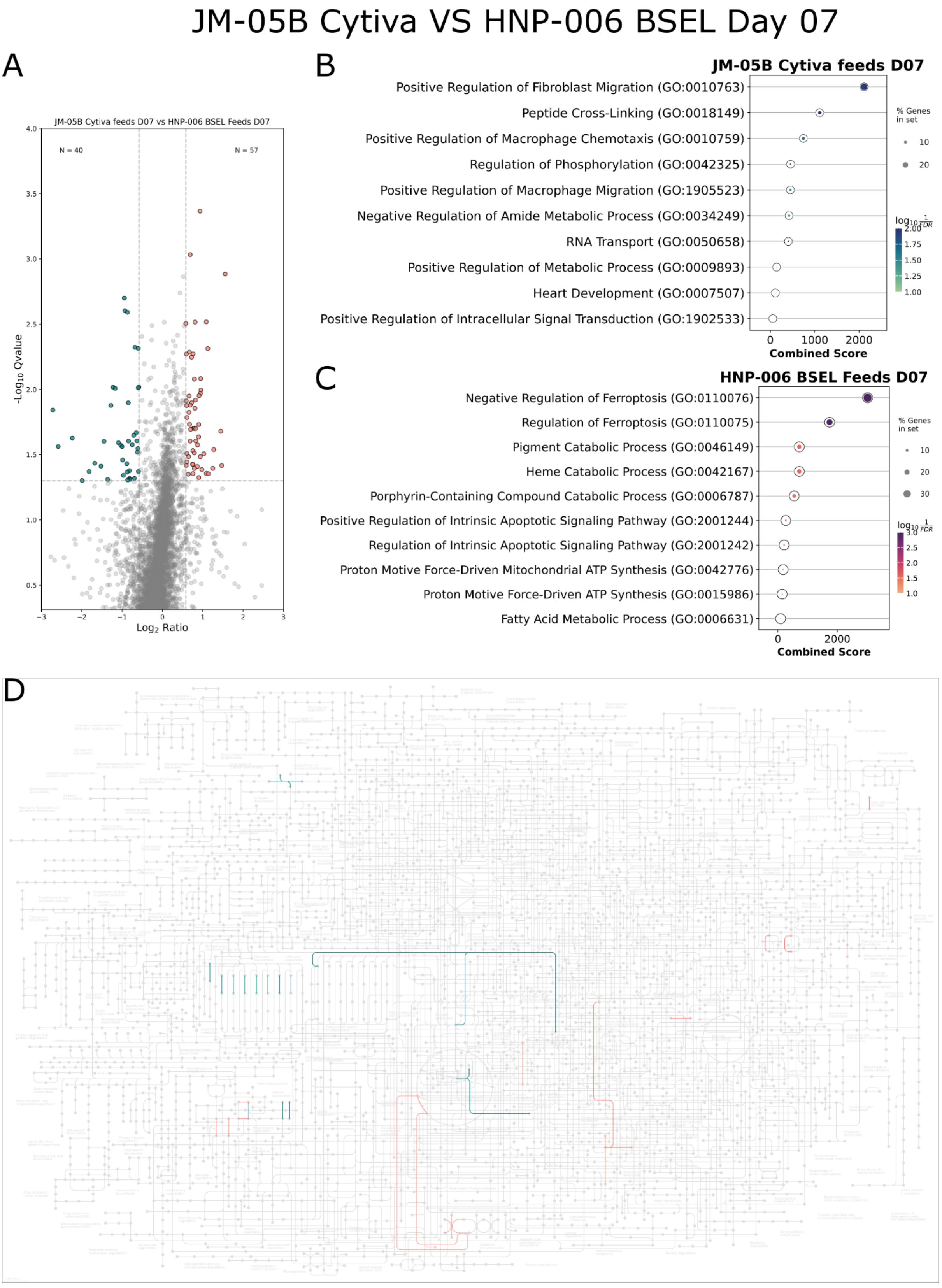


**Figure S7**: JM-05B Cytiva (blue) vs HNP-006 BSEL (pink) D07 A) Volcano plot, B-C) gene set enrichment analysis D) KEGG mapped pathways

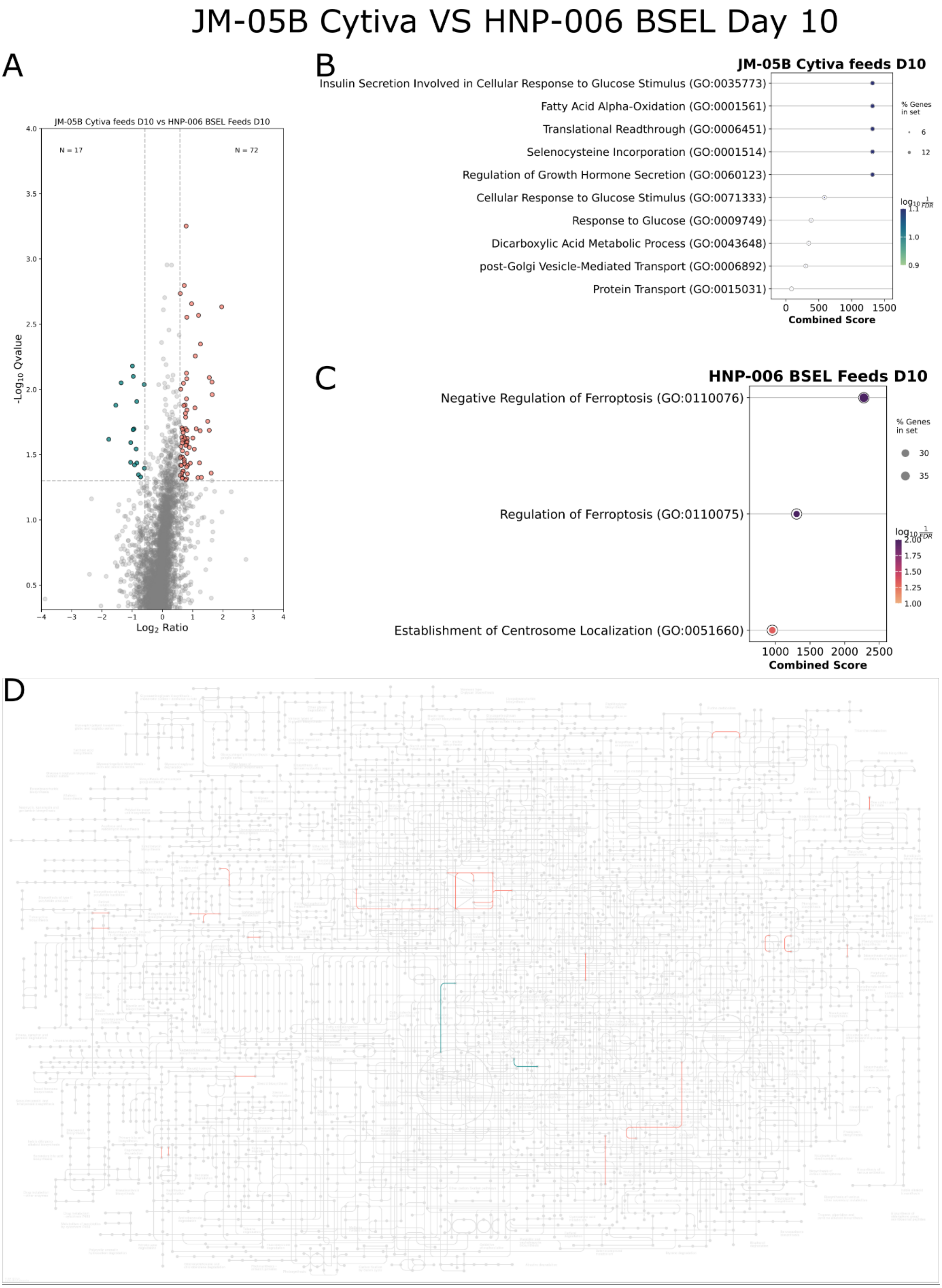


**Figure S8**: JM-05B Cytiva (blue) vs HNP-006 BSEL (pink) D10 A) Volcano plot, B-C) gene set enrichment analysis D) KEGG mapped pathways

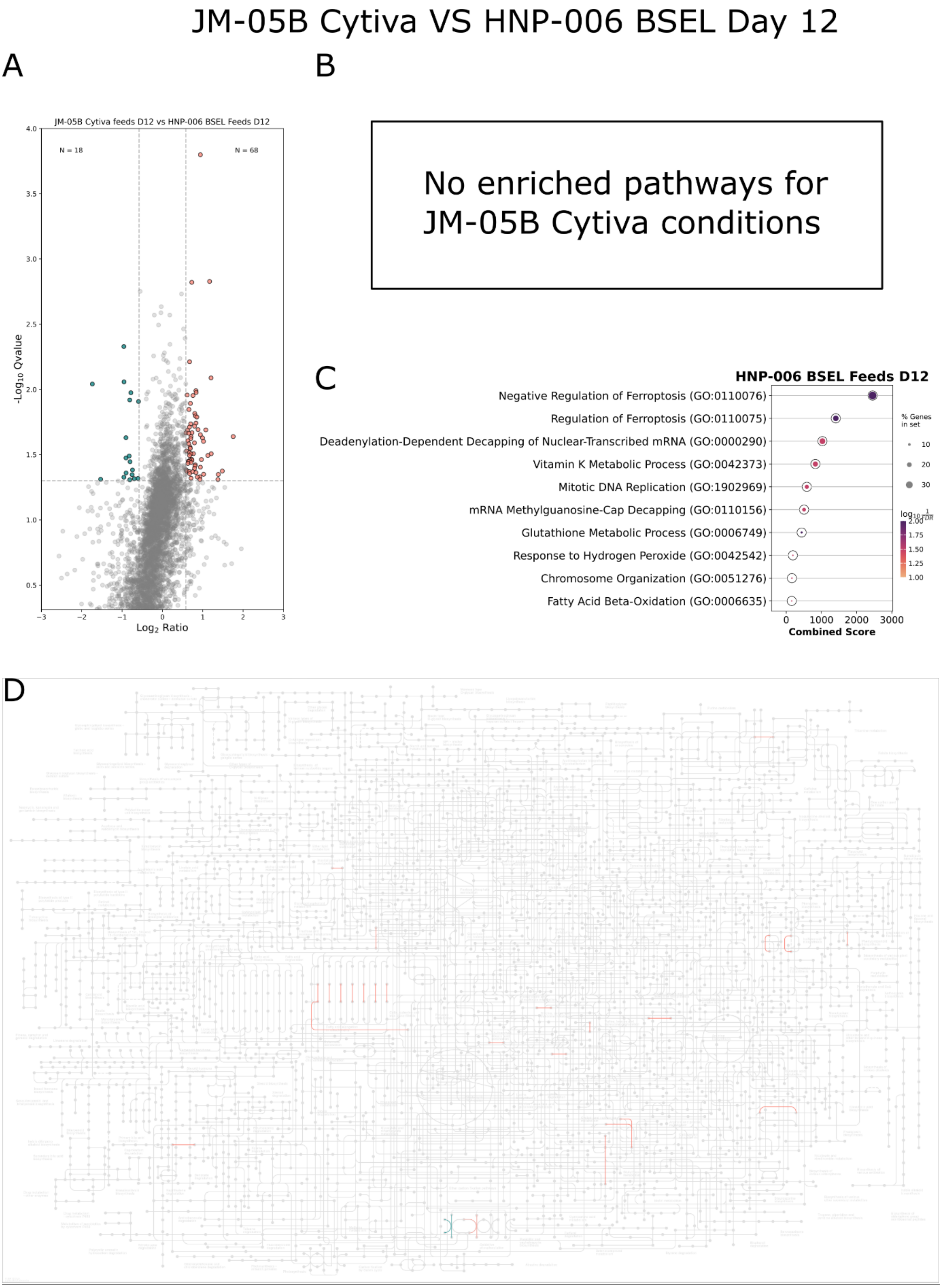


**Figure S9:** JM-05B Cytiva (blue) vs HNP-006 BSEL (pink) D12 A) Volcano plot, B-C) gene set enrichment analysis D) KEGG mapped pathways

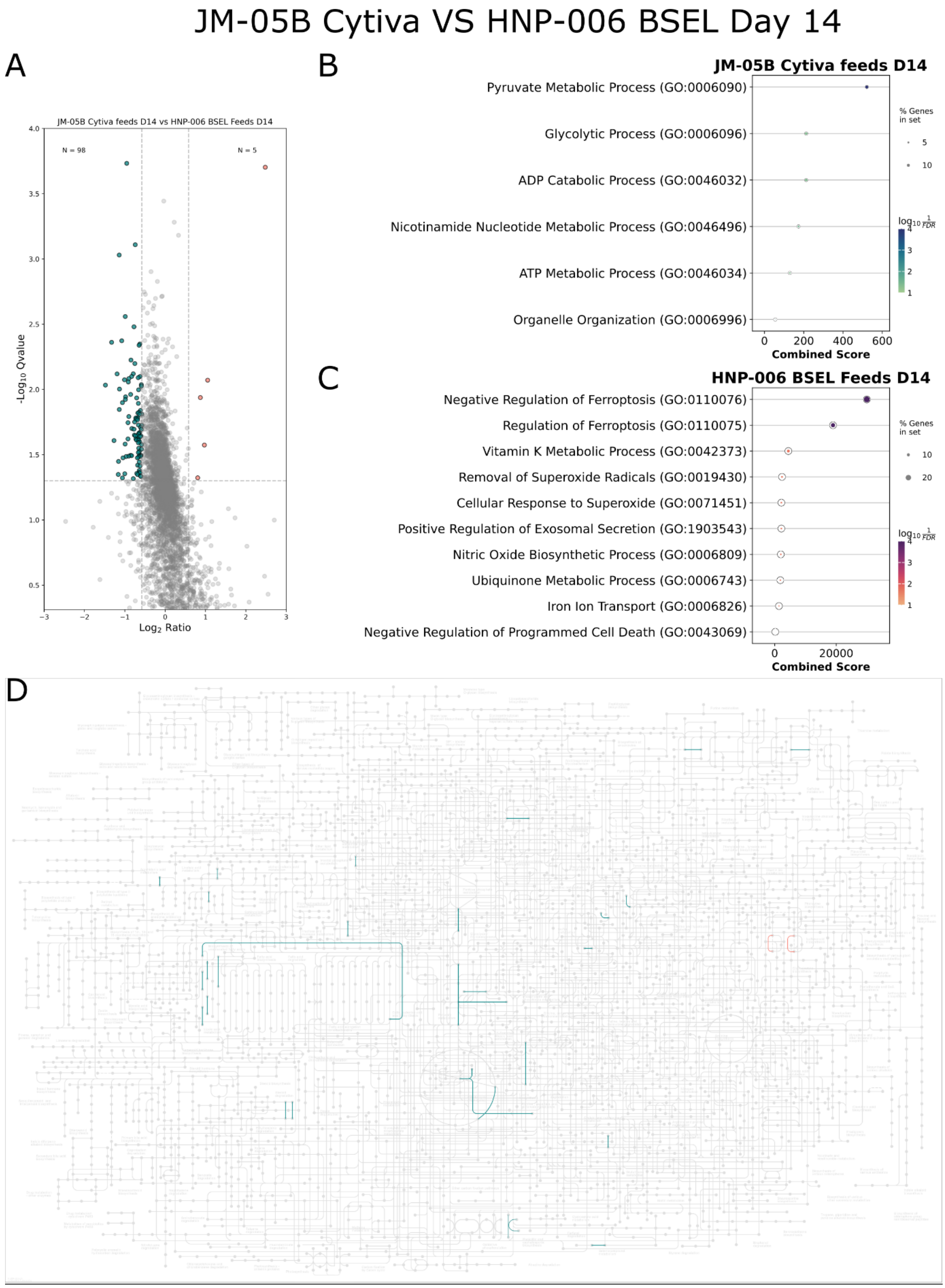


**Figure S10**: JM-05B Cytiva (blue) vs HNP-006 BSEL (pink) D14 A) Volcano plot, B-C) gene set enrichment analysis D) KEGG mapped pathways

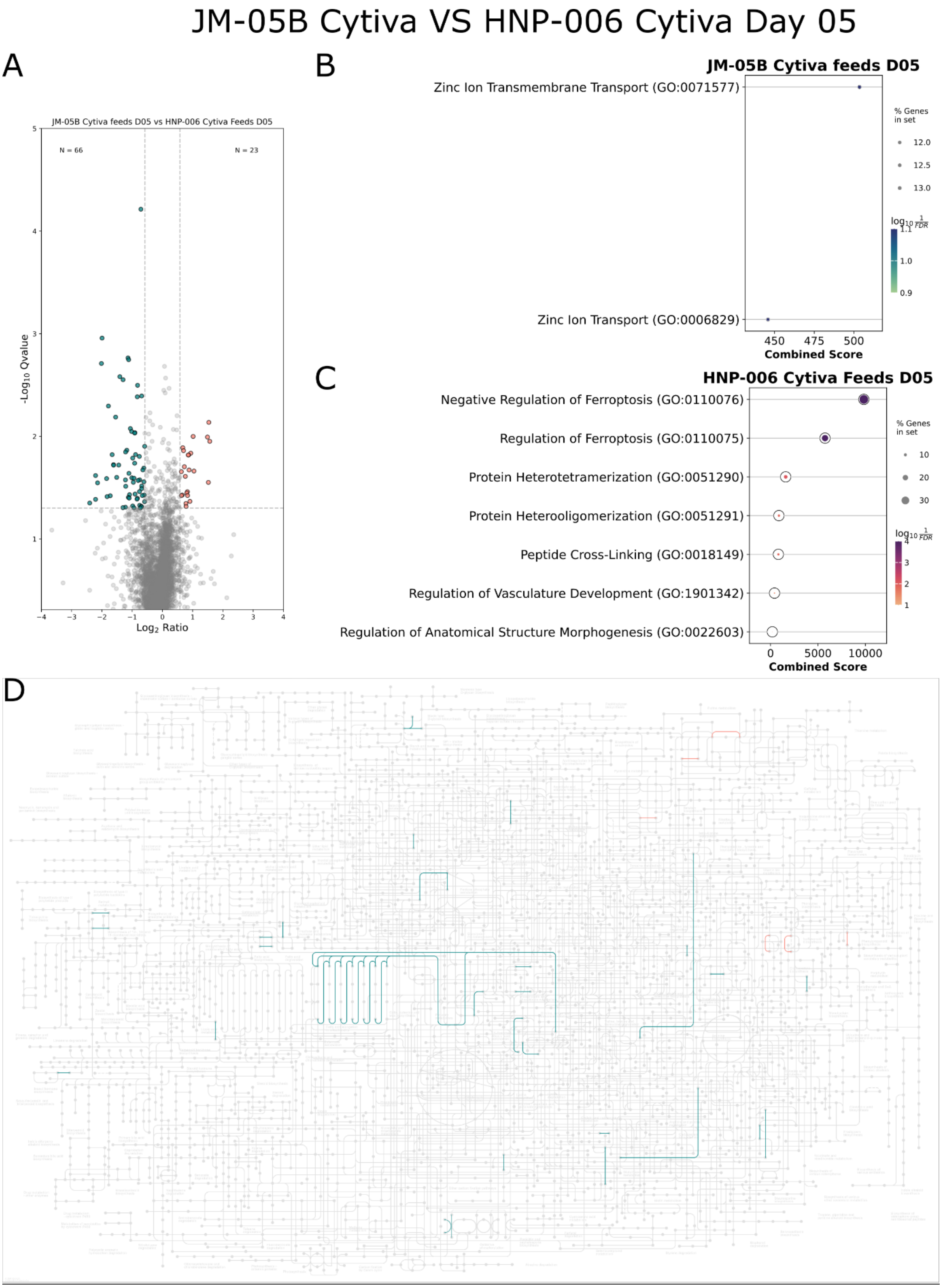


**Figure S11**: JM-05B Cytiva (blue) vs HNP-006 Cytiva (pink) D05 A) Volcano plot, B-C) gene set enrichment analysis D) KEGG mapped pathways

JM-05B Cytiva VS HNP-006 Cytiva Day 07

A

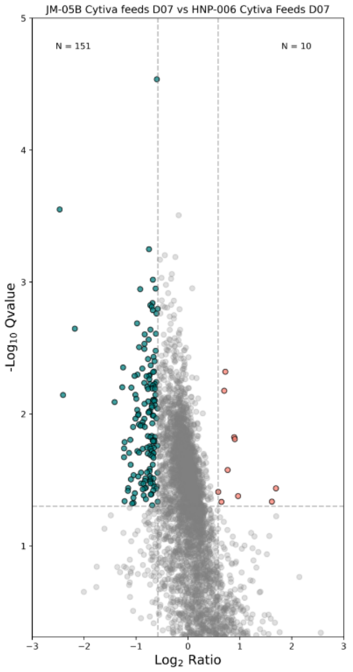


B

No enriched pathways for JM-05B Cytiva conditions

C

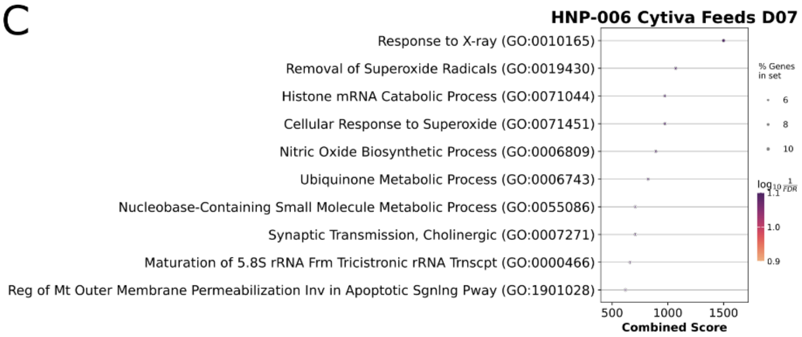


D

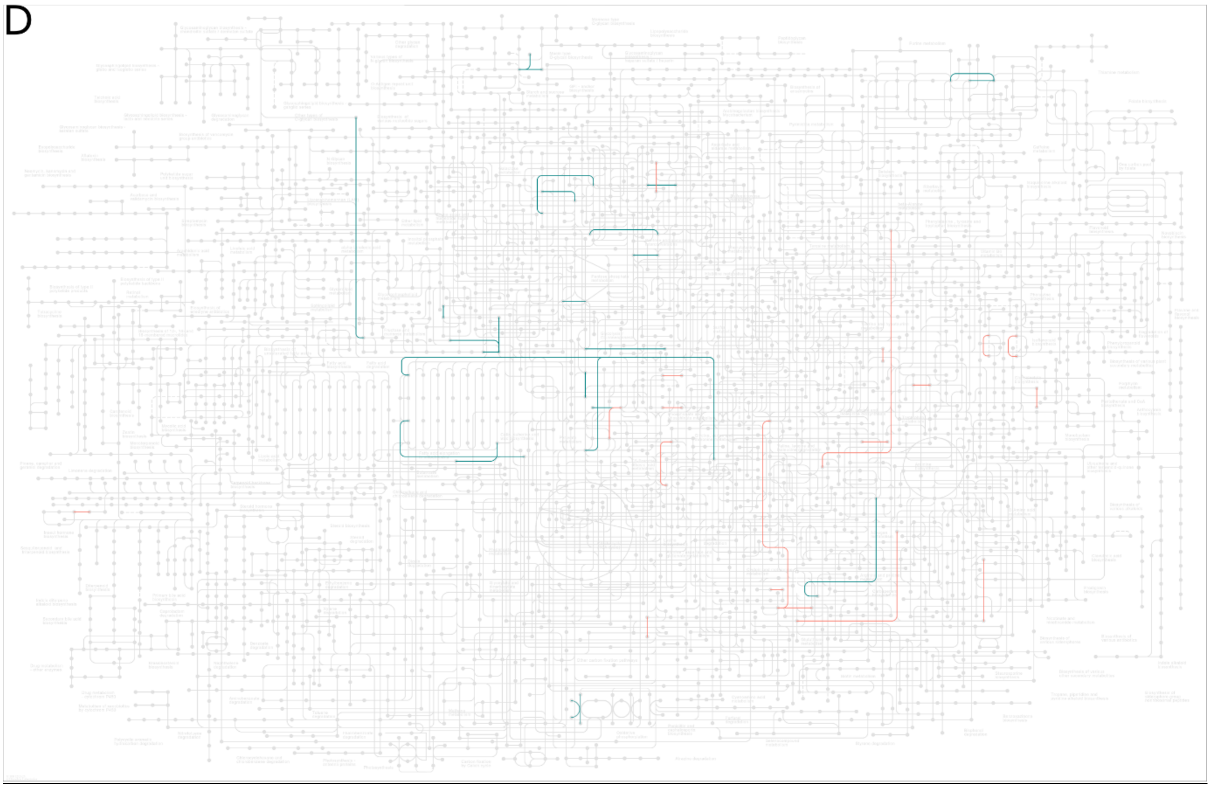

**Figure S12**: JM-05B Cytiva (blue) vs HNP-006 Cytiva (pink) D07 A) Volcano plot, B-C) gene set enrichment analysis D) KEGG mapped pathways

### JM-05B Cytiva VS HNP-006 Cytiva Day 10

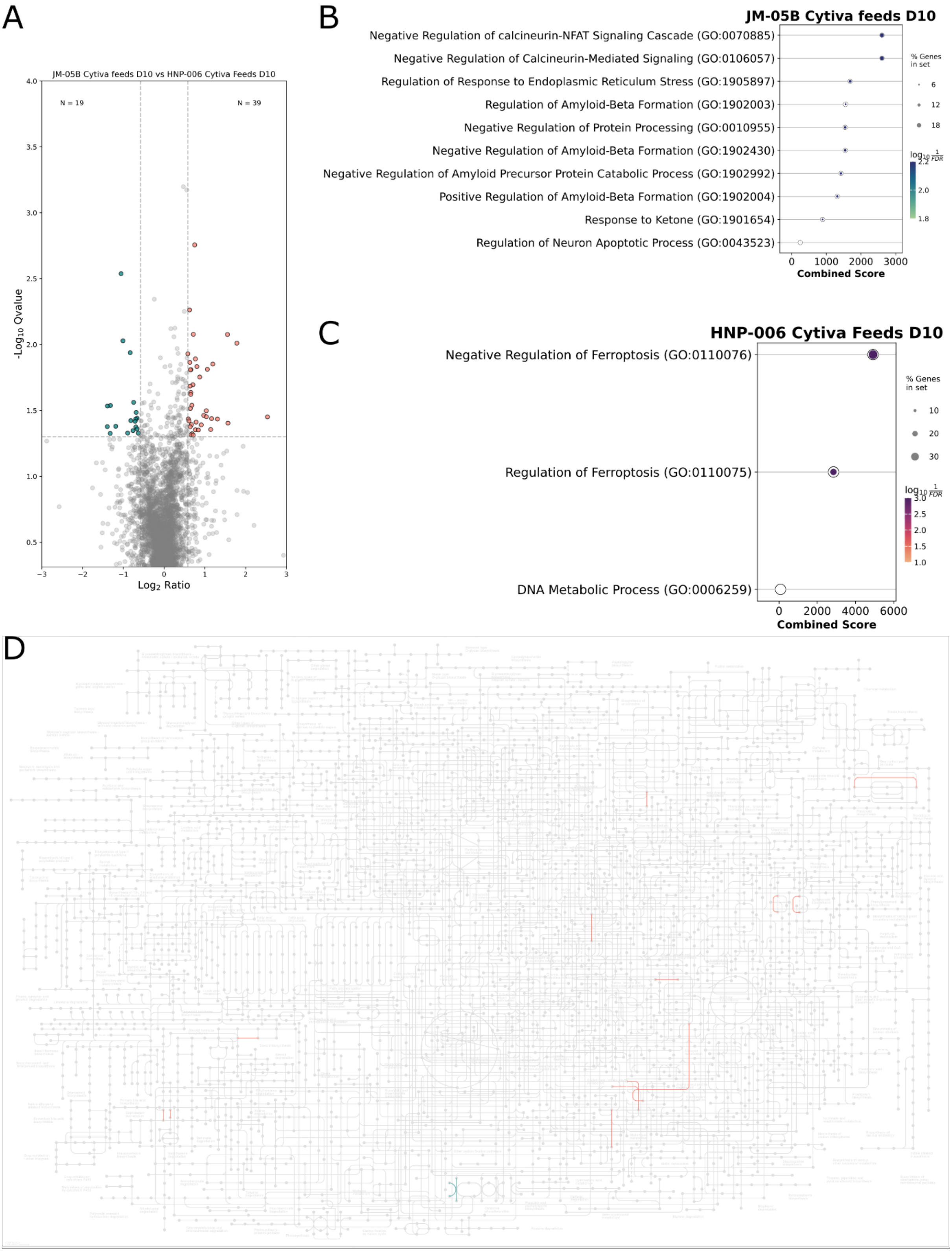


**Figure S13**: JM-05B Cytiva (blue) vs HNP-006 Cytiva (pink) D10 A) Volcano plot, B-C) gene set enrichment analysis D) KEGG mapped pathways

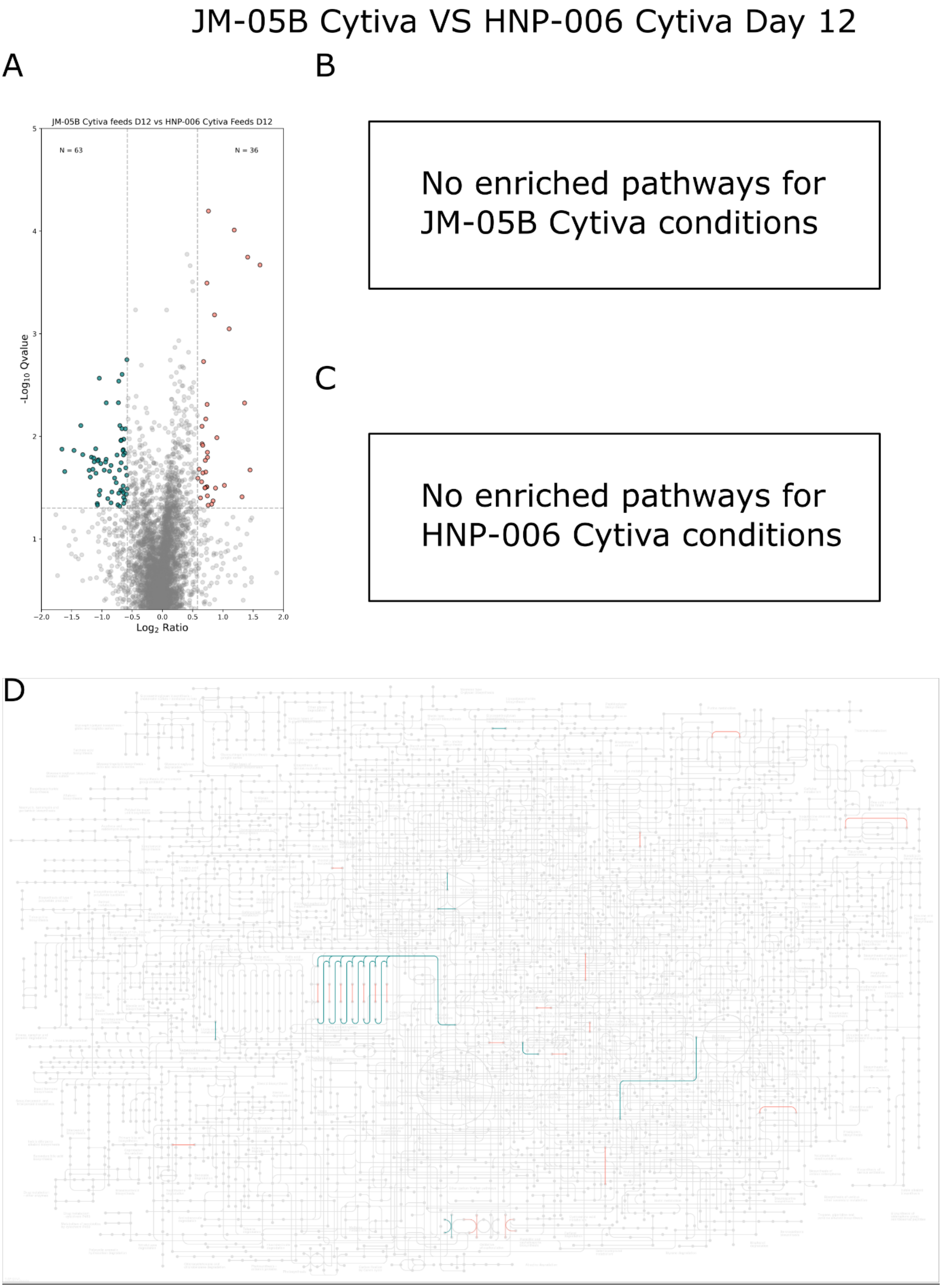


**Figure S14**: JM-05B Cytiva (blue) vs HNP-006 Cytiva (pink) D12 A) Volcano plot, B-C) gene set enrichment analysis D) KEGG mapped pathways

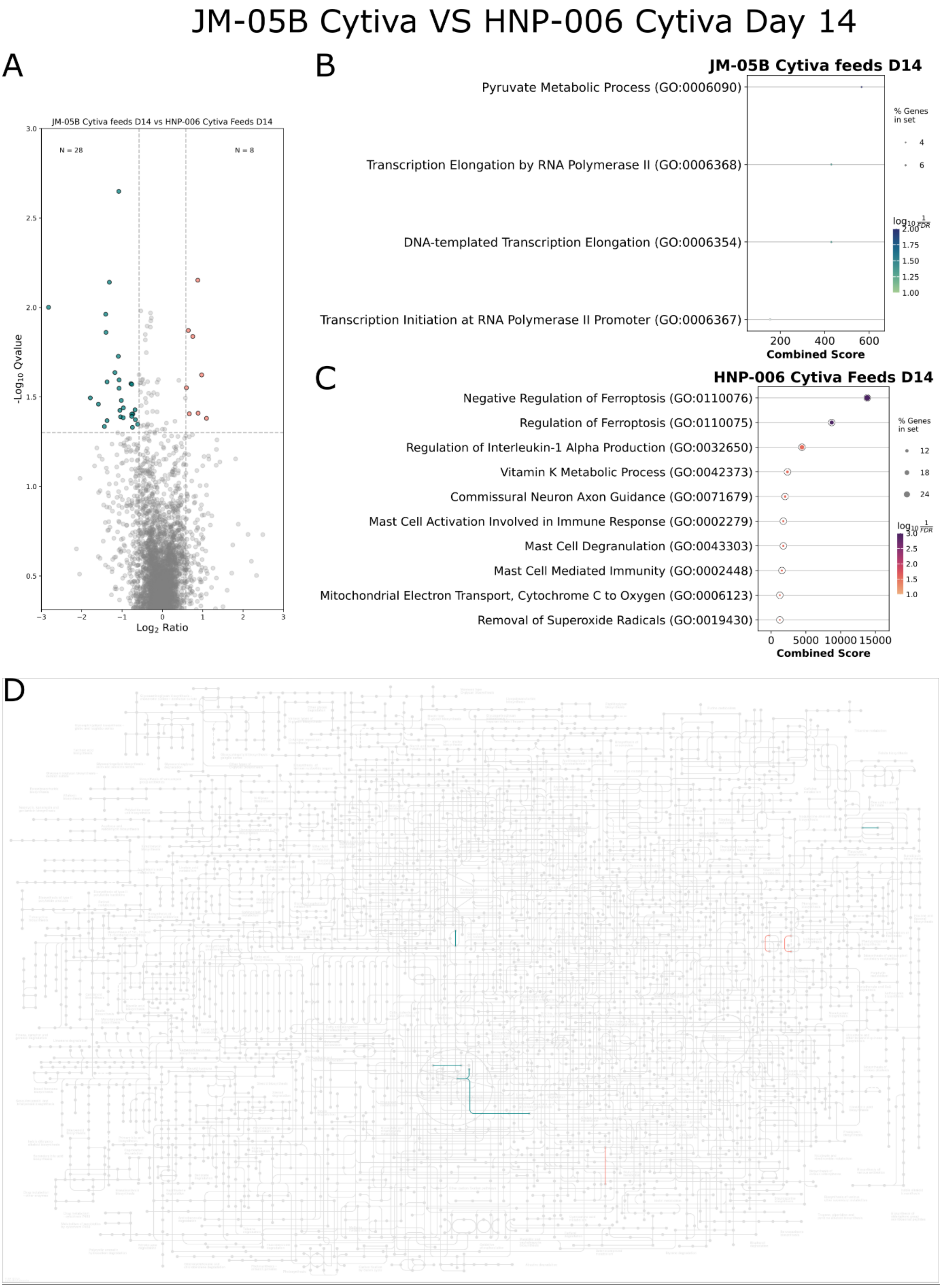


**Figure S15**: JM-05B Cytiva (blue) vs HNP-006 Cytiva (pink) D14 A) Volcano plot, B-C) gene set enrichment analysis D) KEGG mapped pathways

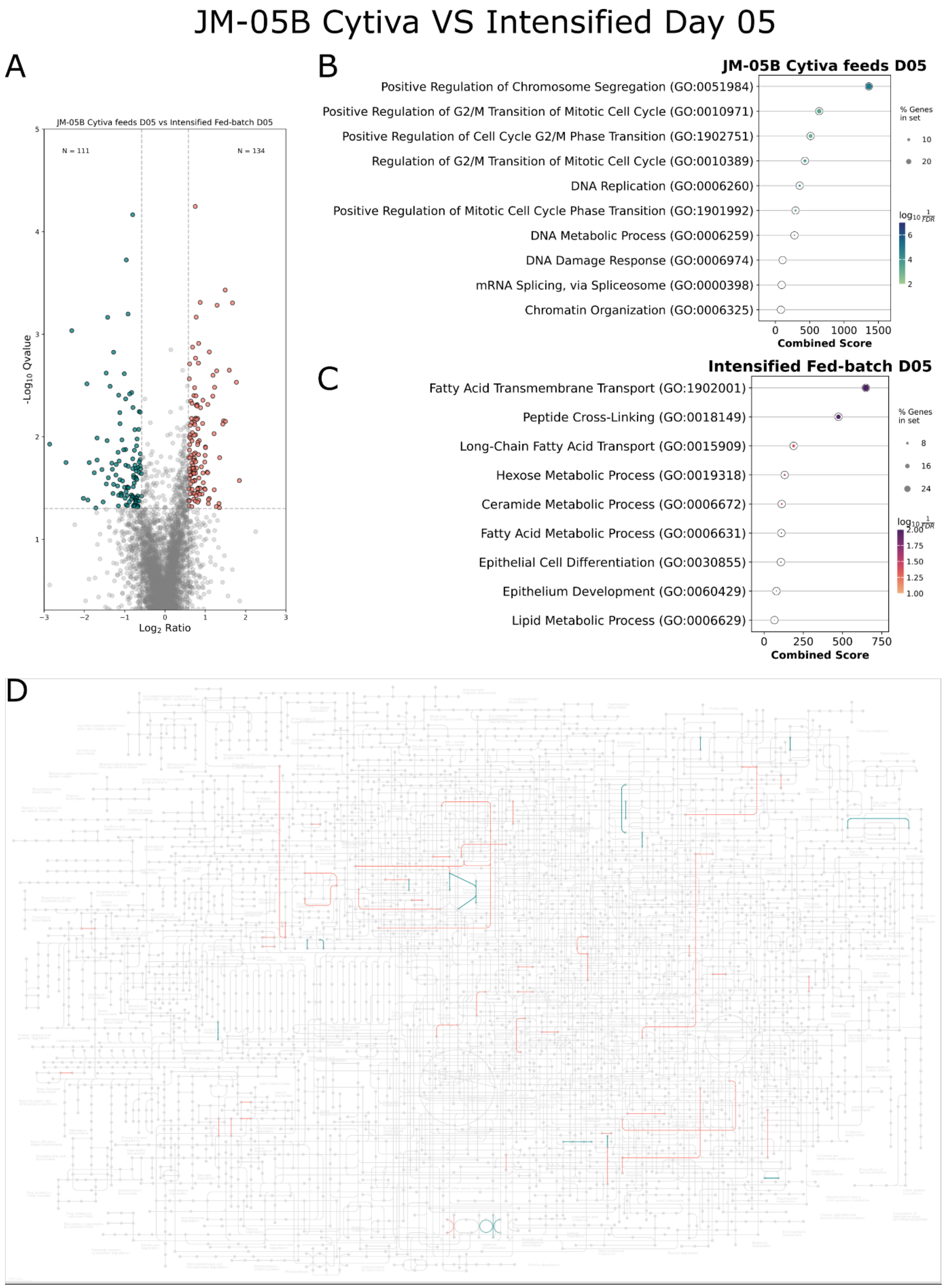


**Figure S16**: JM-05B Cytiva (blue) vs Intensified fed batch (pink) D05 A) Volcano plot, B-C) gene set enrichment analysis D) KGG mapped pathways

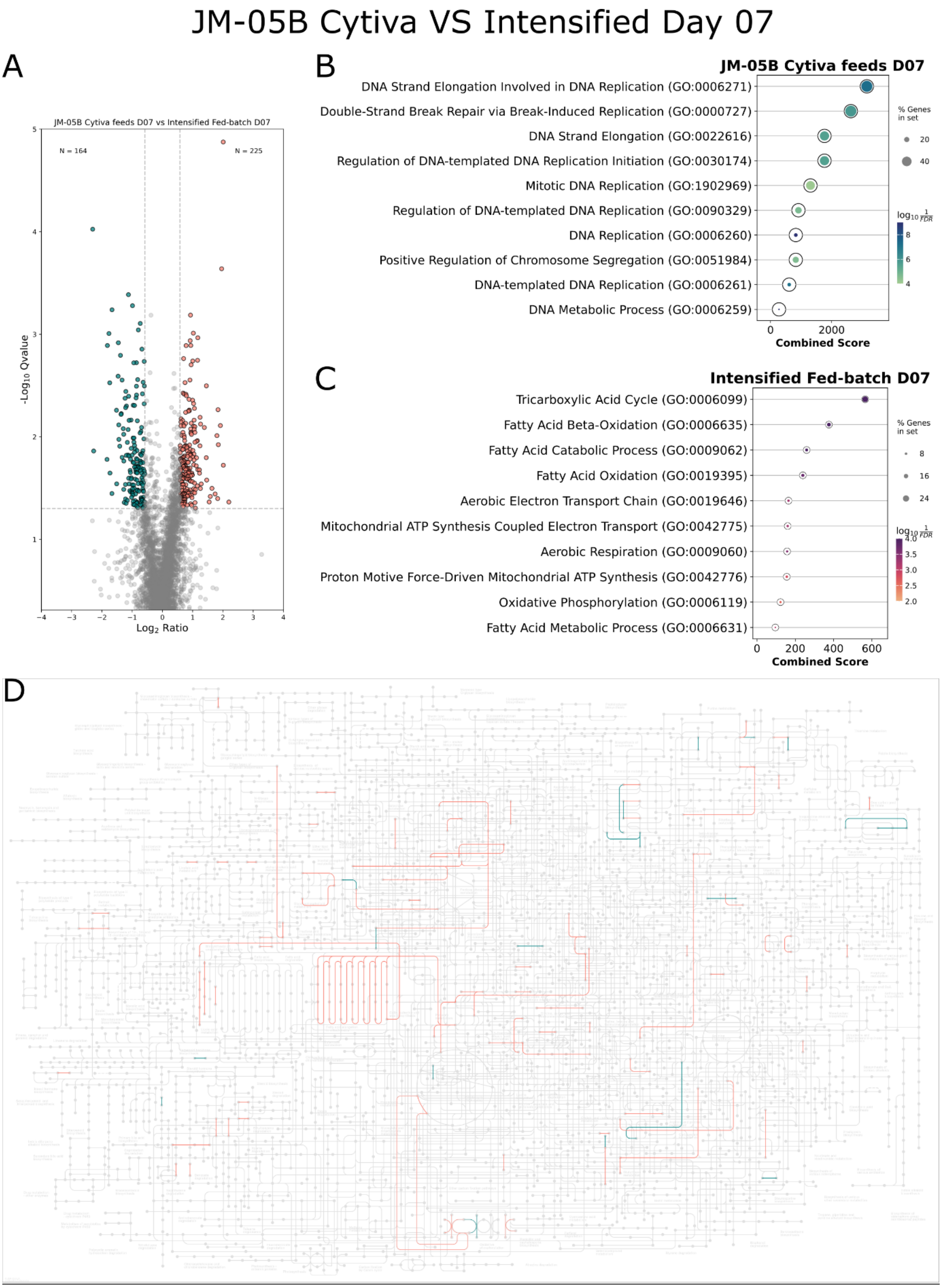


**Figure S17**: JM-05B Cytiva (blue) vs Intensified fed batch (pink) D07 A) Volcano plot, B-C) gene set enrichment analysis D) KEGG mapped pathways

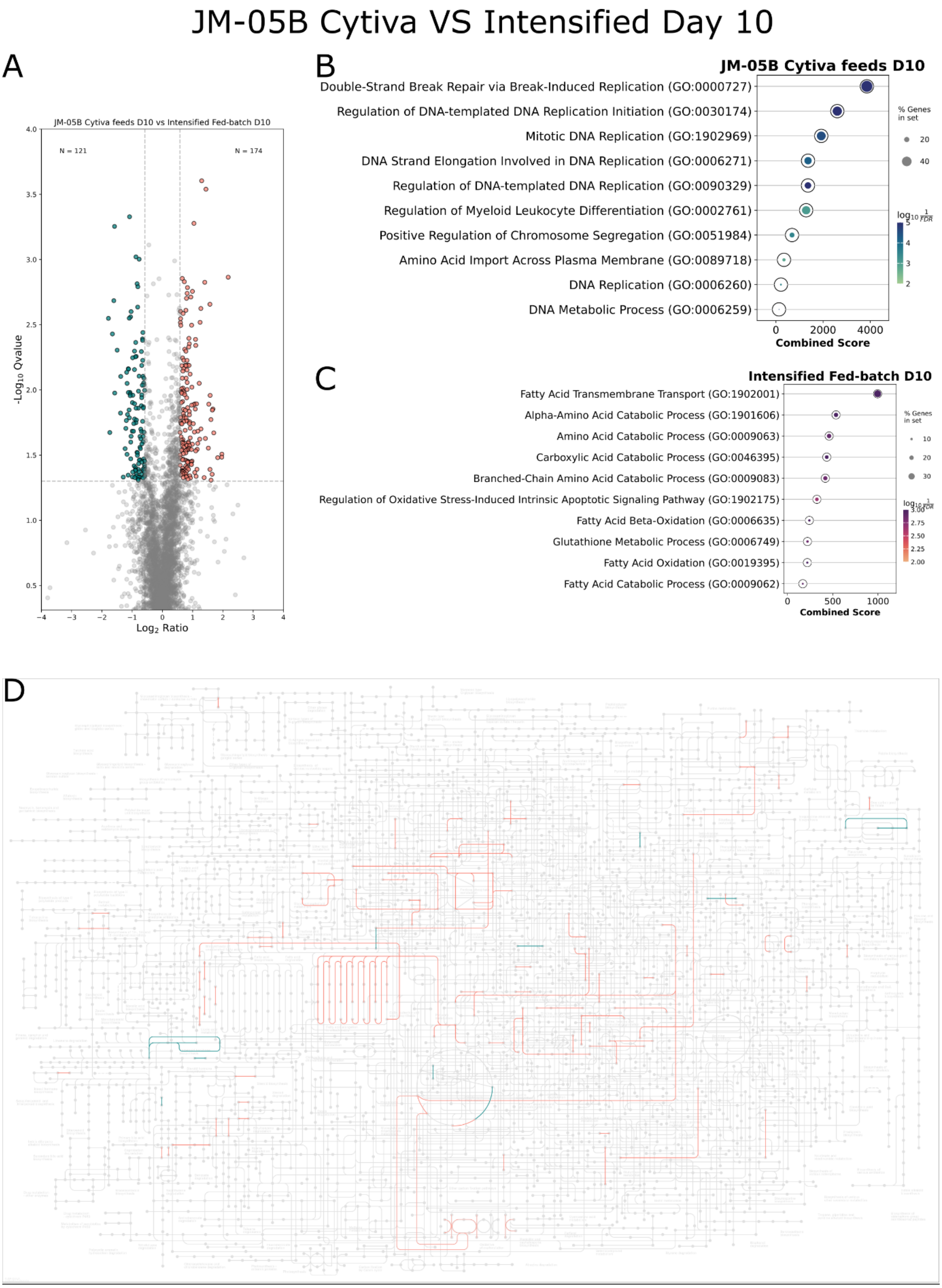


**Figure S18**: JM-05B Cytiva (blue) vs Intensified fed batch (pink) D10 A) Volcano plot, B-C) gene set enrichment analysis D) KEGG mapped pathways

#### JM-05B Cytiva VS Intensified Day 12

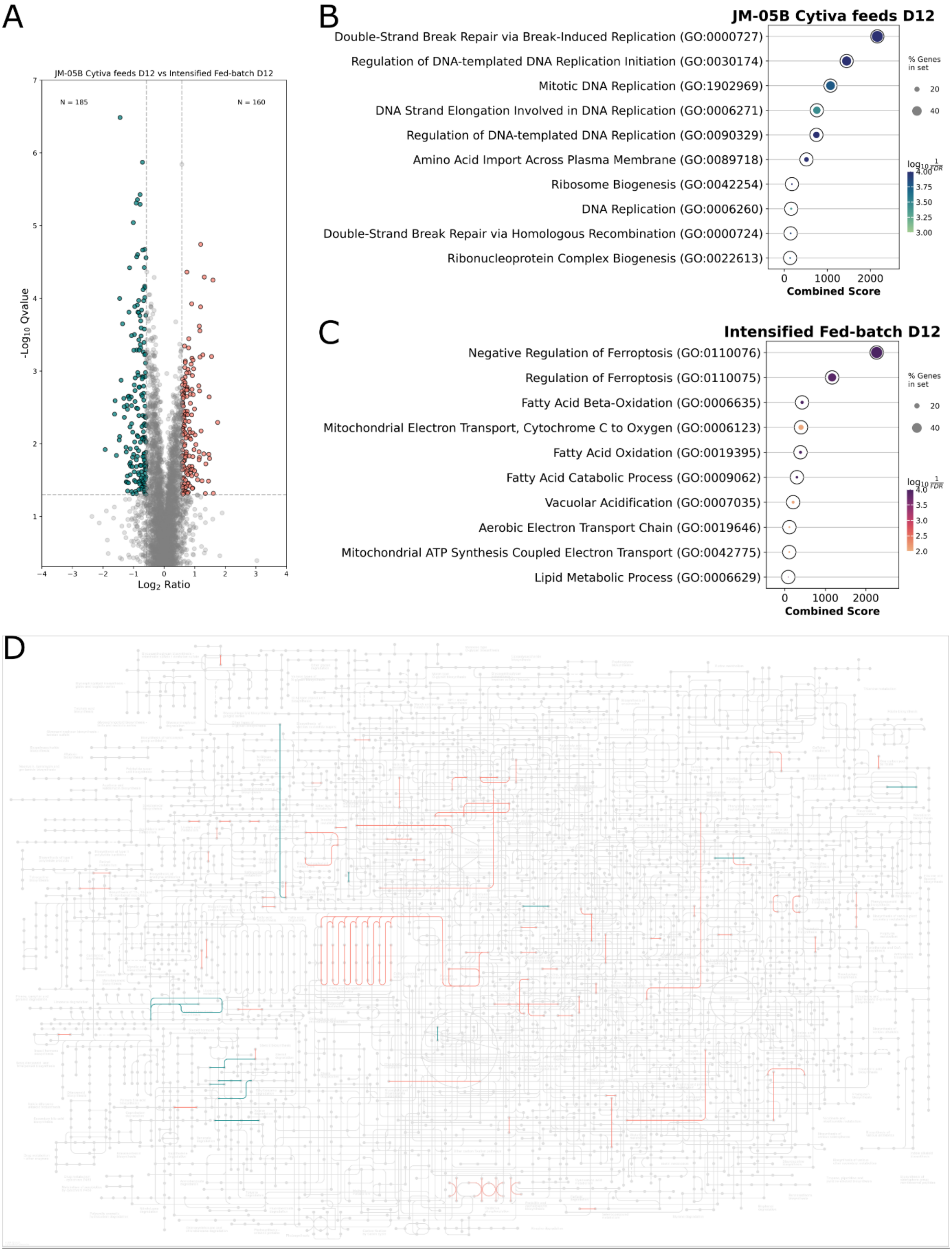


**Figure S19**: JM-05B Cytiva (blue) vs Intensified fed batch (pink) D12 A) Volcano plot, B-C) gene set enrichment analysis D) KEGG mapped pathways

### JM-05B Cytiva VS Intensified Day 14

A

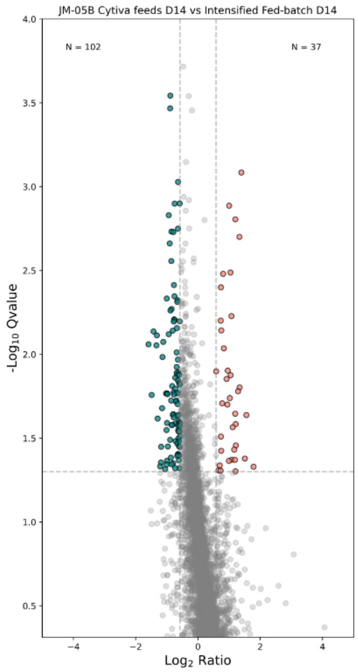


B

No enriched pathways for JM-05B Cytiva conditions

C

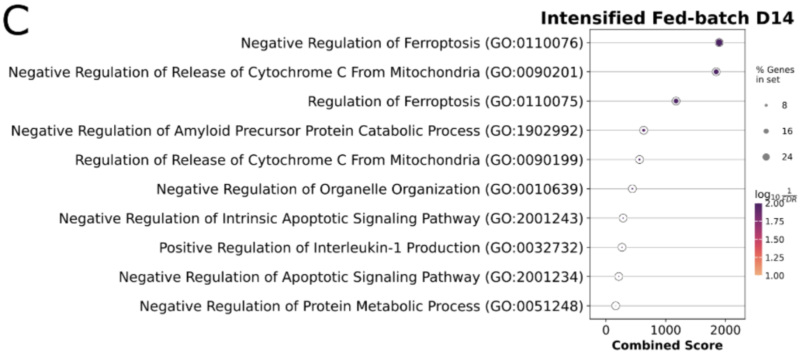


D

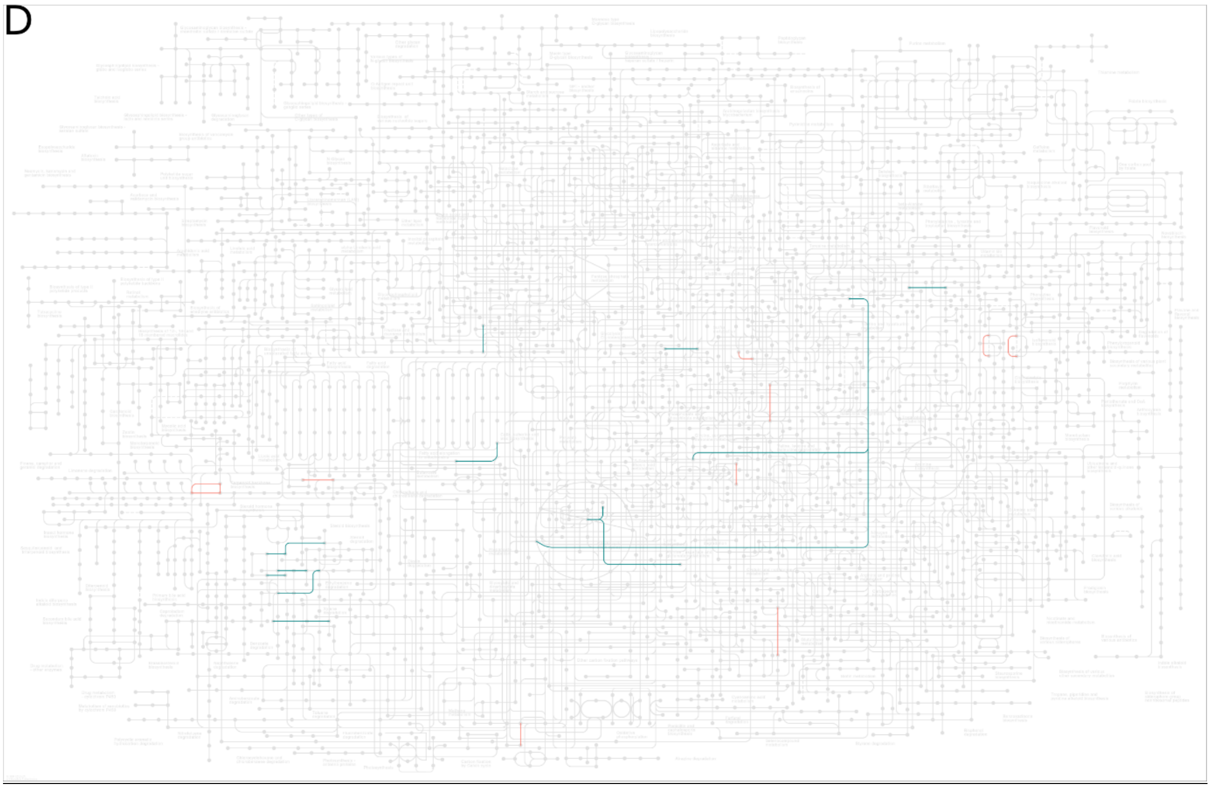

**Figure S20**: JM-05B Cytiva (blue) vs Intensified fed batch (pink) D14 A) Volcano plot, B-C) gene set enrichment analysis D) KEGG mapped pathways

### HNP-006 Cytiva VS HNP-006 BSEL D05

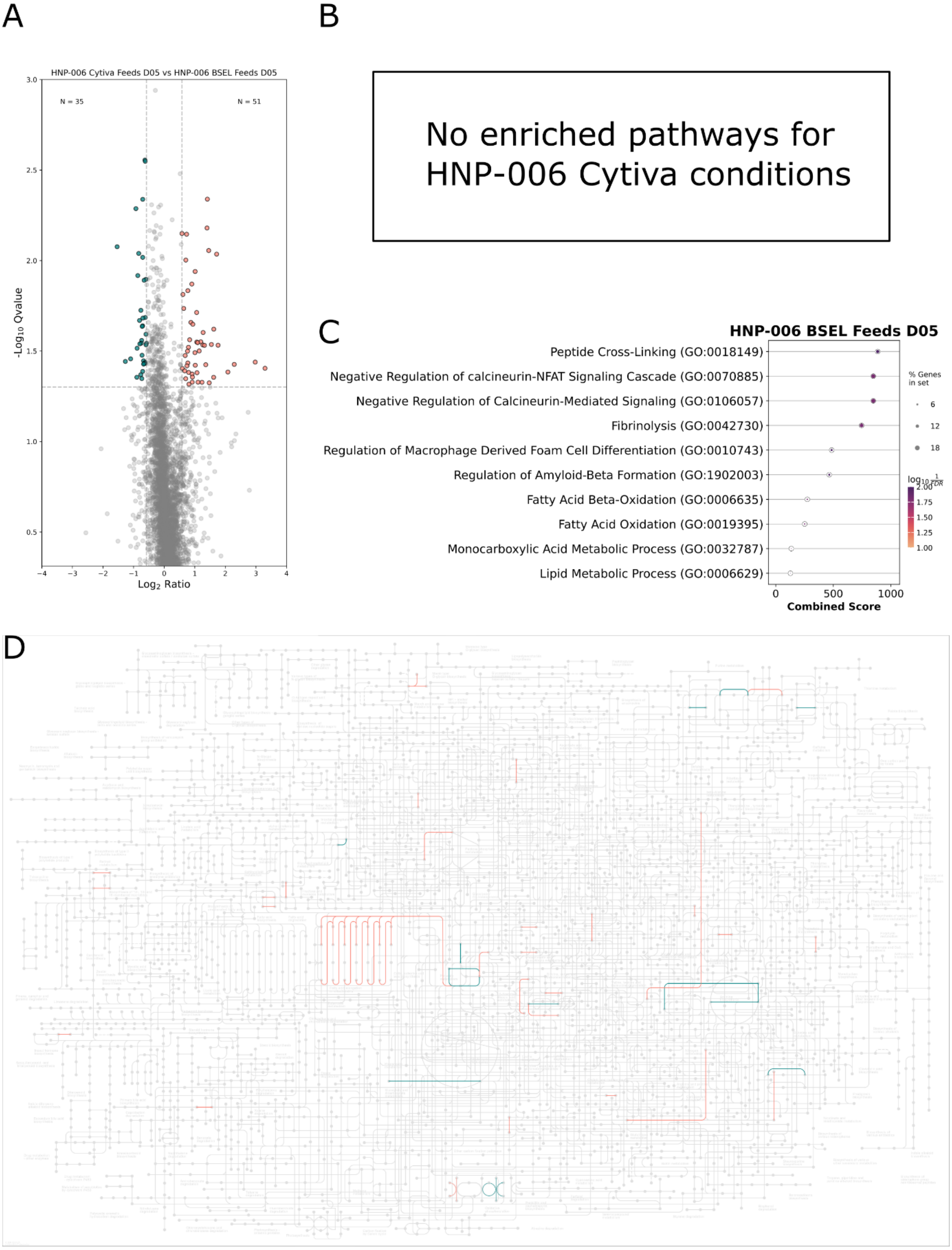


**Figure S21**: HNP-006 Cytiva (blue) vs HNP-006 BSEL (pink) D05 A) Volcano plot, B-C) gene set enrichment analysis D) KEGG mapped pathways

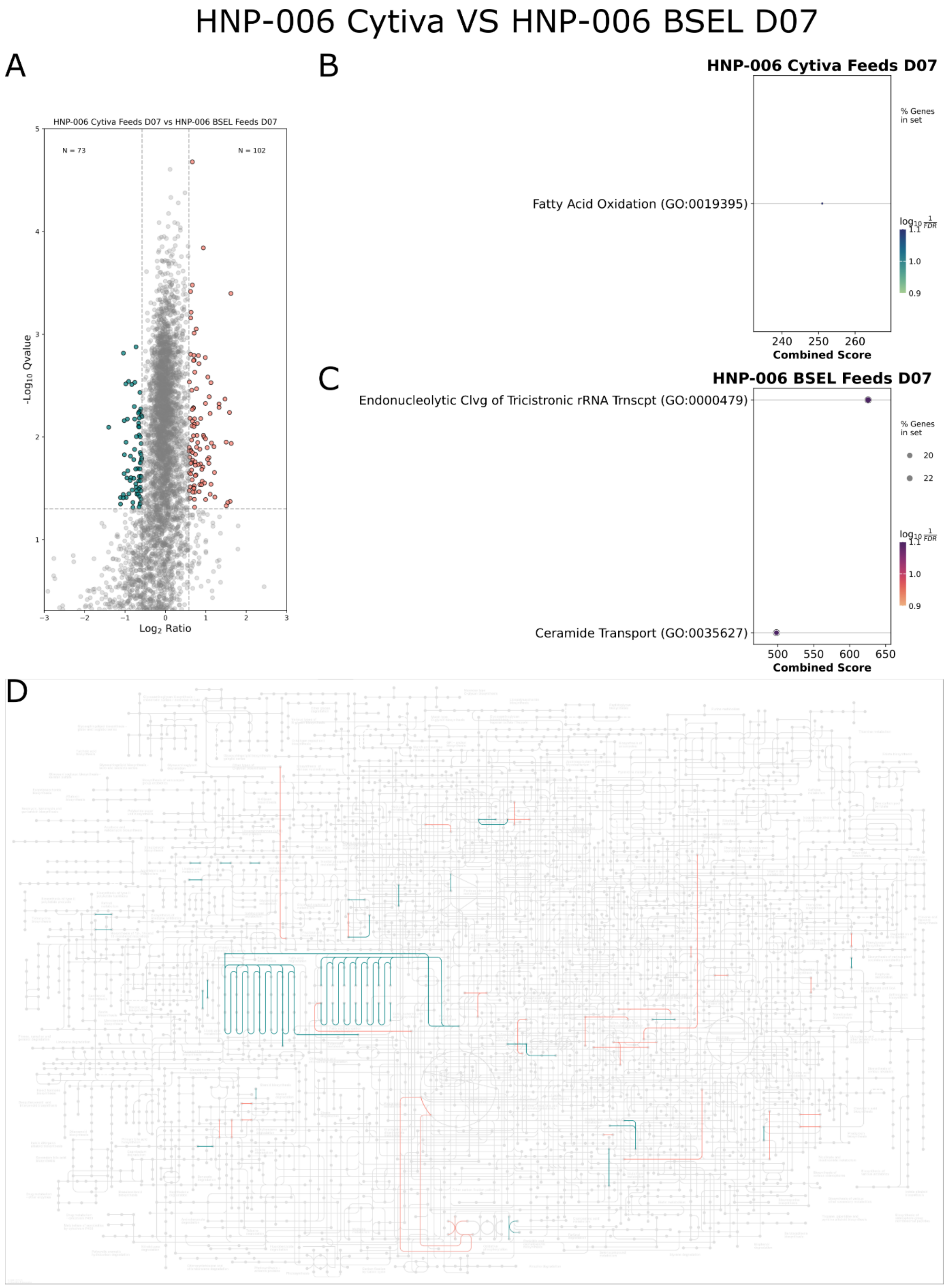


**Figure S22**: HNP-006 Cytiva (blue) vs HNP-006 BSEL (pink) D07 A) Volcano plot, B-C) gene set enrichment analysis D) KEGG mapped pathways

### HNP-006 Cytiva VS HNP-006 BSEL D10

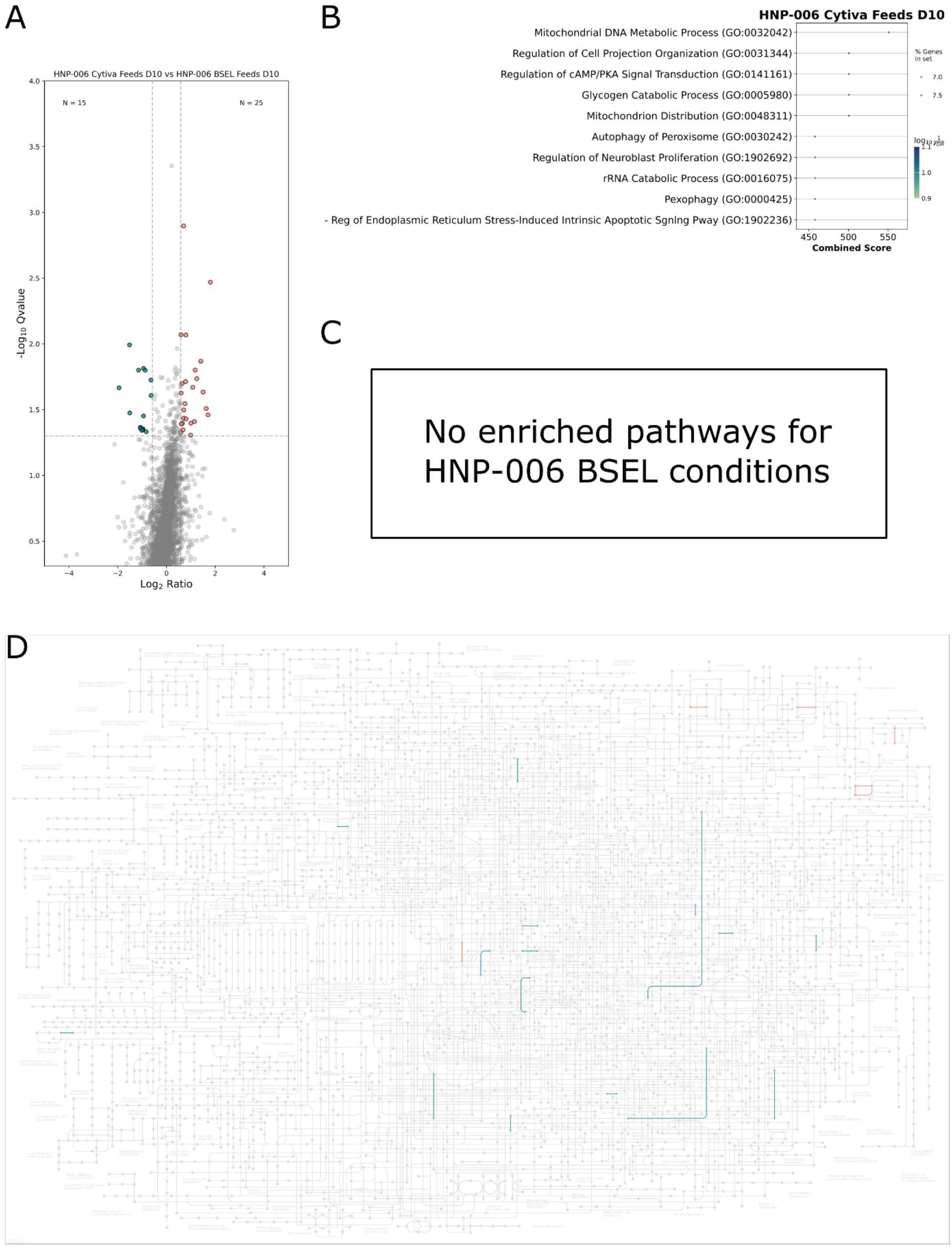


**Figure S23**: HNP-006 Cytiva (blue) vs HNP-006 BSEL (pink) D10 A) Volcano plot, B-C) gene set enrichment analysis D) KEGG mapped pathways

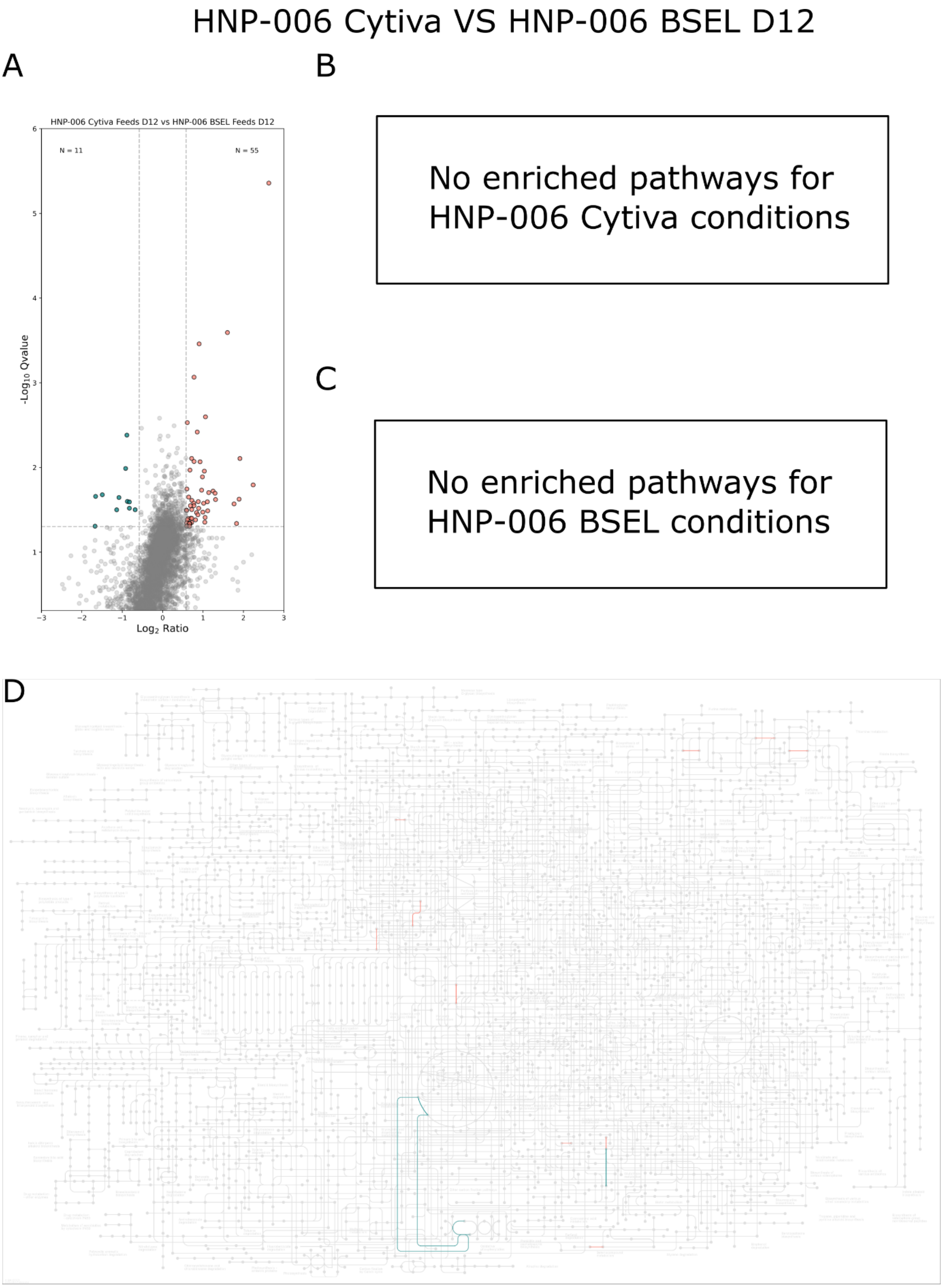


**Figure S24**: HNP-006 Cytiva (blue) vs HNP-006 BSEL (pink) D12 A) Volcano plot, B-C) gene set enrichment analysis D) KEGG mapped pathways

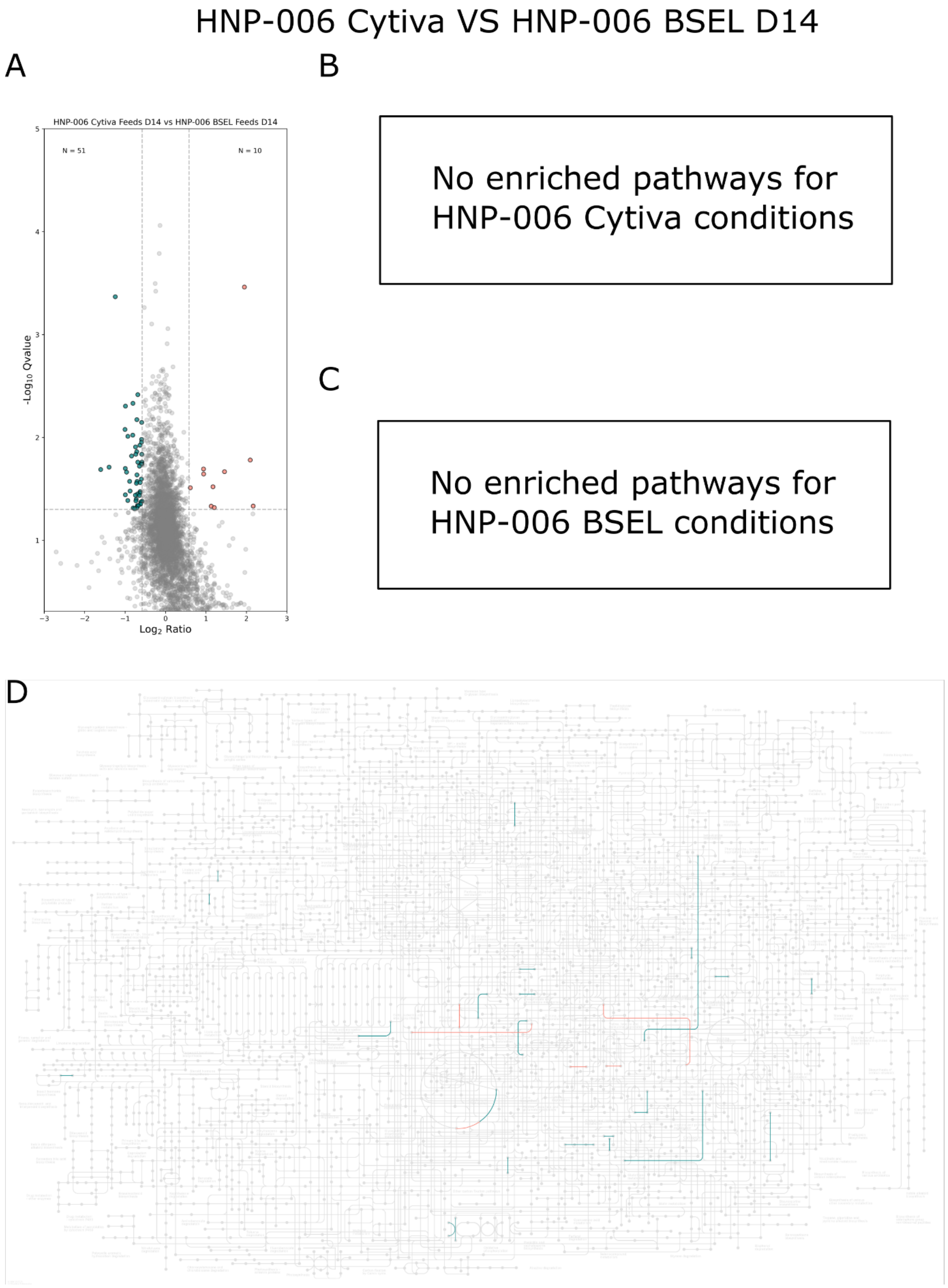


**Figure S25**: HNP-006 Cytiva (blue) vs HNP-006 BSEL (pink) D14 A) Volcano plot, B-C) gene set enrichment analysis D) KEGG mapped pathways

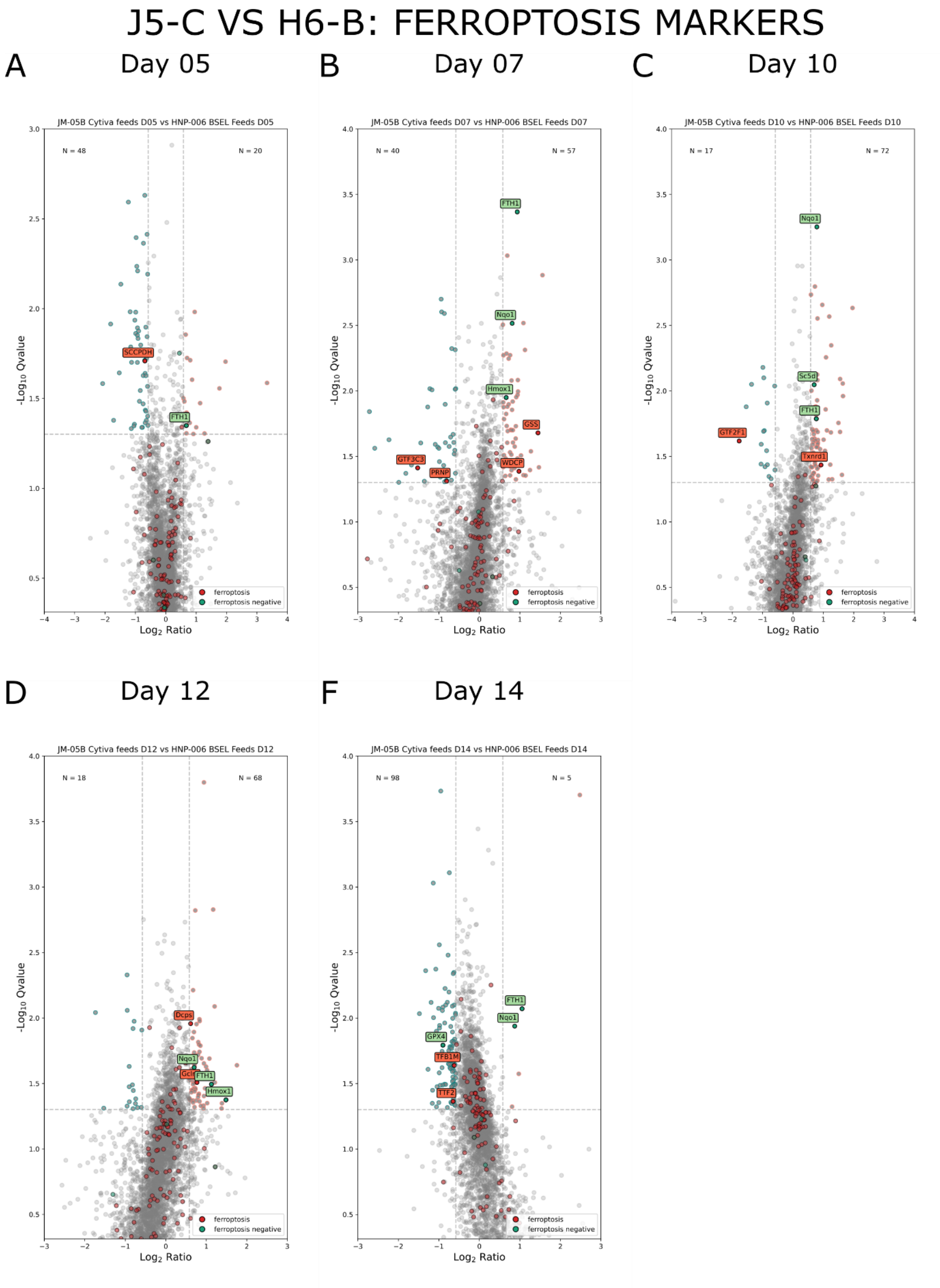


**Figure S26**: Differentially abundant proteins, with pro-ferroptotic (red) and anti-ferroptotic (green) markers labelled for J5-C (blue) vs H6-B (pink) for **A)** day 5 **B)** day 7 **C)** day 10 **D)** day 12 **E)** day 14

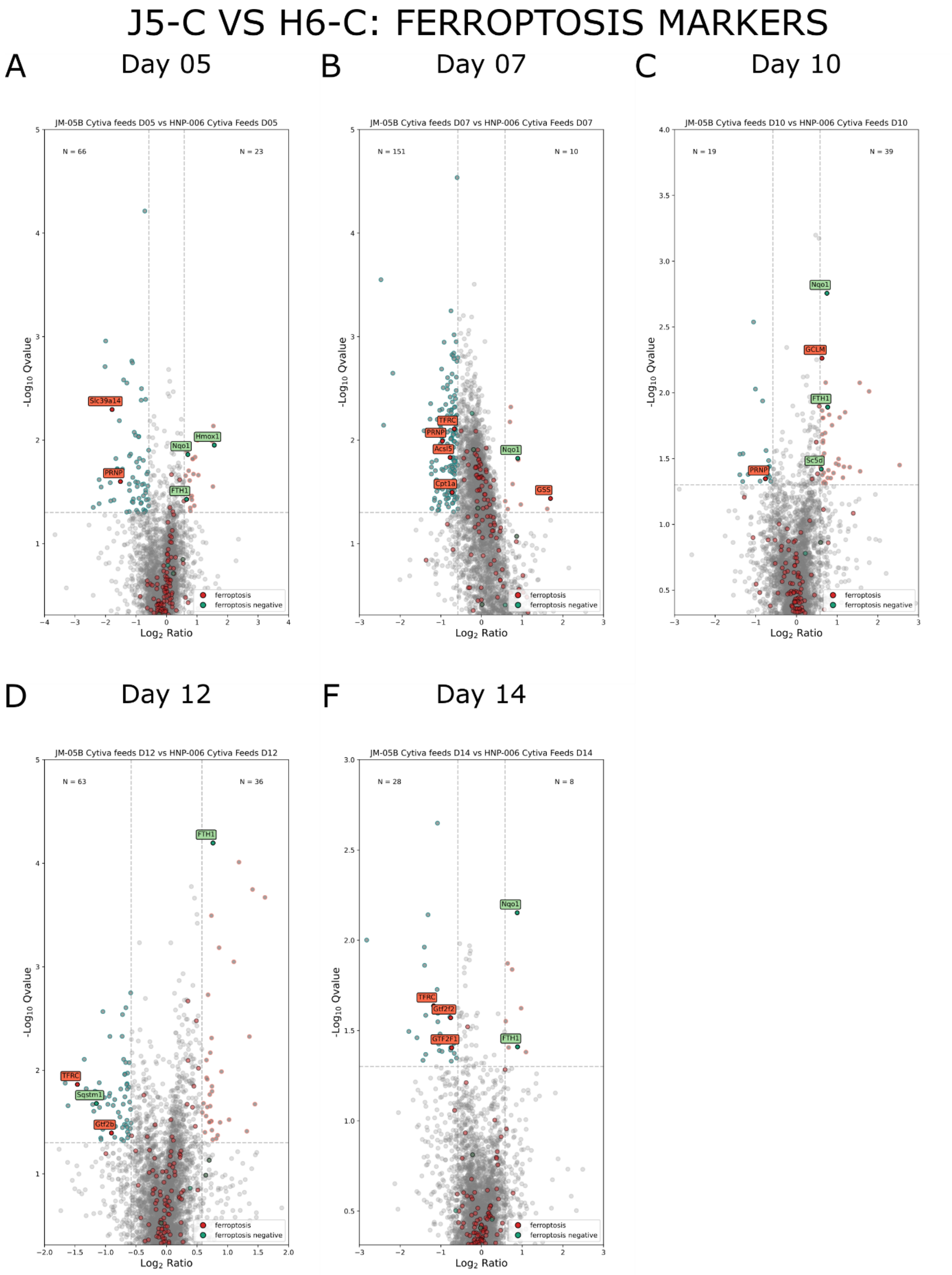


**Figure S27**: Differentially abundant proteins, with pro-ferroptotic (red) and anti-ferroptotic (green) markers labelled for J5-C (blue) vs H6-C (pink) for **A)** day 5 **B)** day 7 **C)** day 10 **D)** day 12 **E)** day 14

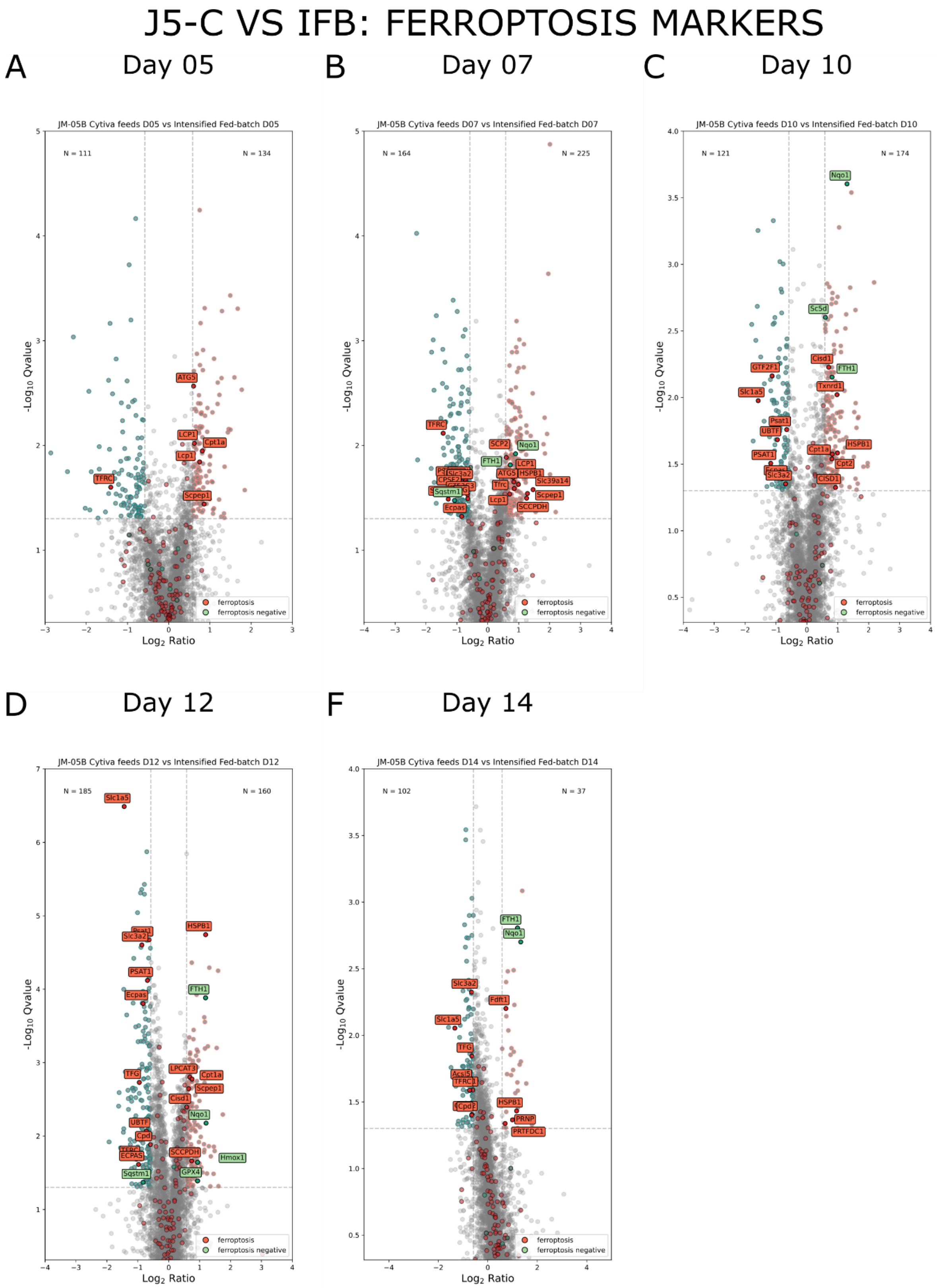


**Figure S28**: Differentially abundant proteins, with pro-ferroptotic (red) and anti-ferroptotic (green) markers labelled for J5-C (blue) vs IFB (pink) for **A)** day 5 **B)** day 7 **C)** day 10 **D)** day 12 **E)** day 14

### J5-C VS H6-B: FAO MARKERS

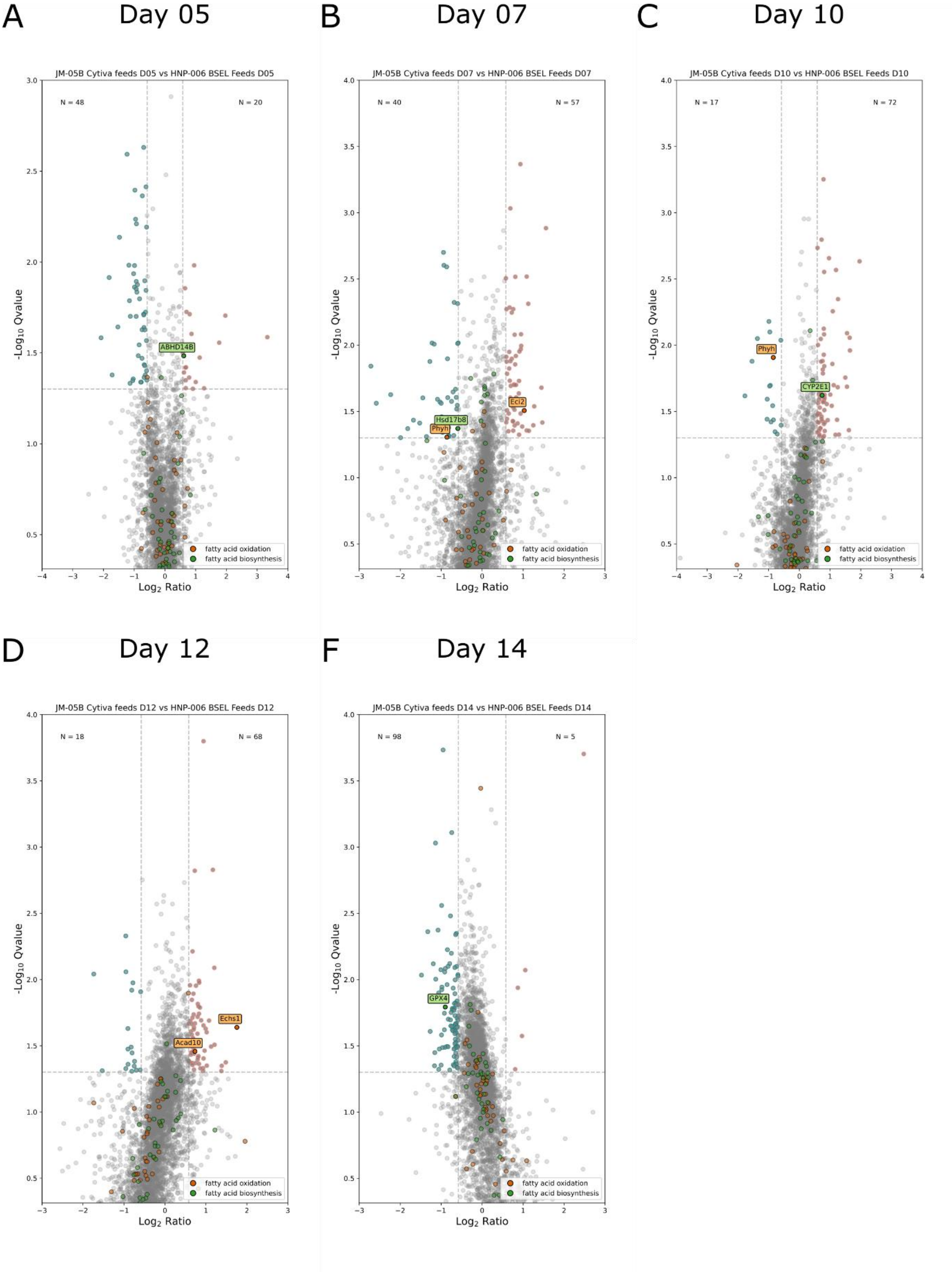


**Figure S29**: Differentially abundant proteins, with fatty acid oxidation (orange) and fatty acid biosynthesis (green) markers labelled for J5-C (blue) vs H6-B (pink) for **A)** day 5 **B)** day 7 **C)** day 10 **D)** day 12 **E)** day 14

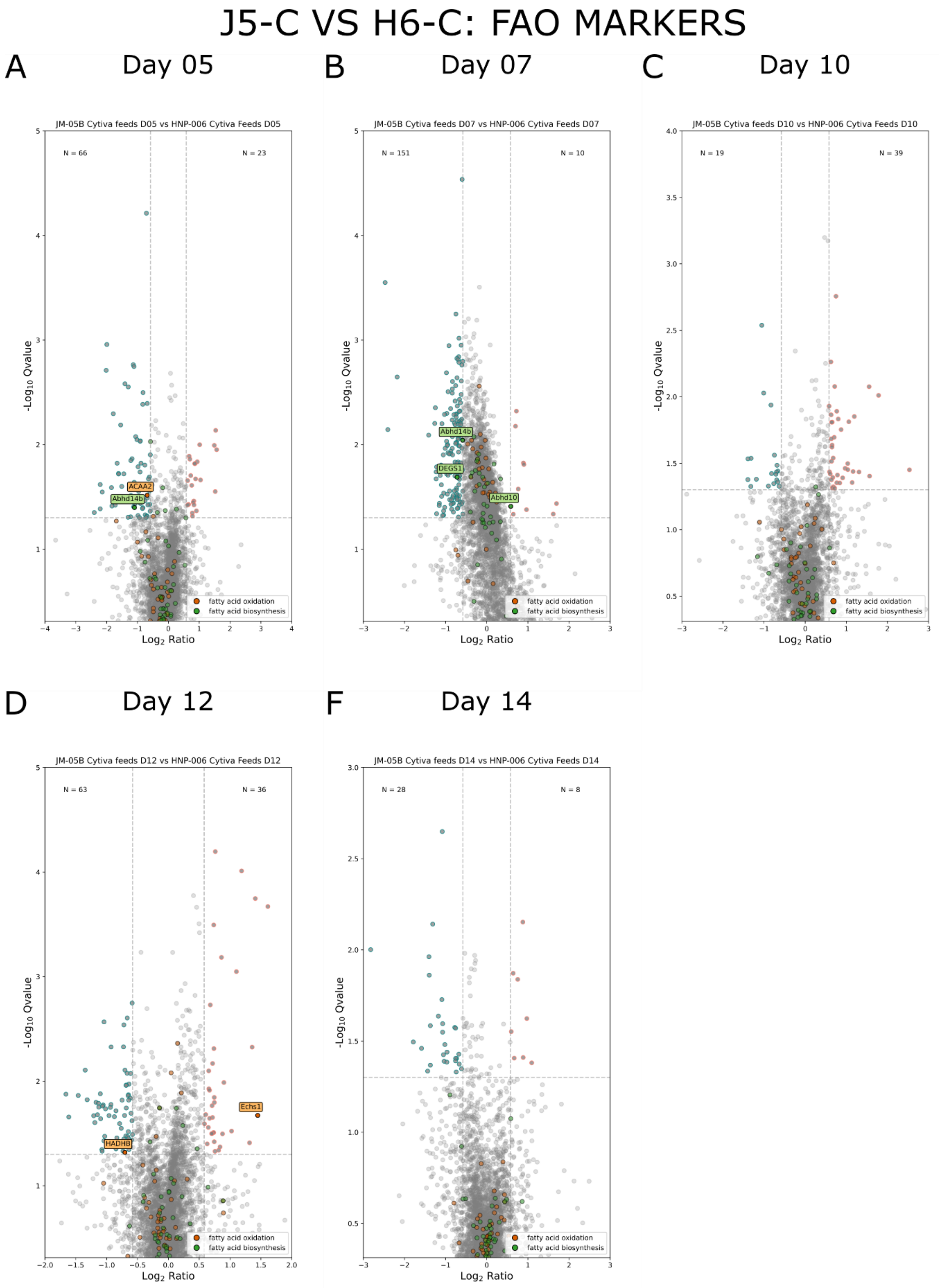


**Figure S30**: Differentially abundant proteins, with fatty acid oxidation (orange) and fatty acid biosynthesis (green) markers labelled for J5-C (blue) vs H6-C (pink) for **A)** day 5 **B)** day 7 **C)** day 10 **D)** day 12 **E)** day 14

### J5-C VS IFB: FAO MARKERS

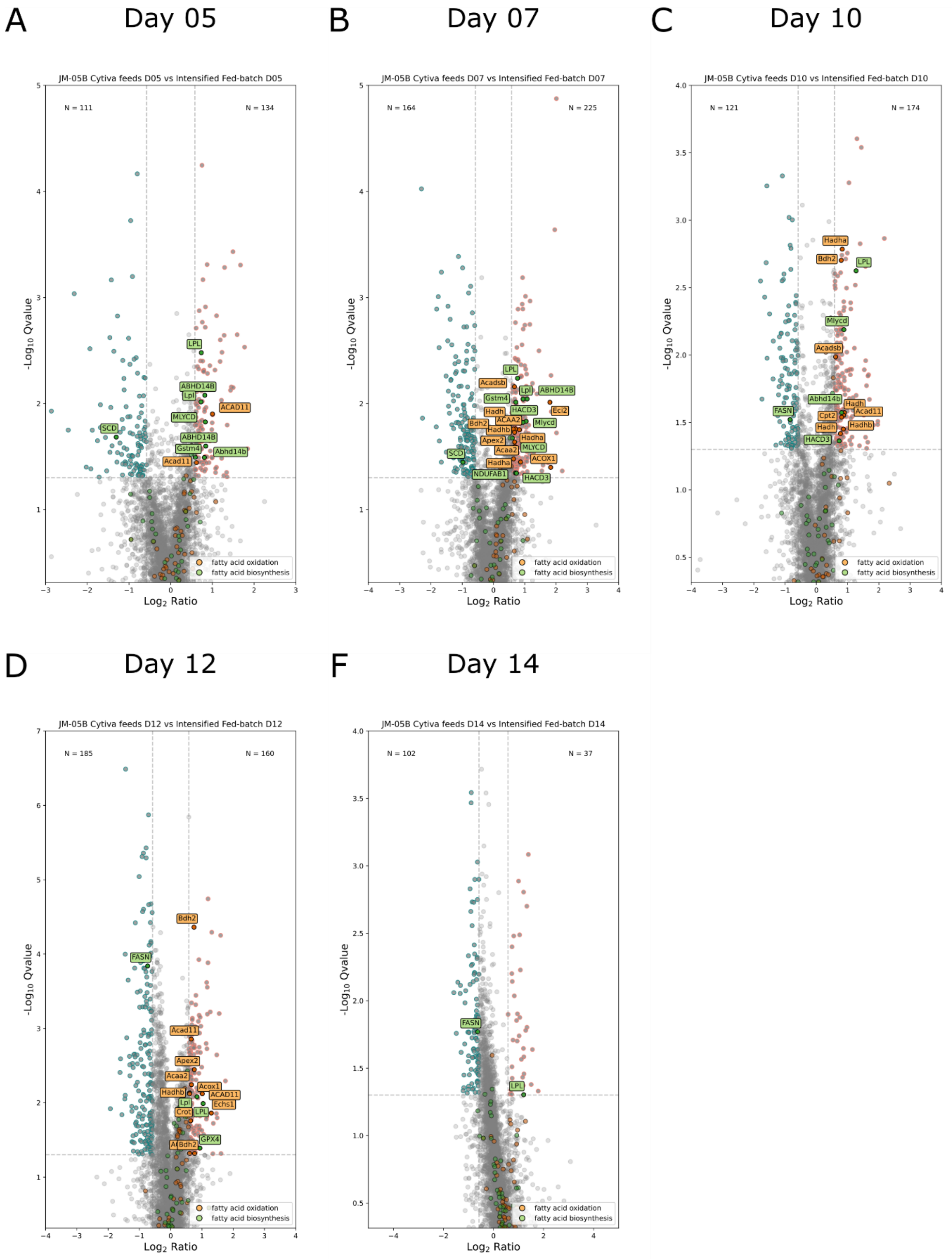


**Figure S31**: Differentially abundant proteins, with fatty acid oxidation (orange) and fatty acid biosynthesis (green) markers labelled for J5-C (blue) vs IFB (pink) for **A)** day 5 **B)** day 7 **C)** day 10 **D)** day 12 **E)** day 14

### J5-C VS H6-B: ROS MARKERS

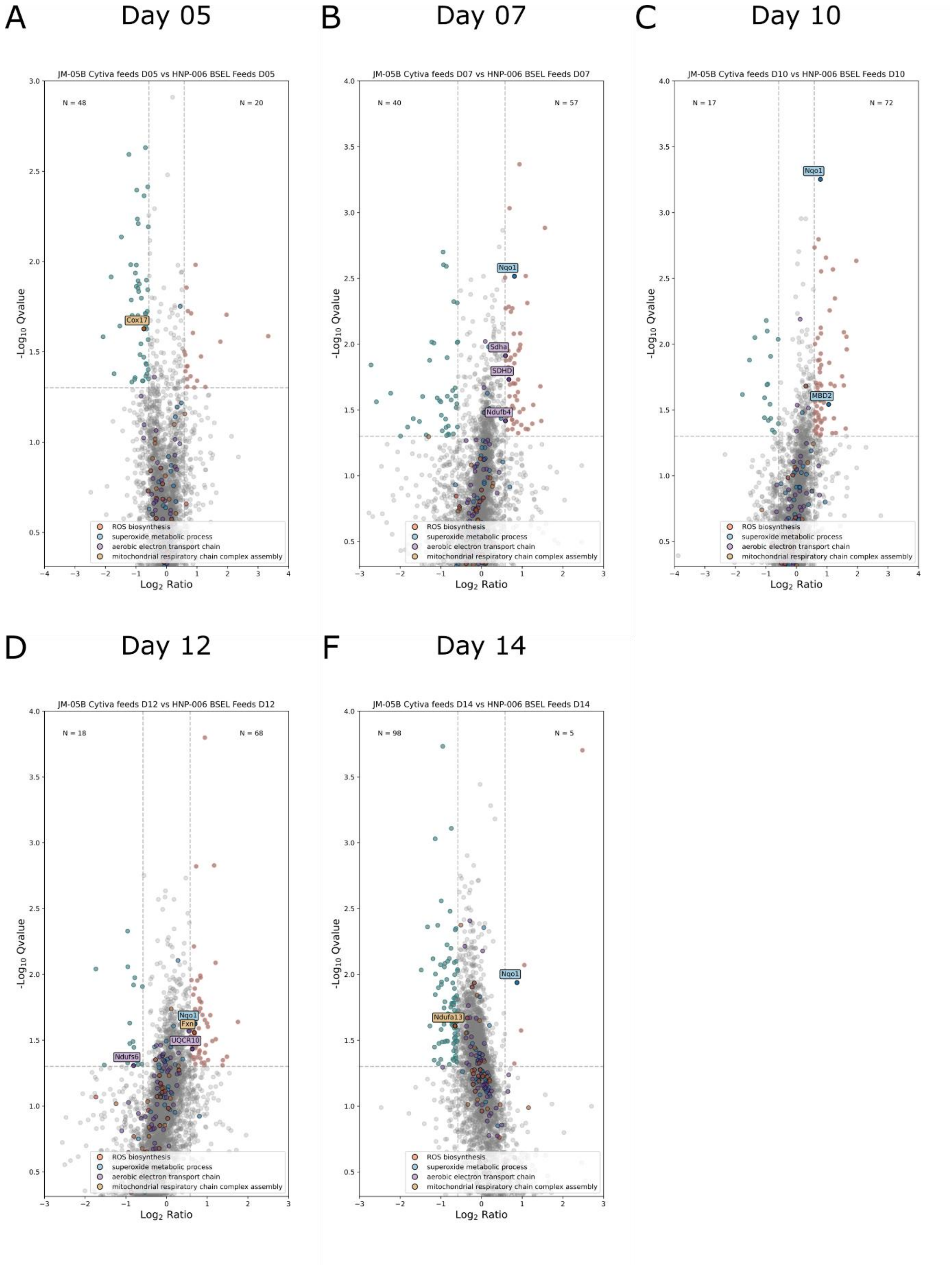


**Figure S32**: Differentially abundant proteins, with ROS biosynthesis (red), superoxide metabolic process (blue), aerobic transport chain (purple) oxidative and mitochondrial respiratory chain complex assembly (tan) markers labelled for J5-C (blue) vs H6-B (pink) for **A)** day 5 **B)** day 7 **C)** day 10 **D)** day 12 **E)** day 14

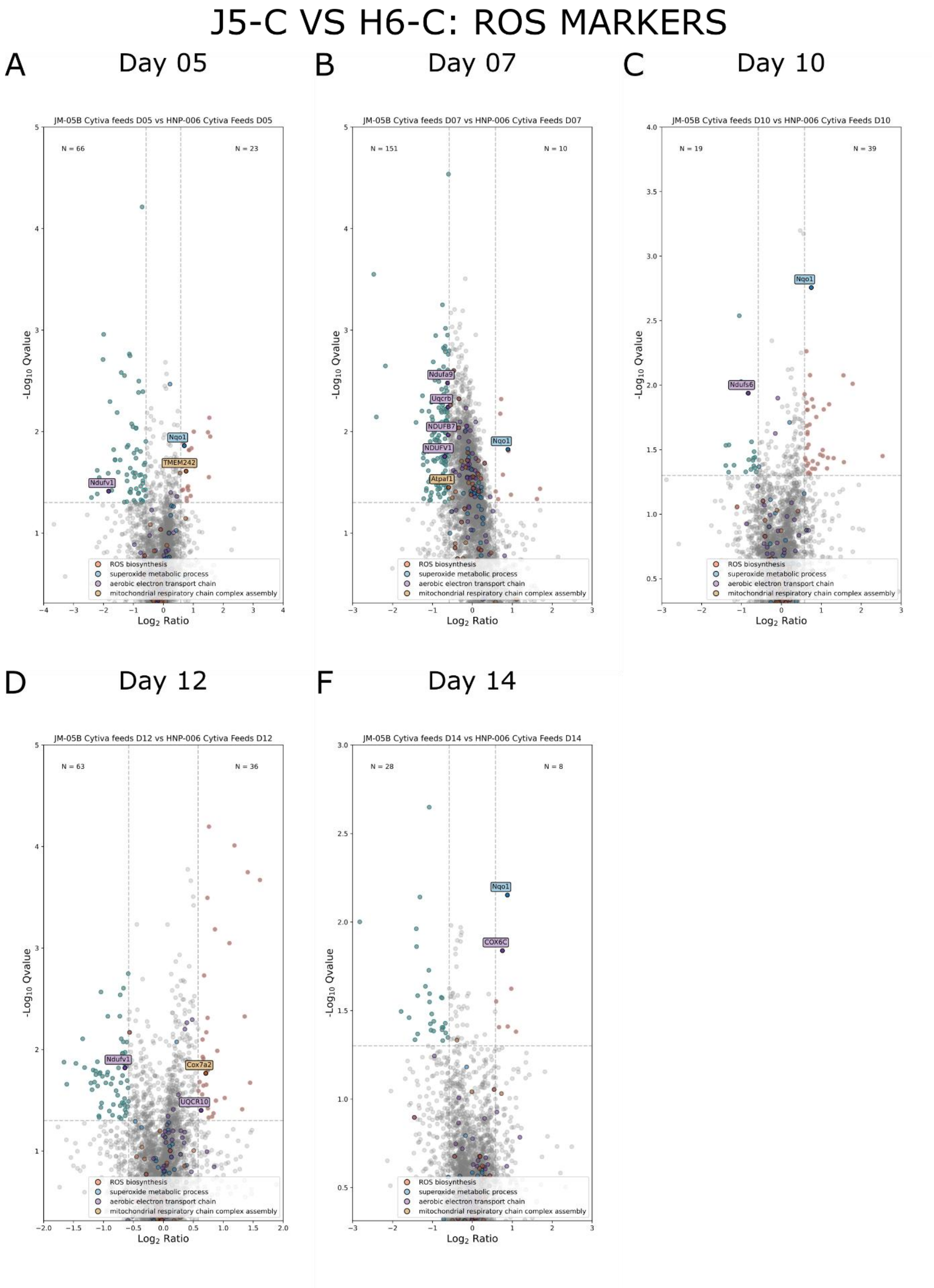


**Figure S33**: Differentially abundant proteins, with ROS biosynthesis (red), superoxide metabolic process (blue), aerobic transport chain (purple) oxidative and mitochondrial respiratory chain complex assembly (tan) markers labelled for J5-C (blue) vs H6-C (pink) for **A)** day 5 **B)** day 7 **C)** day 10 **D)** day 12 **E)** day 14

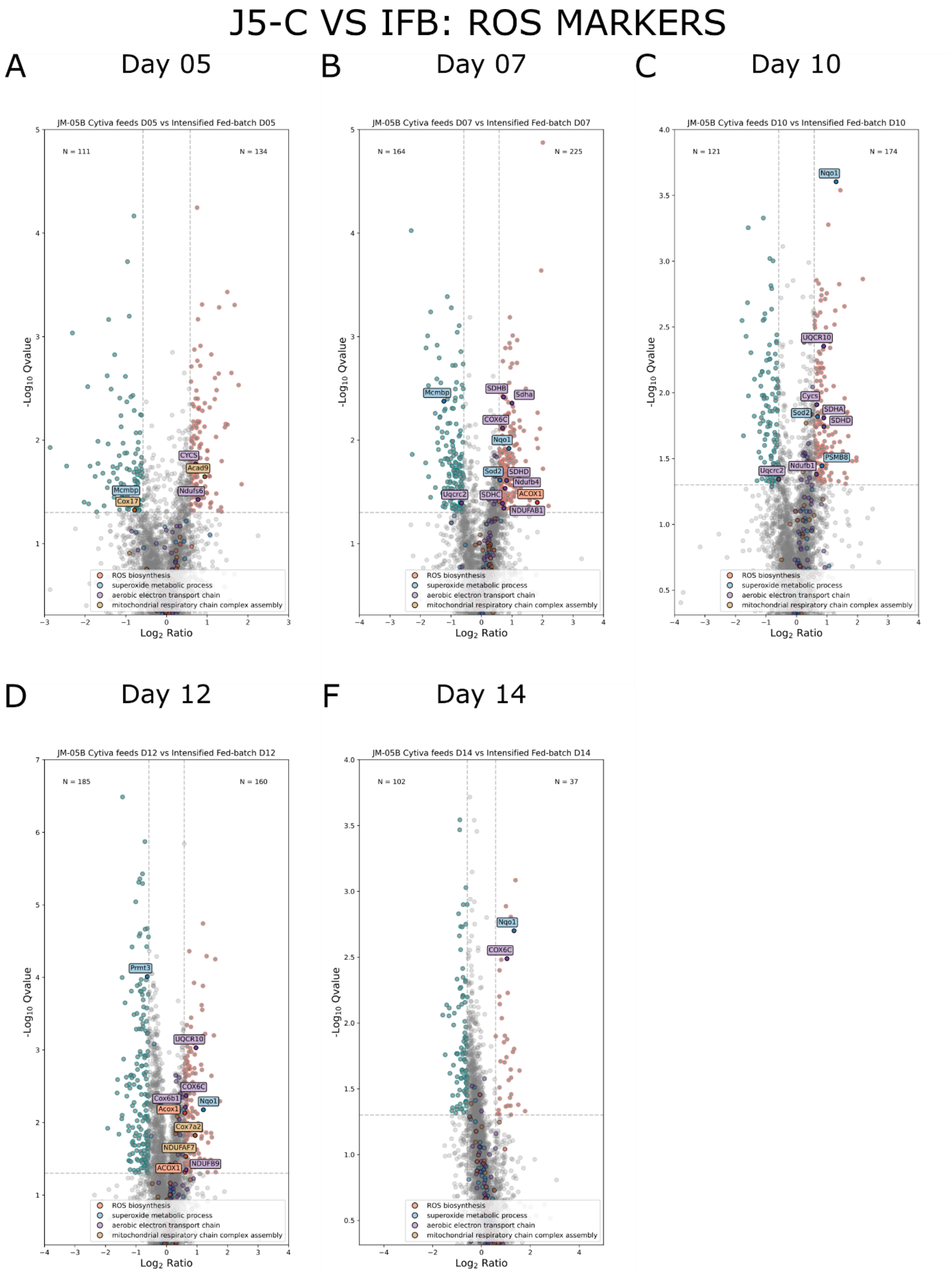


**Figure S34**: Differentially abundant proteins, with ROS biosynthesis (red), superoxide metabolic process (blue), aerobic transport chain (purple) oxidative and mitochondrial respiratory chain complex assembly (tan) markers labelled for J5-C (blue) vs IFB (pink) for **A)** day 5 **B)** day 7 **C)** day 10 **D)** day 12 **E)** day 14

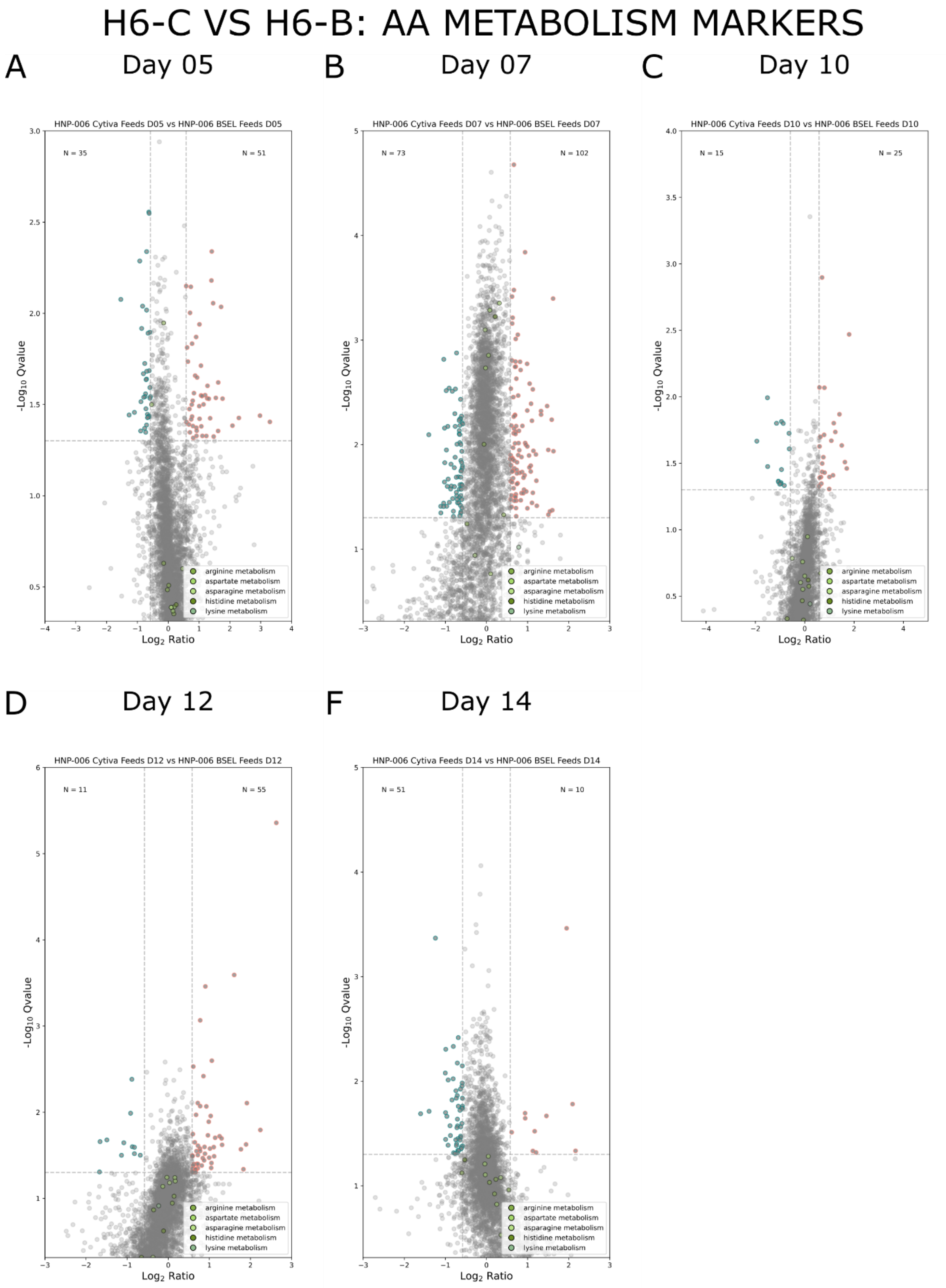


**Figure S 35:** Differentially abundant proteins, with markers for the metabolism of amino acids (arginine, aspartate, asparagine, histidine, and lysine)which are different in BSEL feed in green, with H6-C (blue) vs H6-B (pink) for **A)** day 5 **B)** day 7 **C)** day 10 **D)** day 12 **E)** day 14

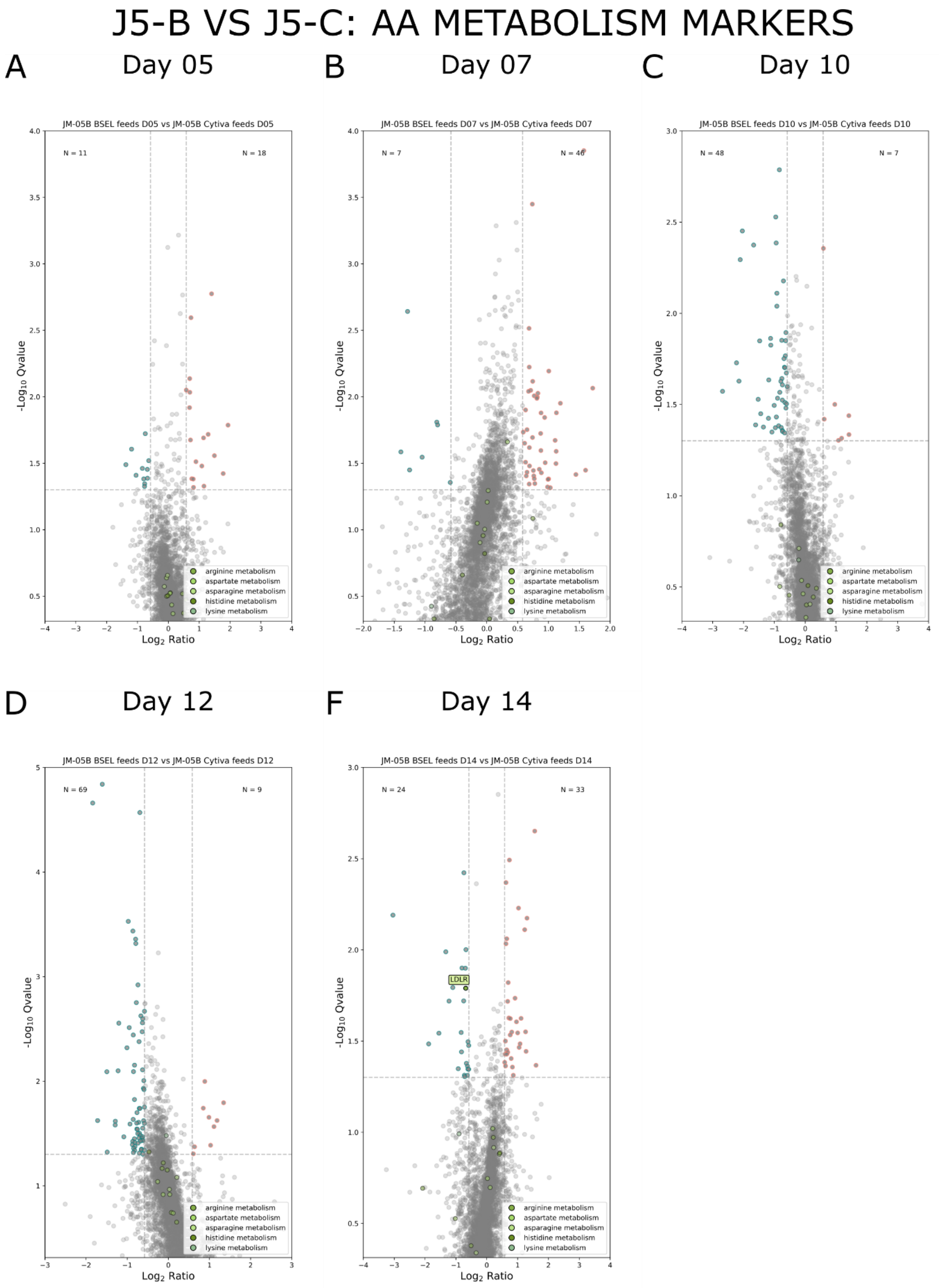


**Figure S36**: Differentially abundant proteins, with markers for the metabolism of amino acids (arginine, aspartate, asparagine, histidine, and lysine)which are different in BSEL feed in green, with J5-B (blue) vs  J5-C (pink) for **A)** day 5 **B)** day 7 **C)** day 10 **D)** day 12 **E)** day 14

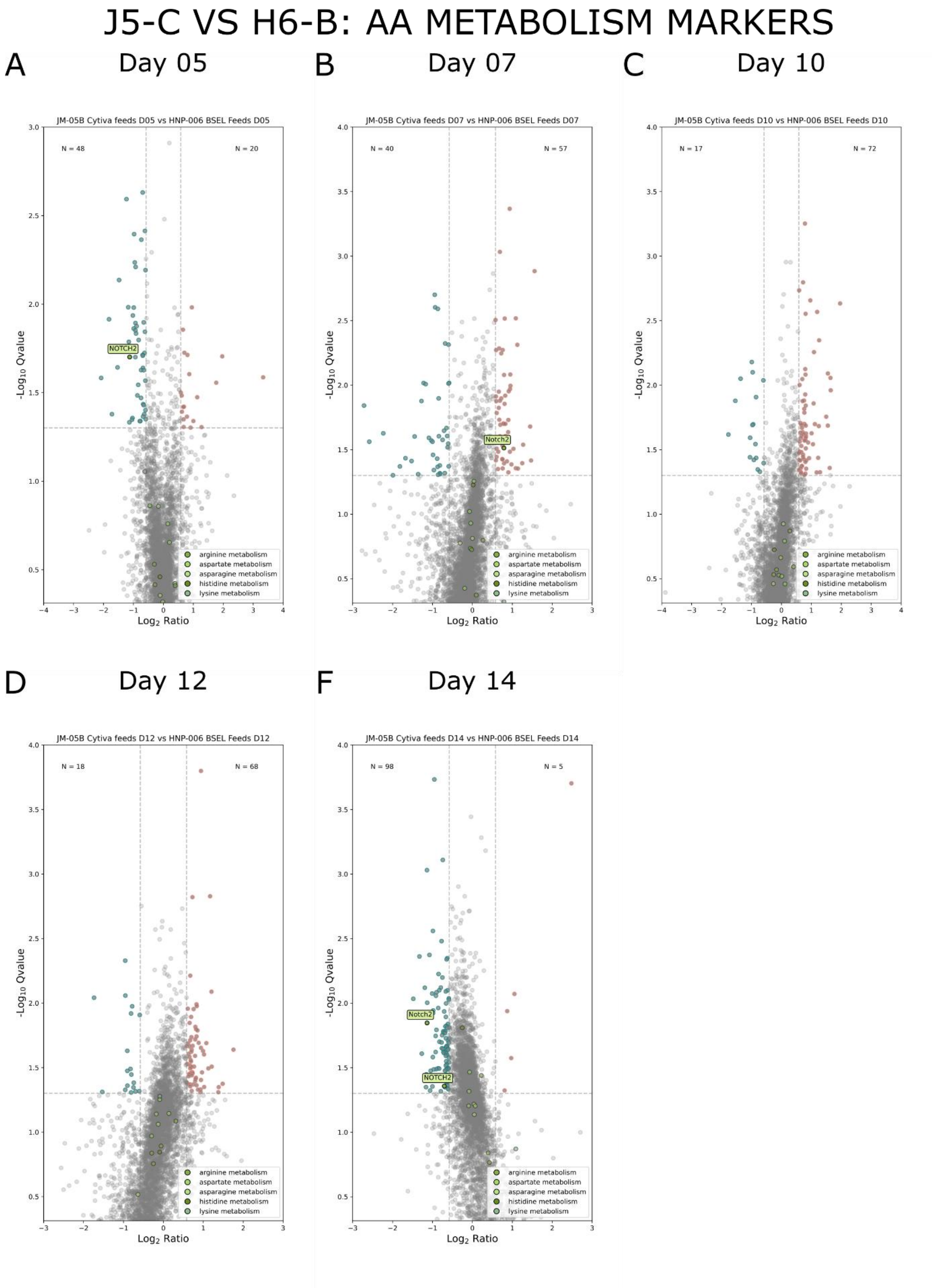


**Figure S37**: Differentially abundant proteins, with markers for the metabolism of amino acids (arginine, aspartate, asparagine, histidine, and lysine) which are different in BSEL feed in green, with J5-C (blue) vs H6-B (pink) for **A)** day 5 **B)** day 7 **C)** day 10 **D)** day 12 **E)** day 14
